# SIMVI reveals intrinsic and spatial-induced states in spatial omics data

**DOI:** 10.1101/2023.08.28.554970

**Authors:** Mingze Dong, David Su, Harriet Kluger, Rong Fan, Yuval Kluger

## Abstract

Spatial omics technologies enable the analysis of gene expression and interaction dynamics in relation to tissue structure and function. However, existing computational methods may not properly distinguish cellular intrinsic variability and intercellular interactions, and may thus fail to capture spatial regulations for further biological discoveries. Here, we present Spatial Interaction Modeling using Variational Inference (SIMVI), an annotation-free framework that disentangles cell intrinsic and spatial-induced latent variables for modeling gene expression in spatial omics data. We derive theoretical support for SIMVI in disentangling intrinsic and spatial-induced variations. By this disentanglement, SIMVI enables estimation of spatial effects (SE) at a single-cell resolution, and opens up various opportunities for novel downstream analyses. To demonstrate the potential of SIMVI, we applied SIMVI to spatial omics data from diverse platforms and tissues (MERFISH human cortex, Slide-seqv2 mouse hippocampus, Slide-tags human tonsil, spatial multiome human melanoma, cohort-level CosMx melanoma). In all tested datasets, SIMVI effectively disentangles variations and infers accurate spatial effects compared with alternative methods. Moreover, on these datasets, SIMVI uniquely uncovers complex spatial regulations and dynamics of biological significance. In the human tonsil data, SIMVI illuminates the cyclical spatial dynamics of germinal center B cells during maturation. Applying SIMVI to both RNA and ATAC modalities of the multiome melanoma data reveals potential tumor epigenetic reprogramming states. Application of SIMVI on our newly-collected cohort-level CosMx melanoma dataset uncovers space-and-outcome-dependent macrophage states and the underlying cellular communication machinery in the tumor microenvironments.

## Main

In recent years, spatial omics technologies have become a prominent tool to explore tissue organization and function at fine-grained resolutions. Emerging imaging-based spatial transcriptomics (ST) technologies, such as SeqFISH [1], MERFISH [2, 3], CosMx [4] are able to profile hundreds to thousands of genes at a single-cell or even subcellular resolution. Sequencing-based spatial omics technologies, such as DBiT-seq [5–7], Slide-seqV2 [8], HDST [9], Slide-tags [10] and Stereo-seq [11, 12], provide genome-wide gene profiling at a cellular or near cellular resolution. These technologies enable characterizations of the spatial microenvironment effect on gene expression at the cellular level.

Conceptually, a cell’s gene expression is determined both by its intrinsic properties and by its spatial microenvironment. Intrinsic properties should be understood as the part of gene expression that is mechanistically independent of neighborhood. They include but are not limited to cell subtypes or phases of cell cycle. Conversely, the spatial microenvironment involve spatially distributed signaling molecules and interacting cells [13]. Deconvolving the two factors is of fundamental importance in spatial omics data analysis, as it would unveil how one cell with a certain phenotype interacts with its local environment giving rise to gene expression changes. This is of particular importance in investigating complex biological processes involving both intrinsic state shift and spatial dynamics.

Yet, the task brings forth two significant challenges. First, cells of the same type often cluster together in space, creating spatial patterns of gene expressions that mostly reflect intrinsic properties rather than spatial interactions. Second, different cell types may have different responses to their local environment, particularly in the case of cellular interaction (Fig. 1a). To the best of our knowledge, most existing computational approaches do not address the challenge of disentangling intrinsic and spatial patterns from spatial omics data. Most existing analysis methods for spatial omics data either aim to identify spatially variable genes [14–19], learn spatial-aware embeddings [20–26], select genes that mediate cell interactions [27], predict patient phenotypes [28, 29], or learn gene expression changes associated with local interactions [30–32]. These methods do not model intrinsic variations thus implicitly improperly assume that the gene expression (with or without conditioning on other covariates), is solely determined by its spatial environments. While several works made preliminary efforts [21, 30, 32] to account for cellular intrinsic variability, a failure of disentanglement has been noted due to the lack of identifiability [30].

**Fig. 1.**
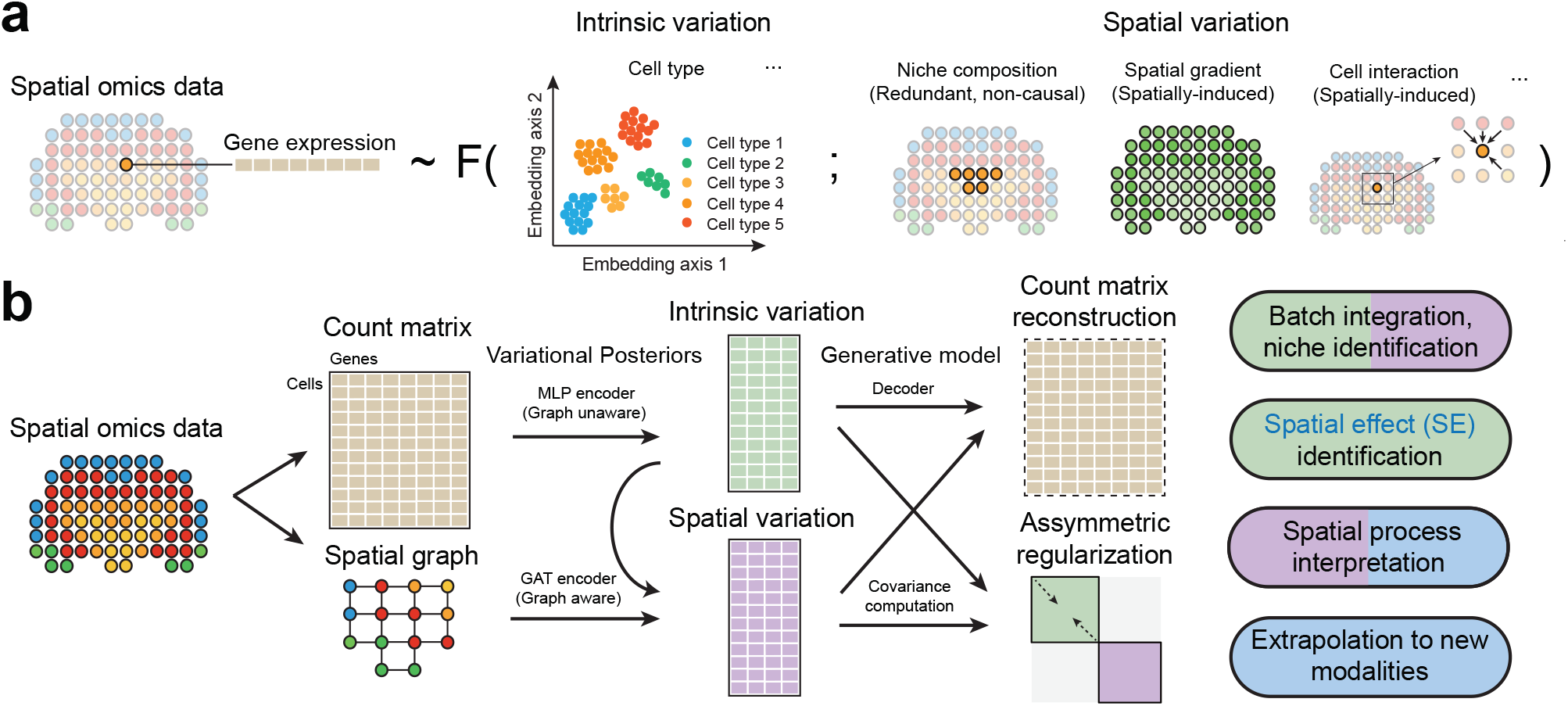
Overview of the SIMVI framework. **a**. Gene expression in spatial omics data is assumed to be a (noisy) function of intrinsic factors (e.g., cell type) and spatial factors (e.g., niche composition, spatial gradient, cellular interaction). **b**. SIMVI workflow. Spatial omics data are transformed into graph-structured format. Intrinsic and spatial variations are estimated by multilayer perceptron (MLP) and graph attention network (GAT) variational posteriors. The loss function consists of the evidence lower bound (ELBO) and additional regularization terms.

In this work, we present Spatial Interaction Modeling using Variational Inference (SIMVI), a novel variational inference framework designed to disentangle intrinsic and spatial-induced variations in spatial omics data. We provide rigorous theoretical support for the identifiability of the SIMVI model in achieving this disentanglement. The learned variations of SIMVI enable various downstream analyses, including but not limited to clustering, visualization and differential expression analysis. Furthermore, the disentangled representations enable the estimation of spatial effect (SE), i.e., the effect of spatial location on gene expression, at a single-cell-gene level. The spatial effect estimation procedure also returns a variance decomposition for each gene, which can return rankings of genes associated with spatial regulation. Finally, the SIMVI SE estimation method offers the versatility to adapt to new modalities in spatial multi-omics data.

We comprehensively benchmarked SIMVI in its ability to reveal disentangled variations and spatial effects. We applied SIMVI to a number of public datasets across different platforms and tissues, including MERFISH human cortex [3], Slide-seqv2 mouse hippocampus [8], Slide-tags human tonsil, and spatial multiome human melanoma [10]. Across all datasets, SIMVI demonstrates superior ability to disentangle intrinsic and spatial variations and infer spatial effects compared to alternative methods. Furthermore, SIMVI uniquely reveals complex spatial interactions and dynamics, providing novel biological insights. In the Slide-tags tonsil data, SIMVI illuminates spatial-dependent dynamics of germinal center B cells during maturation. In the multiome melanoma data, we employed SIMVI to infer refined spatial niches and multi-omics spatial effects, discovering potential epigenetic reprogramming states. Applying SIMVI to our newly collected cohort-level CosMx melanoma dataset, we identified macrophage states that exhibit divergent spatial patterns and are associated with varied patient outcomes. In the dataset, SIMVI further uncovers the underlying cellular interaction machinery within tumor microenvironments, characterized by asymmetric dependencies between the ligand-receptor strength and gene spatial effects.

## Results

### Disentanglement of intrinsic and spatial-induced variations by SIMVI

In our work, we model the gene expression *x*^*i*^ of each cell *i* by two sets of low-dimensional latent variables *z*^*i*^, *s*^*i*^ characterizing the intrinsic and spatial variation respectively (Methods, Supplementary Note 1). The SIMVI framework infers the two sets of latent variables from spatial omics data using variational autoencoders (VAE) [33]. Specifically, SIMVI encodes intrinsic latent variables *z*^*i*^ by variational posteriors of the gene expression *x*^*i*^ of cell *i*, and the spatial latent variables *s*^*i*^ by graph neural network (GNN) variational posteriors, as functions of neighborhood gene expression *x*^*N*(*i*)^ (see Methods). Parameters of the variational posteriors are optimized by maximizing the evidence lower bound (ELBO, see Methods).

The theoretical possibility of successful disentanglement hinges on model identifiability. In our context, model identifiability means that the intrinsic and spatial variations must be uniquely disentangled from observed gene expressions. Without this property, the inferred intrinsic and spatial latent variables could become arbitrary mixtures of the true underlying variations.

In our work, we derived novel theoretical support showing that, under suitable assumptions, the model identifiability can be achieved for disentangling intrinsic and spatial variations. Specifically, it is possible to find a representation of the intrinsic latent variables up to a non-linear transformation. As for the spatial latent variables, it is possible to infer them up to an invertible linear transformation (Supplementary Note 1).

The SIMVI model incorporates essential elements for establishing theoretical identifiability (Supplementary Note 1). They include a shared parameter design for the variational posteriors, and a novel asymmetric regularization term minimizing information in the intrinsic variation (Fig. 1b, Methods). SIMVI further provides an option to randomly permute a subset of genes in the input data as a pretraining step, which was empirically shown to enhance disentanglement performance (see Methods, Supplementary Note 3). After optimization, SIMVI returns two disentangled representations from spatial omics data that capture intrinsic and spatial variations. The disentanglement enhances our capacity to explore and characterize cellular diversity at different levels including cell-intrinsic properties, spatial heterogeneity, and their combination. These representations facilitate a variety of analyses, including clustering, differential gene expression analysis, and visualization. Finally, SIMVI can also incorporate covariates such as batch labels by the conditional design implemented in scVI [34], enabling analyses for datasets comprised of multiple batches.

The disentangled representations provide a natural framework for estimating the effect of spatial location on individual gene expression, by viewing spatial variation as “treatments” and intrinsic variation as “covariates”. This formulation transforms the task into a problem of estimating continuous treatment effects, a well-established paradigm in causal inference. In this study, we employ double machine learning (DML [35]), a method from causal inference that allows estimating the conditional spatial effect on individual cells (see Methods and Supplementary Note 2).

In some cases, discerning the spatial effect may be impossible. For instance, if a particular cell type exhibits unique intrinsic states and localizes to spatial niches that do not overlap with those of other cells, then it is impossible to determine whether its expression shifts are influenced by differences in intrinsic states or by spatial niches. Such an issue is well-known as the violation of the positivity assumption in causal inference ([36, 37], Supplementary Note 2). To this end, we derived single-cell positivity indices that assesses potential positivity violations for each cell. The indices can be used to exclude cells that violate the positivity assumption, enforcing reliable spatial effect inference (Methods, Supplementary Note 2).

### SIMVI effectively disentangles variations

To demonstrate SIMVI’s ability to reveal disentangled variations, we applied SIMVI to two datasets profiled via representative imaging-based and sequencing-based technologies respectively. The first dataset is from a recent MERFISH study, where the researchers profiled 4,000 genes in human cortex [3], including both middle temporal gyrus (MTG) and superior temporal gyrus (STG). As the dataset includes detailed annotations and experimental replicates, it provides an ideal case for benchmarking SIMVI in disentangling intrinsic and spatial variations while removing batch effects. The second dataset is the widely used Slide-seqV2 mouse hippocampus data [8], that offers unbiased spatial profiling of gene expression in the mouse hippocampus at near-cellular resolution.

An overview of the MERFISH MTG/STG dataset is shown in Fig. 2a and Supplementary Fig. 5a. The dataset exhibits spatial structure of cortex layers that do not perfectly overlap between replicates. Apart from the cortex layer structure, another prominent spatial pattern within the data is the “local niche” vascular structure, characterized by the colocalization of mural and endothelial cells expressing *MYH11* [3].

**Fig. 2.**
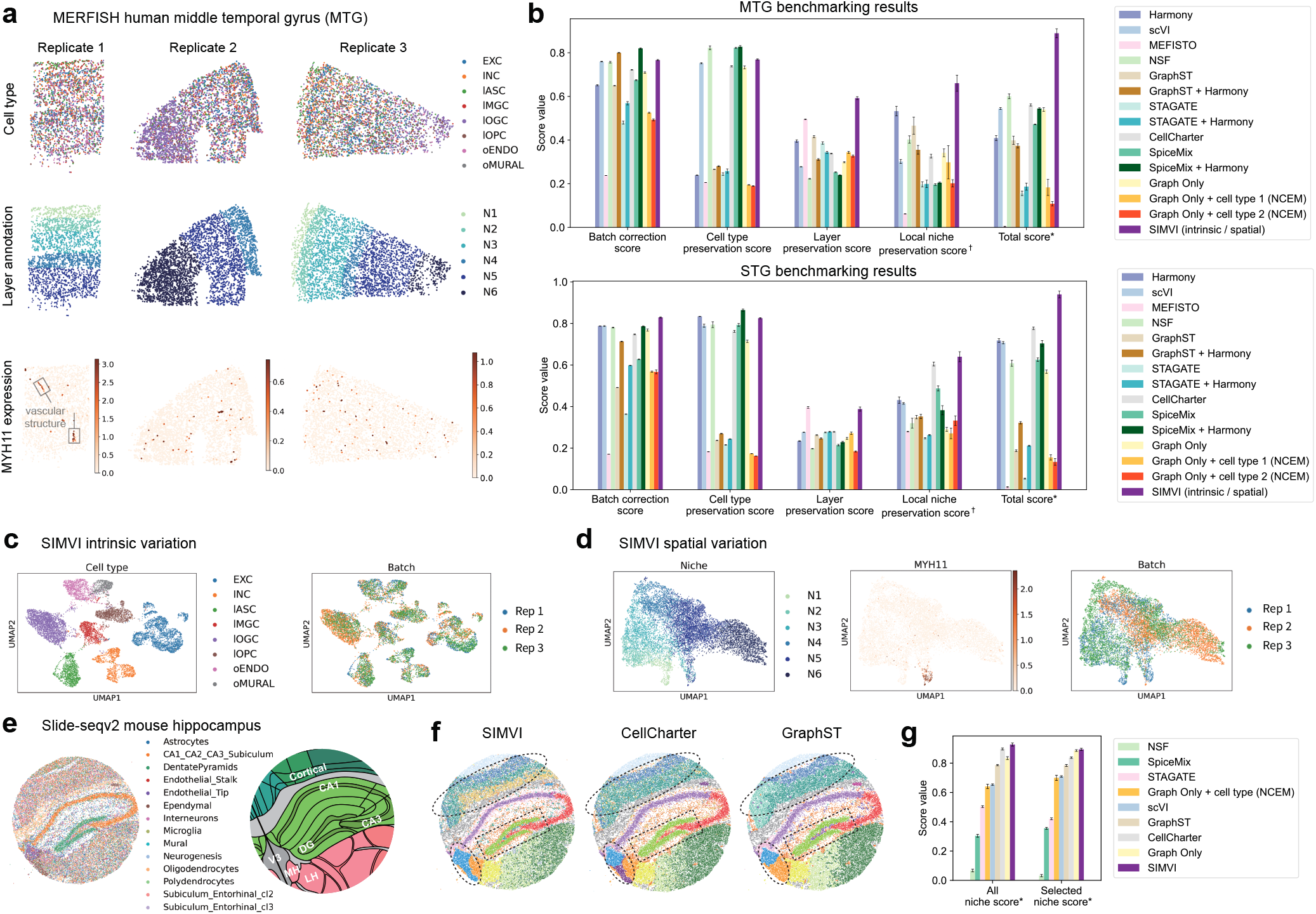
SIMVI reveals intrinsic and spatial variations in MERFISH human cortex and Slide-seqV2 mouse hippocampus. **a**. Overview of the MERFISH middle temporal gyrus (MTG) data, showing spatial organizations of cell type, layer annotation, and log normalized *MYH11* expression for MTG replicates 1-3. **b**. Bar plots showing metric scores for the MERFISH MTG and STG datasets respectively (See Methods for definitions of the scores). The bar heights represent the average performance across 10 different random seeds, with the error bars showing standard errors. **c**. UMAP visualization of the SIMVI intrinsic variation, colored by cell types and replicate labels. **d**. UMAP visualization of the SIMVI spatial variation, colored by layer annotation, log normalized *MYH11* expression, and replicate labels. **e**. Overview of the Slide-seqv2 data showing spatial organization of pixels (left) and Allen Mouse Brain Atlas annotation for the region (right). CA: Cornu Ammonis, DG: Dentate Gyrus, V3: Third Ventricle, MH: Medial Habenula, LH: Lateral Hypothalamus. **f**. Spatial visualization of clusters obtained from SIMVI, CellCharter and GraphST. **g**. Bar plot showing the metric scores for the Slide-seqV2 mouse hippocampus data (See Methods for definitions of the scores). The bar heights represent the average performance across 10 different random seeds (*n* = 9 for SIMVI), with the error bars showing standard errors. *†* Each individual metric from all experiments is scaled to the range [0,1], then the final score is calculated as the average of these rescaled values. *: The score is obtained by first averaging the individual score values for each experiment, then rescaling the average score across all experiments to the range [0,1].

To evaluate the disentanglement performance, we defined four key metrics: batch correction, cell type preservation, layer preservation, and local niche preservation scores (See Methods). These metrics assess the effectiveness of embeddings in representing corresponding intrinsic or spatial information. Prior to benchmarking SIMVI against other methods, we conducted an extensive evaluation of the impact of parameter settings and model modifications on SIMVI’s performance, as detailed in Supplementary Figs. 1-4 and Supplementary Note 3.

Apart from SIMVI, we applied a variety of methods that learn batch integrated embeddings (Harmony, scVI [34, 38]), spatial-aware embeddings (MEFISTO, NSF, SpiceMix, GraphST, STAGATE [20–24]) or perform both simultaneously (GraphST + Harmony, STAGATE + Harmony, CellCharter, SpiceMix + Harmony [22–24, 26, 38]). We further tested baseline models (Graph Only, Graph Only + cell type 1, Graph Only + cell type 2) that have a SIMVI graph encoder design, yet replace intrinsic variation with different settings of covariates (batch label, batch label + cell type label, batch label + cell subtype label, see Methods). Such a design coincides with NCEM [30] when the covariate contains cell types. Overall, SIMVI consistently ranks as a top performer across all tasks, particularly excelling in spatially relevant tasks, achieving a notably higher total score in both MTG and STG (Fig. 2b, Supplementary Fig. 7e,f). Apart from SIMVI, we observed a clear distinction in different methods’ capability. Several methods that do not address disentanglement (scVI, NSF, CellCharter, SpiceMix, Graph Only) also excel at batch correction and cell type preservation. This is expected since cell type represents the most prominent variation in the data. However, these methods underperform in spatially relevant tasks. For these tasks, MEFISTO works reasonably well only in layer preservation, while there is no consistent winner apart from SIMVI in local niche preservation. This further underscores the essence of disentanglement in capturing all layers of information for spatial data analysis. Qualitatively, SIMVI intrinsic variation preserves cell type structures and remove batch effects. SIMVI spatial variation distinguishes different layers, including those not observed in all batches, and localizes cells with high expression levels of *MYH11* (Fig. 2c-d, Supplementary Fig. 5b-f).

We next applied SIMVI to the Slide-seqV2 mouse hippocampus data [8] (Fig. 2e). In this dataset, each pixel represents a mixture of cells, and the cell type annotation for each pixel is based on the dominant cell type obtained from deconvolution (Supplementary Fig. 8a-b). In this case, although the cell type label is no longer appropriate for benchmarking method performance, an evaluation on the spatial representation can still be conducted by leveraging an anatomic annotation for the same brain region (Allen Mouse Brain Atlas, Fig. 2e). Compared to prior methods with good performance (CellCharter, GraphST), SIMVI is the only method that correctly reveals the organizations of cortical layers and the Third Ventricle (V3) region (Fig. 2f). The superior performance of SIMVI is further quantitatively confirmed by our derived niche scores, where SIMVI has an advantage in either revealing all niches or selected niches that show clear spatial structures aligning with the anatomy (See Methods, Fig. 2g). The failure of alternative methods in identifying V3 is likely due to the different cell compositions in V3 upper and lower regions, which further highlights the need of disentangling intrinsic from spatial variations. Interestingly, we observed that the SIMVI Cornu Ammonis (CA) and Dentate Gyrus (DG) clusters show a consistent boundary with the manually derived annotation (See Methods), while CellCharter and GraphST identified thicker CA and DG regions (Fig. 3f, Supplementary Fig. 8e). This suggests a potential “oversmoothing” phenomenon for these methods through spatial neighborhood aggregation. A benchmark of the relative distances between externally annotated CA/DG regions and their spatial neighborhoods confirms our observation, and highlights SIMVI’s advantage in revealing true spatial structures (Supplementary Fig. 8f).

**Fig. 3.**
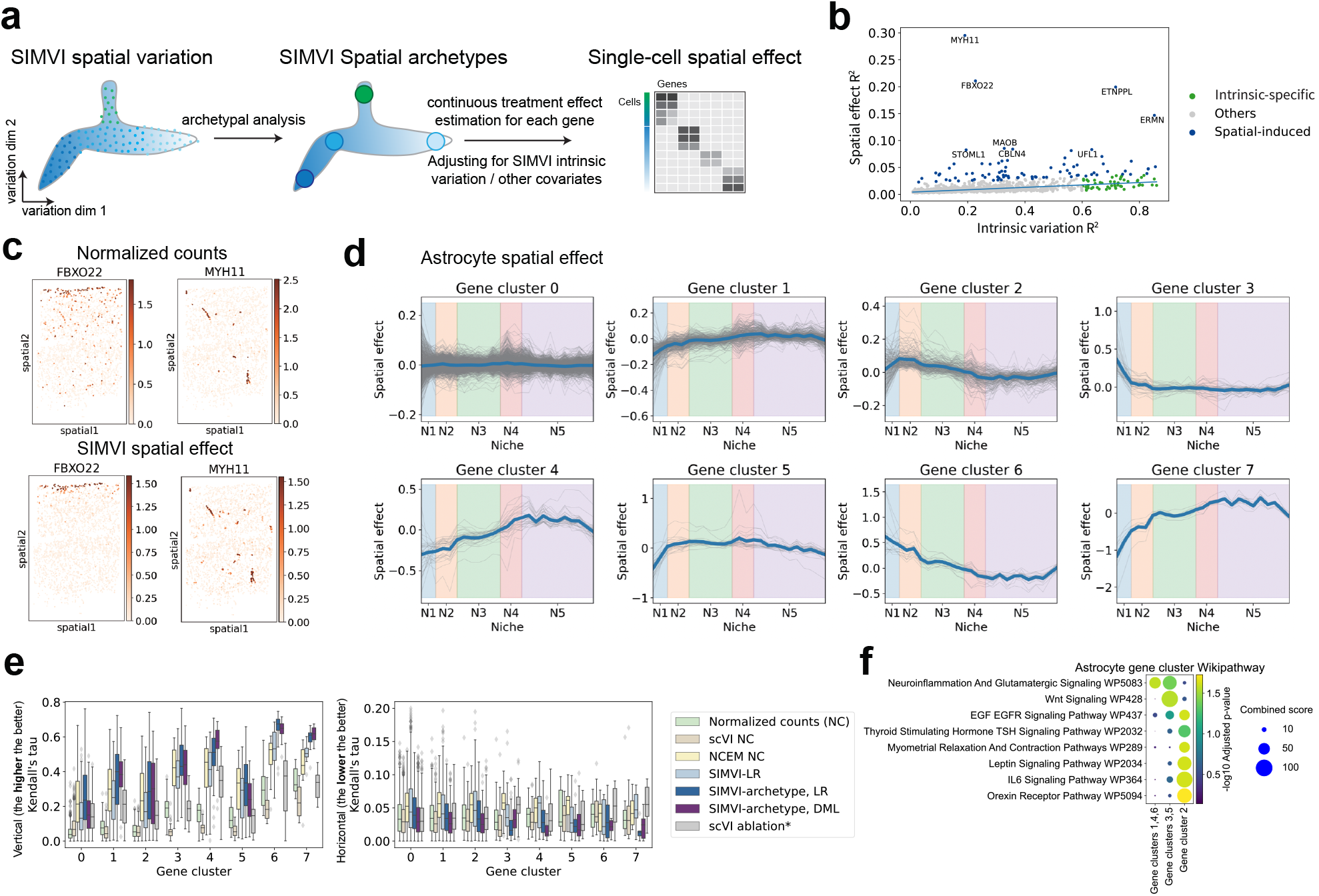
SIMVI infers single-cell level spatial effects. **a**. Overview of the SIMVI spatial effect estimation procedure. **b**. Scatter plot showing the intrinsic variation *R*^2^ and spatial effect *R*^2^ for each individual gene in MERFISH MTG data. Genes with scaled Huber regression residual larger than 10 were annotated as spatial-induced. Other genes with intrinsic-specific *R*^2^ larger than 0.6 were annotated as intrinsic-specific. **c**. Spatial visualizations of log normalized expression for representative genes (upper) and their spatial effects (lower). **d**. Visualizations of binned gene spatial effects grouped by clusters in astrocytes from MTG replicate 1 along data vertical axis. **e**. Boxplot showing Kendall’s tau on each individual gene of astrocytes from MTG replicate 1 and the spatial vertical coordinate (left) and horizontal coordinate (right). *: scVI ablation indicates the DML approach fixing SIMVI spatial variation and substituting SIMVI intrinsic variation with the scVI embedding. The gene number (n) is 1002, 265, 176, 40, 31, 23, 16, 8 for each cluster. **f**. Dotplot showing GSEA Wikipathway enrichment for genes from different spatial patterns.

### SIMVI infers accurate spatial effects

To estimate the spatial effect, we first decompose the spatial variation using archetypal analysis [39]. This method represents the spatial variation as a product of extreme spatial states (archetypes) and “archetype weight vectors”, which quantify the contribution of each archetype to individual cells. Next, we formulate a conditional treatment estimation task by considering the archetype weights as continuous treatments, and adjusting for confounders including intrinsic variation and other optional covariates (Fig. 3a). Specifically, we adopt the double machine learning procedure [35] to estimate the conditional spatial effect for each cell and each gene. The procedure also returns a decomposition of *R*^2^ (coefficient of determination) of gene expressions into the “intrinsic variation explained *R*^2^” and “Spatial effect explained *R*^2^”. Details of the procedure are described in Methods and Supplementary Note 2. Applying our procedure to MERFISH MTG data revealed meaningful archetypes corresponding to cortex layers and vascular structures (Supplementary Fig. 9). Additionally, we identified genes with distinct spatial patterns that showed notably high (outlier) spatial effect *R*^2^ values compared to the bulk (Fig. 3b, Supplementary Figs. 10–11). The spatial effects of these genes show cleaner spatial patterns in visualizations compared with normalized original counts (Fig. 3c, Supplementary Fig. 11).

We further focused on the spatial effect within a specific cell type (astrocyte, lASC, see Methods). We chose astrocytes because they are known to exhibit spatial diversity across cortex layers yet do not have a definitive subtype distinction (morphology, neurotransmitter type, etc) as those seen in neurons [40, 41]. Unsupervised analyses of spatial effects identified gene clusters with divergent spatial patterns (Fig. 3d, Supplementary Fig. 12). To further benchmark the spatial effects, we took advantage of the vertical layered structure of MTG replicate 1, which implies that the spatial effect should capture trends along the vertical axis (true positive) but not along the horizontal axis (false positive). We adopted two non-parametric metrics, Kendall’s tau and Spearman R, to evaluate the dependency between each gene and the spatial coordinates. Apart from our approach (*SIMVI-archetype, DML*), we additionally included six baselines: *normalized counts (NC), scVI NC, NCEM NC*, cell-type conditioned linear regression using SIMVI spatial variation (analogous to the c-SIDE [42] setup, *SIMVI-LR*), cell-type conditioned linear regression using SIMVI archetype representation (*SIMVI-archetype, LR*), and an ablation of DML that substitutes SIMVI intrinsic variation with scVI embedding (*scVI ablation\**). We observed that *SIMVI-archetype, LR* and *SIMVI-archetype, DML* emerge as the two top performers in the benchmark, with DML showing favorable performance in identifying spatial-dependent genes (4/7 gene clusters excluding cluster 0) while filtering out false positives (6/8 gene clusters, Fig. 3e, Supplementary Fig. 13). Furthermore, substitution of SIMVI intrinsic variation with the scVI embedding (*scVI ablation\**) greatly hampered the performance and obscured true spatial patterns (Supplementary Figs. 12-13). This underscores the essence of removing spatial information from intrinsic variation for the spatial effect estimation task. Further gene pathway analysis underscores functional distinctions among these genes exhibiting different spatial patterns (Fig. 3f).

In the Slide-seqV2 mouse hippocampus data, a number of spatial niches also exhibit distinct intrinsic states, presumably due to the lower inherent resolution of the data. This constitutes a case where the positivity condition may be violated and examining the positivity index is essential. The spatial effect analysis of the Slide-seqV2 mouse hippocampus data primarily identified genes upregulated within the V3 upper region (Supplementary Fig. 14a). Similar to the results in MERFISH data, the spatial effects of these genes show cleaner spatial patterns in visualizations compared with normalized original counts, especially for genes exhibiting high sparsity (Supplementary Fig. 14d-e). Furthermore, the positivity analysis identifies potential violation of the positivity assumption in CA, DG, MH, and a layer between CA1 and cortical regions that primarily consist of oligodendrocytes (Supplementary Fig. 14b). These regions are indeed composed of cells with distinct intrinsic states and spatial environments simultaneously, supporting the validity of the approach. The spatial regions violating positivity (archetypes with high positivity index) remain consistent across different clustering resolutions (Supplementary Fig. 14c).

### SIMVI illuminates B cell dynamics in Slide-tags human tonsil

To further highlight the importance of disentangling variations in the study of complex spatial processes, we applied SIMVI to the recently published Slide-tags human tonsil dataset (Fig. 4a) [10]. As expected, the intrinsic variation generated by SIMVI well captured different cell types in the sample (Fig. 4b), while its spatial variation grouped biologically meaningful niches, which we annotated as B cell zone, GC dark / light zone, GC boundary that enriches Tfh cells, and T cell zone (Fig. 4c).

**Fig. 4.**
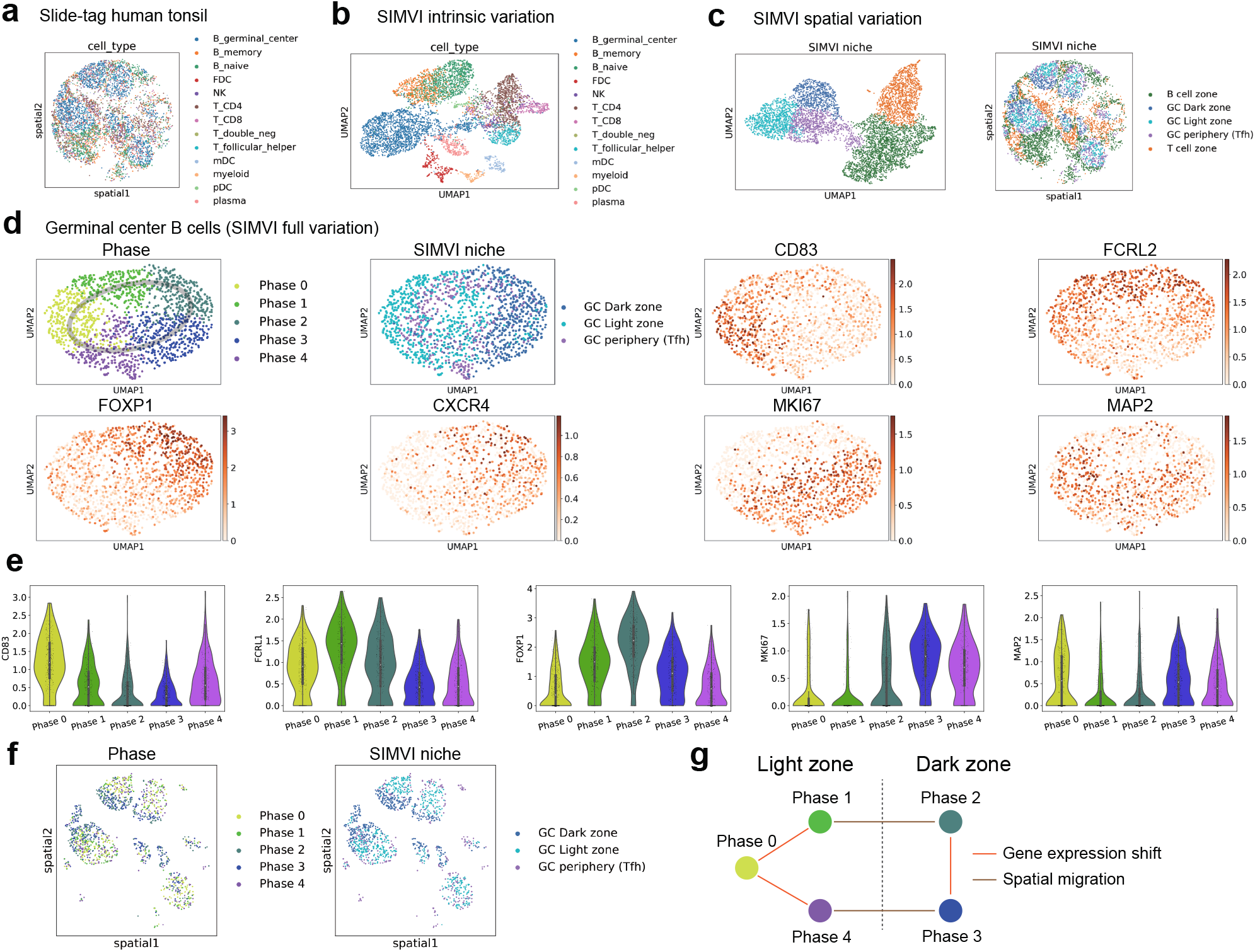
SIMVI identifies cell niches and dynamics in human tonsil. **a**. Spatial visualization of the dataset colored by cell types. **b**. UMAP visualization of the SIMVI intrinsic variation colored by cell types. **c**. UMAP and spatial visualization of the SIMVI spatial variation, colored by annotation derived from SIMVI spatial variation. **d**. UMAP visualization of the SIMVI full variation (concatenation of SIMVI intrinsic and spatial variation) for germinal center B cells, colored by phase annotation, spatial niche, and log normalized expression of representative genes. **e**. Violin plot showing patterns of representative gene log-normalized-expression. **f**. Spatial visualization of germinal center B cells, colored by phase and niche annotation. **g**. Illustration of the phase transition graph.

Our primary focus in the dataset is on germinal center B cells. These cells initially undergo rapid proliferation and somatic hypermutation (SHM) in the dark zone [43, 44]. Following SHM, B cells migrate to the light zone, where they receive survival and proliferation signals. This cycle repeats until the B cells mature, at which point they either undergo apoptosis or survive based on their antigen-binding efficiency [43, 44]. As such, we expect a cyclical organization of germinal center B cells. Using SIMVI’s full variation, we successfully annotated different phases of B cells, which indeed exhibit a circular structure (Fig. 4d). These phases are also characterized by sequentially activated genes (Fig. 4e, Supplementary Fig. 15c), and can be identified via standard pipelines (Supplementary Fig. 15a-b).

The spatial locations of germinal center B cells reveal intricate dependencies with their gene expression states. Among the five annotated phases, B cells from phases 2 and 3 were predominantly found in the dark zone, while those from other phases were mainly located in the light zone (Fig. 4f, Supplementary Fig. 15d). Combining with the phase adjacency relations (0-1-2-3-4-5-0, Fig. 4d-e), we were able to differentiate phase transitions caused by spatial migration from those driven by shifts in gene expression (Fig. 4g). Such a distinction explicitly requires two different representations encoding spatial information and cellular states respectively. Finally, Although external spatial annotations for benchmarking are lacking, an assessment using SIMVI-curated labels demonstrates that SIMVI consistently identifies spatial structures within the tonsil and the phases of germinal center B cells, compared with other methods (Supplementary Fig. 15e). Altogether, these findings underscore the unique strength of SIMVI disentangled representations in revealing distinct mechanisms underlying cellular state transitions.

### SIMVI reveals niches and spatial-dependent epigenetic states in human melanoma

Melanoma is the most common cause of skin cancer related fatalities, resulting in over 7000 deaths per year in the United States [45]. In recent years, the death rate has dropped dramatically due to the advent of immune checkpoint inhibitors which reverse T cell exhaustion and disinhibit effector T cells, among other functions [46]. However, not all patients respond to these therapies, and extensive efforts are ongoing to understand mechanisms of resistance or sensitivity. Apart from the heterogeneity across individuals, substantial heterogeneity is also observed within each tumor, which may promote drug resistance and the transition to pro-metastatic cellular states. In this work, we applied SIMVI to two datasets profiling human melanoma samples, to better understand the underlying biology at both individual and cohort levels.

The first dataset we employed is the spatial multiome melanoma dataset from the Slide-tags work [10]. It consists of a melanoma sample from an individual patient, profiling both spatial gene expression and chromatin accessibility. The dataset is particularly well-suited for exploring tumor heterogeneity within distinct spatial niches and elucidating the role of epigenetic regulation in defining different tumor states. Analyses of the dataset in the original study revealed two tumor subpopulations (tumor 1 and tumor 2) that exhibit distinct gene expression patterns and are also spatially segregated (Fig. 5a-b). We first applied SIMVI (and other baseline methods) on the gene expression modality alone to extract intrinsic and spatial variations. The cell type heterogeneity is well captured by SIMVI intrinsic variation, which further enforces latent space proximity across tumor cells consistent with cell ontology, compared with the scVI baseline (Fig. 5b, Supplementary Fig. 16d).

**Fig. 5.**
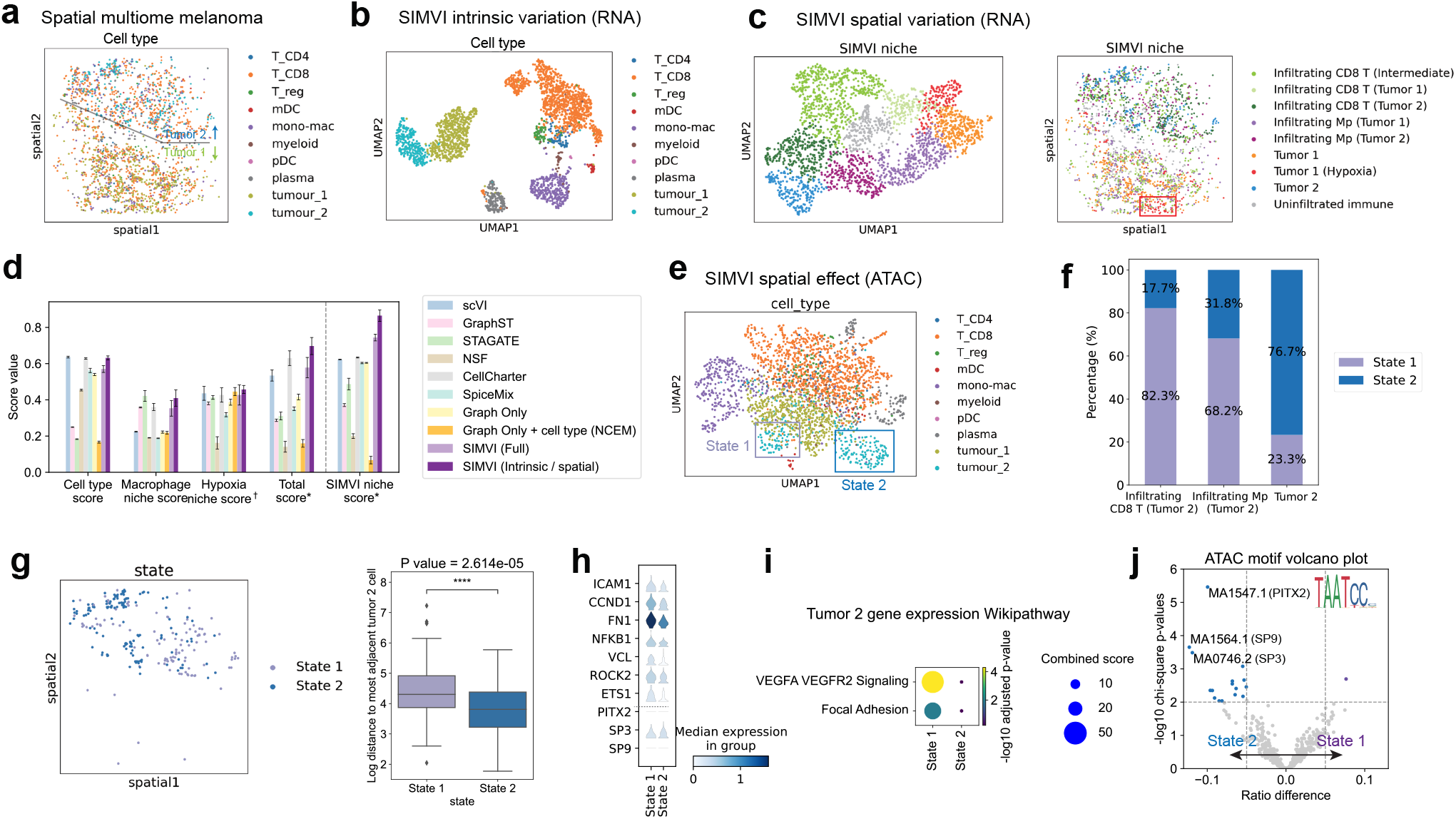
SIMVI uncovers functional niches and potential epigenetic reprogramming states in human melanoma. **a**. Spatial visualization of the dataset colored by cell types. **b**. UMAP visualization of the SIMVI intrinsic variation colored by cell types. **c**. UMAP and spatial visualization of the SIMVI spatial variation, colored by annotation derived from SIMVI spatial variation. **d**. Bar plots showing metric scores for the dataset. The bar heights represent the average performance across 10 different random seeds, with the error bars showing standard errors. *†* Each individual metric from all experiments is scaled to the range [0,1], then the final score is calculated as the average of these rescaled values. *: The score is obtained by first averaging the individual score values for each experiment, then rescaling the average score across all experiments to the range [0,1]. **e**. UMAP of the SIMVI spatial effect of the ATAC modality, colored by cell types. **f**. Bar plot showing distributions of tumor 2 state 1 / 2 in spatial niches. **g**. Spatial distribution of tumor 2 state 1 / 2 cells, and boxplot showing each cell’s log distance to nearest tumor 2 cells. n = 108, 139 for state 1 and 2 cells respectively. A Mann-Whitney test was performed. ****: p-value *<* 1 *×* 10^−4^. **h**. Stacked violin plot showing log normalized gene expression in state 1 and 2. Above the grey dashed line: representative genes in VEGFA-VEGFR2 signaling and focal adhesion pathways. Below: transcriptional factors whose corresponding motifs were enriched in differential peaks. **i**. Dotplot showing GSEA Wikipathway enrichment for differentially expressed genes across state 1 and 2. **j**. Volcano plot showing enriched motifs across differential ATAC peaks across the two states of tumor 2. X axis indicates the difference of a motif’s frequency in state 1 / 2 differential peaks.

The SIMVI spatial variation further depicts refined microenvironments within the sample, including niches primarily composed of tumor cells and those involving infiltrating immune cells. In particular, it differentiates two intermediate states of the immune infiltrating microenvironment: one involves an non-infiltrating intermediate state physically between two tumors (gray cluster in Fig. 5c). The other one shows a continuous shift with intermediate states found in both tumor 1 and 2 regions (Fig. 5c). Furthermore, the spatial variation discriminates a spatial niche in tumor 1 with highly expressed *SLC6A8* and *MIR3681HG* (Supplementary Fig. 16a-c), which we labeled as “hypoxia” due to the role of *SLC6A8* in metabolism and previous literatures highlighting its upregulation in hypoxic tumor cells [47].

Spatial effect analysis on the dataset using the gene expression modality alone reveals various genes influenced by the spatial context. One notable example is the macrophages that exhibit differential expression in tumor 1 and tumor 2 niches (*AQP9, MRC*, Supplementary Fig. 17a-c). We next benchmarked the quantitative performance across different methods in identifying cell types and macrophage / hypoxia niches based on external annotation labels (See Methods). Our results show that SIMVI excels in both cell type identification and spatial niche characterization, achieving the highest overall score. Moreover, multiple runs of SIMVI result in highly consistent spatial niches, marked by a notably high SIMVI niche score (See Methods, Fig. 5d). The advantage of SIMVI disentangled representations over SIMVI full variation further highlights the disentanglement ability of SIMVI and the benefits of utilizing disentangled representations (Fig. 5d).

Due to flexible design of the SIMVI spatial effect estimation procedure, we were able to extend the procedure to estimate the spatial effect for any single-cell-level quantities of interest. Noting the high noise of the ATAC-seq dataset, we performed spatial effect estimation on its Latent Semantic Indexing (LSI) components, aiming to reveal a global organization of spatial effect clusters. Surprisingly, our results highlight two distinct clusters of tumor 2 in the epigenetic spatial effect space, which we termed as state 1 and 2 (Fig. 5e). State 1 was mostly observed in immune infiltrating microenvironments, whereas state 2 was primarily localized in the non-immune-infiltrating tumor regions (Fig. 5f). Compared with state 2, cells from state 1 also exhibited more spatially dispersed distributions (Fig. 5g).

We further investigated the functions of differential genes / peaks across state 1 and state 2 cells. The gene expression differences are notably distinctive in the VEGFA-VEGFR2 signaling and focal adhesion pathways (Fig. 5i). These pathways are crucially linked to the metastatic capability and proliferation of tumor cells. The ATAC-seq differential peaks between these two states display diverse yet subtle functional enrichments, and relatively attenuated TSS accessibility in state 1 (Supplementary Fig. 18). Our motif analyses identified transcriptional factor motifs enriched specifically in state 2 (in particular the PITX2 motif, Fig. 5j), yet the gene expression changes of these TFs were not observed (Fig. 5h). These results provide evidences for a epigenetic reprogramming state in tumor 2 cells, which also exhibits a focused shift in gene expressions. This finding was not noted in the original study, likely relevant to substantial noise in the ATAC data that even obscured the distinction between tumor 1 and tumor 2 (Supplementary Fig. 17f,g). These results highlight the power of SIMVI in characterizing spatial-dependent states in spatial multi-omics datasets.

### SIMVI uncovers spatial interactions in cohort Melanoma data

The second dataset we employed is our newly collected CosMx samples from 25 melanoma patients treated with immune checkpoint inhibitors with various outcomes (Fig. 6a). The phenotype diversity within this single spatial array makes it an ideal case to systematically understand melanoma-associated spatial biology at a cohort level. In the dataset, we observed reasonable overlap for non-tumor cells across patients, while a huge heterogeneity of tumor cells across patients is still evident (Supplementary Fig. 19a). Further exploratory analyses revealed a cell type composition difference across patients with different outcomes (Supplementary Fig. 19b).

**Fig. 6.**
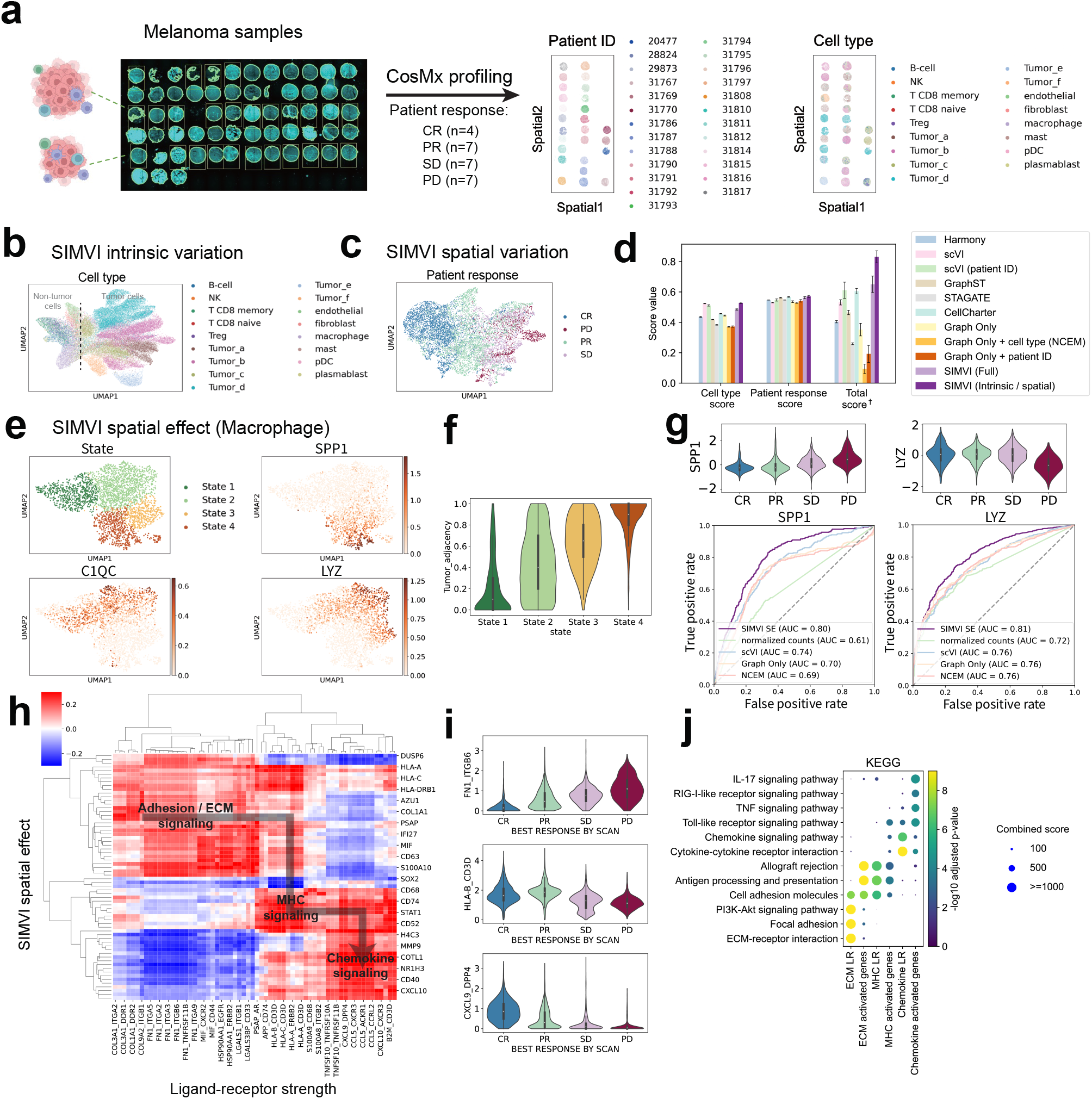
SIMVI characterizes macrophage subtypes and reveals cell interaction landscape in cohort-level spatial melanoma profiles. **a**. Overview of the dataset (left), spatial visualization of the data colored by annotated patient ID and cell types (right). **b**. UMAP visualization of the SIMVI intrinsic variation, colored by cell types. **c**. UMAP visualization of the SIMVI spatial variation, colored by response to treatment. The response to treatment was defined as disease progression (PD), stable disease (SD), partial or complete response (PR, CR), as described in prior publications [48]. **d**. Bar plots showing metric scores for the dataset. The bar heights represent the average performance across 5 different random seeds, with the error bars showing standard errors. *†* Each individual metric from all experiments is scaled to the range [0,1], then the final score is calculated as the average of these rescaled values. **e**. UMAP visualization of the SIMVI spatial effect, colored by annotated states and spatial effect of representative genes. **f**. Violin plot showing the relative tumor adjacency (defined as the tumor cell ratio within nearest 10 neighbors) across different states. **g**. Violin plots showing the spatial effect of *SPP1* and *LYZ* grouped by patient outcomes (upper); Receiver Operating Characteristic (ROC) curves comparing SIMVI spatial effects of *SPP1* and *LYZ* and other methods in classifying PD condition. **h**. Spearman correlation map across representative genes and ligand-receptor pairs. **i**. Violin plot of representative ligand-receptor strength across patient outcomes. **j**. Dotplot showing KEGG pathway enrichment analysis of genes and ligand-receptor pairs.

The heterogeneity across tumors includes differences of both intrinsic and spatial variations. Therefore, it constitutes a case where positivity, the assumption for estimating spatial effects, is not satisfied. Indeed, we observed a higher positivity index of all tumor cell subtypes compared with non-tumor cell types (Supplementary Fig. 19i). Nevertheless, in this challenging case, we found that the SIMVI intrinsic variation correctly merged tumor cells of the same subtypes (Fig. 6b, Supplementary Fig. 19). Meanwhile, the SIMVI spatial variation captures the pattern of patient responses in non-tumor cells, while separating tumor cells from different patients (Fig. 6c, Supplementary Fig. 19). In our benchmarking, SIMVI achieves state-of-the-art performance in accurately identifying both cell types (via intrinsic variation) and patient response labels (through spatial variation), leading to substantially improved overall performance (Fig. 6d, see Methods).

The proportion of macrophages among all cells remained consistent across patients with different outcomes, as opposed to other prevalent non-tumor cell types (Supplementary Fig. 19a). Moreover, tumor-associated macrophages (TAMs) are known as important indicators for patient outcomes [49, 50]. Based on these, we anticipate macrophages contribute to patient outcomes through spatial-dependent subpopulations, and therefore focused on spatial effect analysis for macrophages. Our analyses delineated four states in macrophages, characterized by canonical TAM markers (State 1: *C1QC+*, State 2: *C1QC+,LYZ+*, State 3: *LYZ+*, state 4: *SPP1+*) (Fig. 6e), showing a clearer representation than analyses using original counts (Supplementary Fig. 20a). These states well aligns with existing knowledge on TAMs [49], where *C1QC+, LYZ+* and *SPP1+* macrophages were categorized as tissue resident, classical tumor-infiltrating, and angiogenesis macrophages respectively [49]. We observed a monotonicly increasing tumor proximity in macrophage states from 1 to 4 (Fig. 6f). This observation is supported by visualization of a representative patient sample, where *SPP1* expression is primarily observed in macrophages adjacent to tumor, and the other two markers are mostly observed in immune niches (Supplementary Fig. 20c-e).

By comparing the expression of the canonical markers across patients with different outcomes, we observed an increase of *SPP1* and a decrease of *LYZ* in patients with the worst outcome PD (progressive disease, Supplementary Fig. 20b). The pattern was also observed in SIMVI spatial effects (Fig. 6g). A quantitative evaluation suggests a notable advantage of SIMVI spatial effect over alternative baselines in predicting patient PD (progressive disease) outcome by either *SPP1* or *LYZ* (Fig. 6g, see Methods).

Finally, we investigated the relationships between ligand-receptor strength and gene spatial effects among non-tumor cells. Strikingly, the correlation analysis reveals a statistically significant asymmetric relationship between ligand-receptor strength and spatial effects (Supplementary Fig. 21), indicating a potential latent trajectory for cell interaction machinery (Fig. 6h). Specifically, a high strength of adhesion / ECM signaling ligand-receptor (LR) pairs (dominant in innate immune cells and tissue resident cells) induces MHC signaling genes. The high level of MHC related LR across all cell types further activates chemokine signaling in lymphoid cells (Fig. 6h, Supplementary Fig. 21g). Other methods mostly recovered two regions and failed to identify the asymmetric pattern (Supplementary Figs. 21, 22). This is likely because alternative methods return “normalized expressions” convolving intrinsic and spatial effects. We also found that the LR strength from different stages in the machinery varies with the patient outcome (Fig. 6i), which may be explained by the cell composition difference across patient outcomes. This latent trajectory can be also represented by gene pathways, suggesting a directed information transfer landscape across cells in the tumor microenvironment (Fig. 6j).

## Discussion

We introduced SIMVI (Spatial Interaction Modeling using Variational Inference), a powerful approach to disentangle intrinsic and spatial-induced variations in spatial omics data. To the best of our knowledge, SIMVI is the first model that shows capability for the task, enabling further estimation of spatial effects at a single-cell level. SIMVI outperforms alternative methods in terms of various quantitative metrics and qualitative comparisons. We applied SIMVI to five real datasets from different tissues and platforms, including MERFISH human cortex, Slide-seqV2 mouse hippocampus, Slide-tags human tonsil, spatial multiome human melanoma, and CosMx cohort-level melanoma. SIMVI provides new biological insights for all analyzed datasets, with notable findings in the latter three. Given the rapid development of high-resolution spatial omics, we anticipate SIMVI to be of immediate interest to the spatial omics community.

SIMVI was designed to handle spatial omics data with single-cell resolution, such as imaging-based spatial omics data and high-resolution sequencing-based spatial omics data like Slide-tags [10] and Stereo-seq [11]. SIMVI may encounter limitations when applied to lower-resolution spatial omics datasets, but these challenges could potentially be addressed by incorporating complementary methods. One limitation is that these datasets have more than one cell in each pixel, which may obscure the cellular interactions and make the spatial gradient and gradual shifts in cell composition difficult to distinguish. Another limitation is that some of these technologies may have non-negligible gaps between pixels, which restricts the interpretation of local interactions between observed pixels. To address these limitations, advanced deconvolution methods such as Tangram [51], CARD [52], and DestVI [53] could help reveal the single-cell profile within each pixel by using scRNA-seq references. Moreover, computational techniques that model the spatial image and spatial transcriptomics datasets, such as XFuse [54] and TESLA [55], may be extended to provide imputations for cells not covered by pixels.

SIMVI uses a double machine learning (DML) procedure to estimate the spatial effect. While our implementation of DML was based on linear models, other more advanced machine learning models could possibly be adopted to improve the performance. However, the use of non-linear models might raise complexities related to model overfitting and interpretation [56]. Additionally, fine-tuning of trained SIMVI models on scRNA-seq datasets may facilitate niche annotation and spatial effect identification in single-cell sequencing datasets [57]. The spatial effect may require careful interpretation, as it could arise from different biological mechanisms such as cell migration and ligand-receptor interaction. Leveraging established gene interaction relationships (e.g. ligand-receptor) [58–62] can aid in interpretation of the observed spatial effects. Incorporating prior knowledge into the SIMVI model may also enhance SIMVI’s power in challenging spatial omics datasets, where intrinsic and spatial variations substantially coincide.

## Data availability

All datasets analyzed in this paper from previous publications are publicly available, with downloading and preprocessing instructions available in Methods. The newly generated dataset (the CosMx melanoma dataset) will be released soon.

## Code availability

We have made SIMVI available as a public open-source Python package, which can be accessed at https://github.com/KlugerLab/SIMVI.

### Acknowledgements

The authors thank Boaz Nadler for extensive discussions and Henry Li for helpful feedbacks. R.F. and Y.K. disclose support for the research of this work from NIH [U54AG076043, U54AG079759]. H.K. and Y.K. disclose support for the research of this work from NIH [P50CA121974]. Y.K. also discloses support for the research of this work from NIH [R01GM131642, UM1DA051410, and U01DA053628].

## Author contributions

M.D. conceived the study, developed SIMVI, and performed the computational analysis in the study. M.D. established the theoretical foundation of SIMVI with input from Y.K. H.K. provided melanoma tumor samples and clinical data. Y.K. and H.K. provided the CosMx melanoma dataset. D.S. provided feedback on the melanoma data analysis. M.D., H.K., and Y.K. wrote the manuscript with input from R.F.

## Methods

### The SIMVI model

We now provide a more detailed description of the SIMVI model. Let *X ∈* ℝ^*n×p*^ denote the count matrix of a spatial omics dataset, where *n* is the number of cells/points and *p* is the number of genes. The coordinate matrix of cells/points is represented by *C ∈* ℝ^*n×*2^, with *C*^*i*^ indicating the (*x, y*) coordinates of cell *i*. We preprocess the spatial information to construct a k-nearest neighbor graph (k = 10 throughout the study): *G* = (*V, E*), where the vertex set is *V* = {1, …, *n*}, and the edge set *E* consists of ordered pairs (*i, j*) with *i, j ∈ V*. The neighbors of cell *i* are defined as *N* (*i*) = {*j* | (*i, j*) *∈ E*}. Using these notations, we proceed to describe the generative model and inference procedure of SIMVI.

#### Generative process

In this work, we consider the following generative process for modeling the distribution of entries 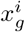 (cell *i*, gene *g*) in the count matrix of spatial omics data, given the graph *G*:

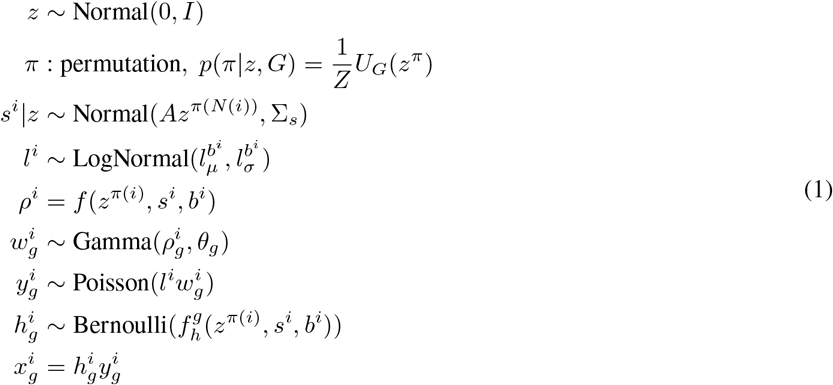

In the generative process, *z* = [*z*^1^, *z*^2^, …, *z*^*n*^] represents the intrinsic variation for all cells 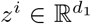 follows i.i.d. standard normal distributions, thus is independent of *G*. To further incorporate the possible spatial dependence, the generative process includes a probabilistic permutation *π* allocating *z*^*i*^ to spatial locations *π*(*i*). The permutation *π* is sampled from the distribution 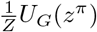, accounting for intrinsic cell information *z* and spatial proximity *G*. After the permutation, z^*π*(*i*)^. represents the intrinsic information of the cell at node *i*.

*s*^*i*^ represents the spatial variation of the cell at node *i*. It depends on the neighborhood intrinsic variation *z*^*π*(*N*(*i*))^, and follows a normal distribution with the form shown in (1), where the matrix 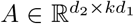, and the covariance matrix 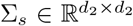 is positive definite. *b*^*i*^ represents experimental covariates such as batch label. The library size is modeled as a latent variable following log normal distribution with parameters 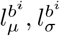, following the scVI design [34]. In practice, we have found the statistical estimation of library size *l*^*i*^ is usually sufficient. In this case, the generative process of library size *l*^*i*^ in (1) can be straightforwardly replaced with 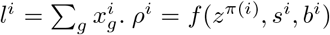 represents the expectation of scaled gene expression. The function *f* employs a soft-max layer so that *ρ*^*i*^ has positive entries that sum to 1. *θ ∈* ℝ^*p*^ specifies the gene-specific shape parameter of the Gamma distribution. 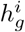 represents the zero inflation level. Together, 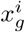 follows a zero-inflated negative binomial (ZINB) distribution, parametrized by functions of (*z*^*π*(*i*)^, *s*^*i*^, *b*^*i*^, *l*^*i*^). The generative process of SIMVI mostly follows the scVI framework [34, 63], with the addition of the probabilistic spatial assignment *π* and the spatial variation *s*.

### Approximate posterior inference of SIMVI

In order to infer the parameters in the generative process, we approximate the posterior distribution via variational inference. The posterior distribution is approximated to be of the following factorized forms:

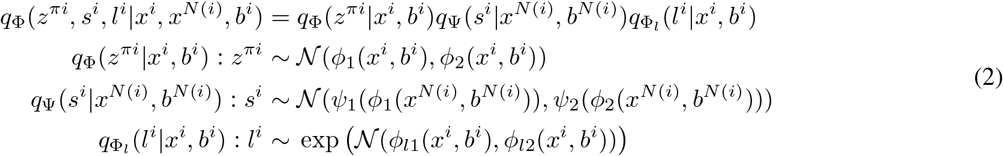

Here (Φ, Ψ, Φ_*l*_) denotes neural network weights that determine the variational posterior, composed of ((*ϕ*_1_, *ϕ*_2_), (*ψ*_1_, *ψ*_2_), (*ϕ*_*l*1_, *ϕ*_*l*2_)) as shown above. Our treatment of *z*^*i*^ and *l*^*i*^ aligns with the scVI family of models [34, 63]. In SIMVI, we additionally model the spatial variation *s*_*i*_ using transformed cell neighborhood gene expression *ϕ*_1_(*x*^*N*(*i*)^), where *ϕ*_1_ encodes the expectation term of intrinsic variation. Such a design enforces the spatial variation to be determined by the intrinsic variation in the cell neighborhood. We use a one-layer graph attention network (GAT) with dynamic attention [64] to model the variational posterior *ψ*_1_, *ψ*_2_. Finally, we note that the constructed variational posterior directly starts from *z*^*π*^ and does not explicitly model *z* and *π*. Consequently, the variational posteriors are best viewed as descriptive models of the previously outlined generative process.

After the factorization, we derive the evidence lower bound (ELBO) as follows. Instead of considering each cell, we consider ELBO of the full likelihood. As samples in the generative process are not independent, the full likelihood is not a simple sum of individual likelihoods. In the derivation of the ELBO, terms without an upper index represent those from all data points.

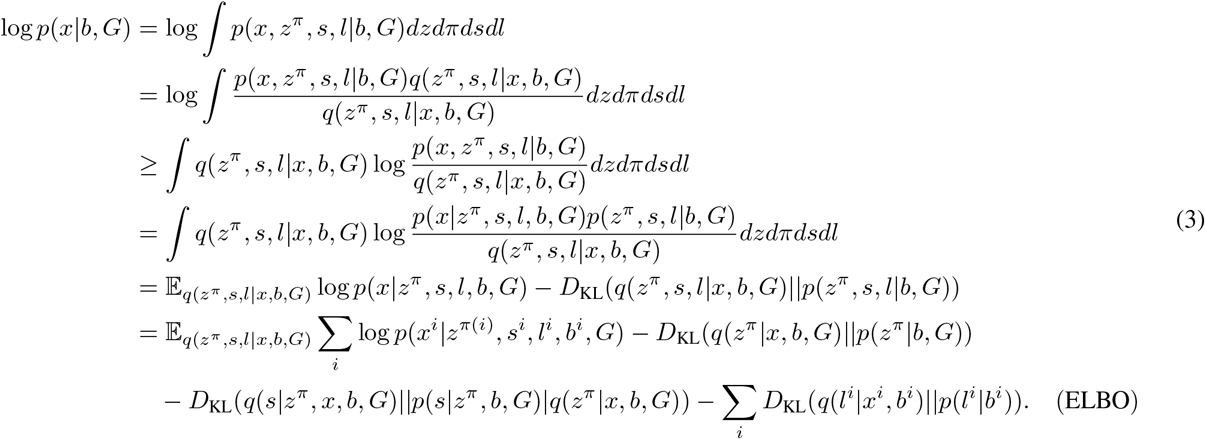

For the first KL divergence term, we have

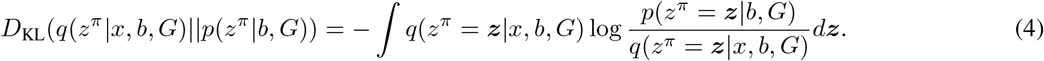

Considering the term *p*(*z*^*π*^ = ***z***|*b, G*), we have

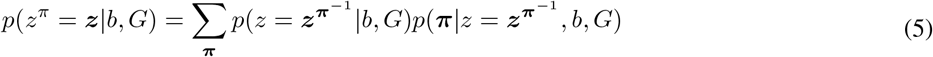

Because *z*^*i*^|*b, G* ~ *N* (0, *I*) are i.i.d. across different 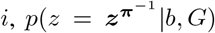 is a constant with respect to ***π***, and equals *p*(*z* = *z*| *b, G*). Moreover, from the definition of 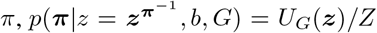 is also a constant with respect to ***π***. The total number of ***π*** is *n*!. Therefore we have

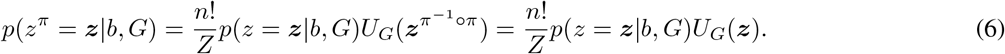

Altogether, this KL divergence term equals

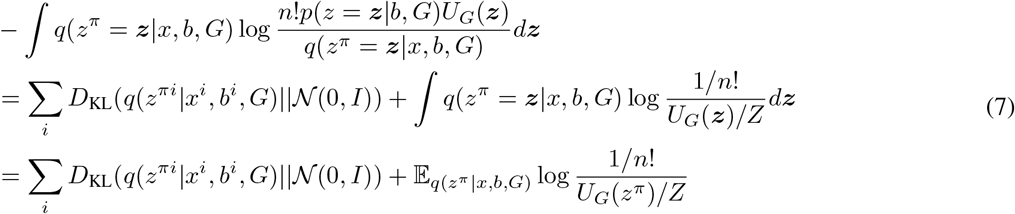

Our derivation shows that the KL divergence regarding *z* can be decomposed into two parts. The first part coincides with the KL divergence term in vanilla VAEs and scVI [34], while the second term is minimized when *U*_*G*_(*z*^*π*^) achieves maximum under the variational posterior *q*(*z*^*π*^ | *x, b, G*). In practice, we only used the first part in the loss function for two reasons. First, the function *U* is not tractable in practice; second, if the variational posterior *q*(*z*^*π*^ | *x*) well characterizes individual observations, then it should automatically capture the underlying colocalization pattern, i.e., *U*_*G*_(*z*^*π*^) is large.

For the second KL divergence term, we have

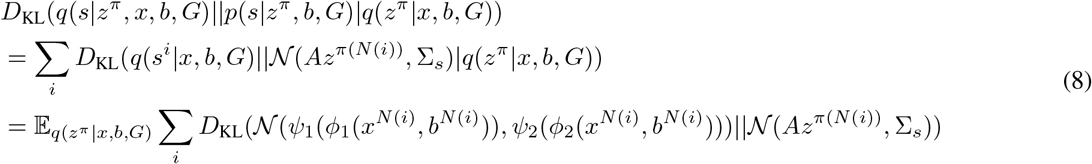

As *A*, Σ_*s*_ are not known in priori, the term is not tractable in general. Therefore, we instead formulate a penalty term regularizing the variational posterior towards standard normal distribution with a small coefficient *α* (0.01 in this study). This coincides with (8) when *A* = 0, Σ_*s*_ = *I*. Such a regularization with a small weight is also applied in state-of-the-art vision models such as latent diffusion [65].

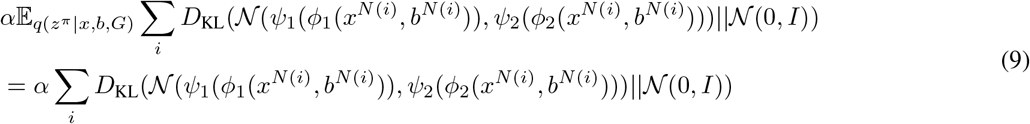

In summary, we have derived the (effective) ELBO of the SIMVI model, which can be indeed decomposed for each cell *i*, after dropping the term associated with *U*_*G*_:

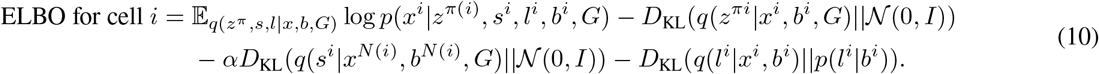

Optimizing solely ELBO is not enough to achieve disentanglement. In particular, the inferred intrinsic variation may encompass all information in the dataset, if no additional constraints are specified. Our theoretical analysis shows that, the model identifiability can indeed be achieved, when the intrinsic variation *z* encodes minimal information (Supplementary Note 1). In order to incorporate this into the SIMVI objective, we utilize an independence regularization term between *s* and *z*, and only regularize *z* by the term. We implement two versions of the regularization, utilizing closed-form mutual information and kernel maximum mean discrepancy (MMD) respectively. The expressions for both are introduced as follows.

1. **Closed-form mutual information**. Assuming [*z, s*] follows joint Gaussian distribution, the mutual information term can be derived analytically:

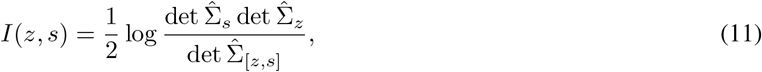

where 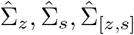 denote the sample covariance matrices for *z, s*, [*z, s*] respectively. If [*z, s*] follows a joint Gaussian distribution, then this term can be viewed as a closed-form estimator of mutual information. Even when [*z, s*] does not follow a joint Gaussian distribution, it still serves as a meaningful measure of the dependency between *z* and *s*.
2. **Kernel MMD based independence regularization**. We consider the regularization of the following form:

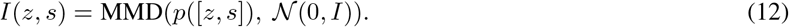

Where the function MMD is defined as:

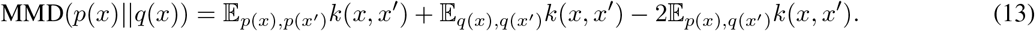

Here *k* indicates Gaussian kernel function with the denominator equal to the dimension number, consistent with the setting in InfoVAE [66]. The MMD based independence regularization jointly enforces Gaussianity and the independence between *z* and *s*.

To enforce meaningful disentanglement, we only apply the regularization term on *z*. The resulting optimization problem is formulated as follows:

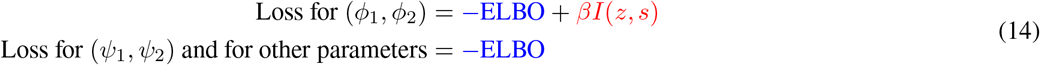

We finalize by noting that the train / validation set construction and the mini-batch setting for the SIMVI model is different from most scVI-based models [67, 68]. This arises from the fact that a general connected graph does not admit a natural partitioning scheme. In this work, in order to define train and validation sets or mini-batches, we adopt the semi-supervised node classification framework for graph neural networks [69]. Specifically, the full dataset is fed into the SIMVI model to compute the intermediate cell-level outputs (embeddings) for each cell. The cell-level outputs are then divided into training and validation sets, with each further segmented into mini-batches. During model training, the loss function is computed using only the embedding outputs from cells in the training set.

#### Permutation-based pretraining

SIMVI also provides an option to employ the denoising autoencoding scheme [70] as a pretraining step. Specifically, we first sample a subset of genes in each training batch, and then permute the gene values across cells. This step adds noise in the training data while still preserving the marginal distribution of genes. Then the model is trained to reconstruct the original data from this noisy version of data.

#### Spatial effect identification with SIMVI

The spatial effect, i.e., the gene expression changes due to spatial microenvironment, is usually determined by both intrinsic and spatial variations of a cell. We propose to estimate the spatial effect via continuous treatment effect estimation framework in causal inference [36]. The methodology we employ (archetype transformation, and double machine learning) is also described in Supplementary Note 2 with an emphasis on the mathematical foundation. A related task to ours is the treatment effect estimation problem for single-cell RNA-seq data [71, 72]; however, a key distinction in our case is that the “treatment”, spatial variation, is inferred from the model.

We first use archetypal analysis [39] to transform the spatial variation *s* into archetype weight representation *s*′ using the PCHA package [39]. The number of archetypes is customized across datasets in our study. We set *δ* = 0.1 consistent with the PCHA usage example to increase the tolerance for archetypes outside data support. We further adjust the default setting to reduce the number of iterations (conv crit=1E-5, maxiter=200 compared with the default conv crit=1E-6, maxiter=500). All other parameters are selected as PCHA defaults.

Then we use linear regression to fit the two models and obtain residuals 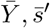 :

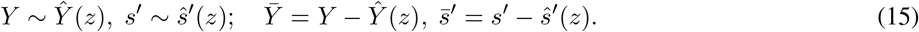

Here *Y* represents log normalized expression of a gene (or any compatible term of interest), *z* represents the covariate, which is the SIMVI intrinsic variation, optionally concatenating other covariates; *s*′ represents the transformed archetypal weight vector. After we obtained residuals 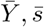, we run a linear regression model

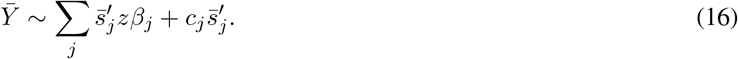

After the model parameters are obtained, the spatial effect can be obtained as:

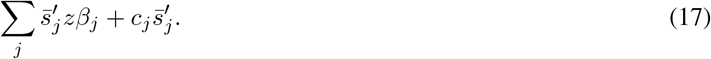

The coefficient of determination, *R*^2^, for intrinsic variation is represented as 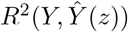, while the *R*^2^ for spatial variation is denoted by 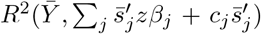. All linear regression models in our study were implemented using `sklearn.linear_model.LinearRegression`.

We further developed an “individual mode” to estimate the spatial effect for each archetype. In this case, we have one regression model for each archetype on the residual 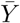

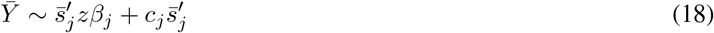

After fitting all regression models, the spatial effect is still obtained as the sum 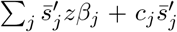 whereas the *R*^2^ for each gene is defined as the max *R*^2^ among all archetypes. Our analysis revealed high concordance between spatial effects generated by the original and individual modes (Supplementary Fig. 11c). Therefore we used the individual mode throughout the study, leveraging its enhanced capacity to estimate spatial effects for each archetype. To highlight the distinction between intrinsic variable genes and spatial-induced genes in the *R*^2^ scatter plot, we labeled genes with intrinsic variation *R*^2^ larger than a threshold as “Intrinsic-specific”. We further labeled outlier genes from Huber regression (scaled residual larger than a threshold) as “Spatial-induced”.

Finally, we provide an option to derive label-based positivity scores to evaluate the feasibility of spatial effect estimation for each cell. A mathematical formulation is provided in Supplementary Note 2. The cell-level positivity index involves the selection of two thresholds: thres1, which is used in filtering out archetypes, and thres2, which is applied in filtering out individual cells. An evaluation on the real Slide-seqV2 dataset suggests that the spatial regions violating positivity (archetypes with high positivity index) remain stable with respect to the clustering resolution (Supplementary Fig. 14c).

### MERFISH human cortex dataset

We downloaded the data from https://datadryad.org/stash/dataset/doi:10.5061/dryad.x3ffbg7mw. We used the following samples from the dataset: MTG sample: H18.06.006.MTG.4000 rep1-3, with 11059 cells in total and 4000 genes; STG sample: H19.30.001.STG.4000 rep1-3, with 14924 cells in total and 4000 genes. The cell type label was provided in the dataset named cluster L1. We annotated the layers by manual segmentation, referring to the cell subtype annotations provided in the dataset (cluster L2), which additionally labeled layers of excitory neurons and subtypes of inhibitory neurons. We further derived a local niche annotation based on *MYH11* expression. In the MTG dataset, we labeled cells with log normalized *MYH11* expression ≥ 1 as “MYH11+”, and the remaining as “MYH11-”. In the STG dataset, due to the overall higher sparsity, we labeled all cells expressing *MYH11* as “MYH11+”, and the remaining as “MYH11-”. We conducted a thorough parameter sweeping study using MTG and STG samples to better understand the SIMVI performance under different model configurations, with details available in Supplementary Note 3.

We included a variety of methods in the benchmarking apart from SIMVI, including Harmony, scVI, MEFISTO, NSF, SpiceMix, GraphST, STAGATE, CellCharter [21–24, 26, 34, 38]), and included a list of baselines that adds a batch integration step for methods do not address batch correction (GraphST + Harmony, STAGATE + Harmony, SpiceMix + Harmony). We further tested batch-correcting scVI-backbone models that do not incorporate intrinsic variation but include covariate labels instead (Graph Only; Graph Only + cell type 1; Graph Only + cell type 2). We tested all methods using the same set of 10 random seeds. For all methods that transform spatial information into graphs (SIMVI, SpiceMix, GraphST, STAGATE, CellCharter, GraphST + Harmony, STAGATE + Harmony, SpiceMix + Harmony, Graph Only, Graph Only + cell type 1, Graph Only + cell type 2), we used a k-NN graph encoding spatial proximity with k=10. For the remaining methods that require spatial locations (MEFISTO, NSF), we combined the datasets and positioned different samples at sufficiently distant locations. For scVI-based approaches (SIMVI, scVI, CellCharter, Graph Only, Graph Only + cell type 1, Graph Only + cell type 2) that involve mini-batch training, we used the same number of training epoches (100) and the same batch size (500). We used the default set of parameters of SIMVI (Supplementary Note 3), with 25 pretraining epoches. We used default settings for Harmony and applied it on the PCA embedding of log normalized expression of all 4000 genes. For scVI, we used a compatible setting with the SIMVI setting that is also similar to scVI default, with latent space dim = 20, number of encoder and decoder layers = 2. For methods that involve normalization (MEFISTO, NSF, GraphST, STAGATE, SpiceMix), we used log normalized counts by Scanpy default. For MEFISTO and NSF, we used 500 inducing points (Gaussian likelihood for NSF), with the spatial variation dim = 10. We observed that the version of NSF we used that outputs both spatial and non-spatial variations (NSFH) outputs NaN for the non-spatial part. Therefore, only the spatial part is used in our benchmarking. We used default parameter settings for GraphST and STAGATE. For SpiceMix, we used k-means initialization, and a compatible parameter setting with SIMVI, namely K = num pcs = 20. For CellCharter, we leveraged the trained scVI embeddings and used the first order neighborhood and set the aggregation function as mean aggregation. For (GraphST + Harmony, STAGATE + Harmony, SpiceMix + Harmony), we utilized the formerly trained embeddings from GraphST, STAGATE, and SpiceMix as inputs for Harmony, applying a fixed random seed for Harmony. For (Graph Only, Graph Only + cell type 1, and Graph Only + cell type 2), latent space dim = 20, and all remaining applicable parameter settings were consistent with those of SIMVI.

We defined a list of benchmarking scores for evaluating performances across different methods. Each score is an ensemble of batch correction or biological preservation metrics implemented in scIB [73]. Specifically, we used graph connectivity, silhouette batch / label, and KBET (k-nearest-neighbor batch-effect test) as batch correction metrics, and Leiden NMI (normalized mutual information), Leiden ARI (adjusted Rand index), Silhouette label as biological preservation metrics. All the metrics were implemented by the scib-metrics Python package [73]. Batch correction metrics require both batch and biological annotation labels as inputs, and were applied to datasets containing multiple batches. In contrast, biological preservation metrics only require biological annotation labels, and were applied to both single-batch and multi-batch datasets.

We defined three aggregation rules for summarizing metrics as final scores. Scores without auxillary labels indicate that it is a result of averaging metrics without rescaling. Scores labeled as * indicate that it is a result of averaging metrics and then rescaling the score to have min 0 and max 1. Scores labeled as *†* indicate that it is an average of rescaled metrics with min 0 and max 1. Using the aggregation rules, we derived four scores evaluating batch correction, cell type preservation, layer preservation, and local structure (MYH11+/-label) preservation respectively. These scores are further summarized as total scores by the described aggregation rules. For SIMVI, we assessed batch correction and cell type preservation using the intrinsic variation, while layer and niche preservation performances were evaluated based on the spatial variation.

We performed the archetype transformation in the dataset with number of archetypes = 7. Then we computed the spatial effect and associated *R*^2^s as previously described, with covariates as the concatenation of SIMVI intrinsic variation and one-hot labeling of the cell subtype annotation. Genes with log normalized mean expression larger than 0.1 (1561 genes in total) were used in the analysis. The spatial effect of astrocytes from replicate 1 was binned into 25 segments based on spatial Y coordinates and subsequently clustered using k-means (k = 8) to generate gene clusters 0-7. For baseline models, *normalized counts (NC)* represents log normalized original expression, and *scVI / NCEM NC* represent the normalized expression returned by either a scVI or a Graph Only + cell type model as previously described. We implemented two additional linear regression baseline models: one regressing log-normalized expression against the concatenation of cell subtype label and SIMVI spatial variation, and another regressing log-normalized expression against the concatenation of cell subtype and archetype weights of SIMVI spatial variation. We computed Spearman correlation and Kendall’s tau between spatial coordinates and each gene from different models’ output (without binning) of replicate 1 astrocytes. We computed the Wikipathway enrichment using EnrichR wrapped by GSEAPY [74, 75].

### Slide-seqV2 mouse hippocampus dataset

We acquired the annotated dataset in AnnData format from Squidpy [76]. As the dataset does not provide raw counts, we additionally accessed the raw count matrix from the Broad single-cell portal. An anatomic map from the Allen Mouse Brain Atlas [77] (P56, Coronal, Image 74, plate = 100960228) was downloaded and shown in Fig. 2e. In the analyses from the original work [8], it has been noted that not all pixels correspond to individual cells. Instead, they can represent subcellular structures such as dendrites [8]. As a result, we applied a strict filtering scheme by including the top 2000 highly variable genes (via Scanpy default) and included cells with at least 20 highly variable genes. Our preprocessing scheme yielded to 32838 cells and 2000 genes. As the dataset only contains one batch, a smaller list of methods were included for comparison (scVI, MEFISTO, NSF, SpiceMix, GraphST, STAGATE, CellCharter, Graph Only, Graph Only + dominant cell type label (NCEM) [21–24, 26, 30, 34]). We note that here due to the absence of batch labels, the Graph Only model here does not have condition variables, and the Graph Only + cell type model uses the dominant cell type label from prior deconvolution. MEFISTO was not included due to its extensive time and memory usage. Due to the larger dataset size, we chose a larger batch size (2000) for scVI-based approaches. Other details of implementation were the same as those adopted in the MERFISH benchmarking.

We visualized SIMVI and two other top performers’ (GraphST, CellCharter) Leiden clustering results. The clustering resolutions were manually picked for best consistency with the Allen Mouse Brain Atlas annotation. Based on the SIMVI clustering results, we defined two scores. One is the niche preservation score that accounts for all clusters shown in the SIMVI panel of Supplemental Fig. 8e (All niche score); the other is based on the subset of SIMVI clusters that show clear spatial structures and aligns with the Allen Brain Atlas Annotation (cluster 7,8,9,10,11,12,13,14, Supplementary Fig. 8e), which we term as Selected niche score. The scores were defined as aggregation of biological preservation metrics using cluster labels. To avoid confirmation bias, in the benchmarking, we excluded the SIMVI run used to generate the clustering results.

We further used the existing deconvolution results provided in Squidpy to derive an annotation for CA / DG regions and their neighborhoods and compared with the clustering results. Specifically, for each cell, we computed a score by summing its own deconvolution ratios (CA / DG) with 1.5 times the average of its 10 nearest neighbors. We then applied thresholds to the scores, resulting in spatially smooth structures corresponding to CA and DG regions (Supplementary Fig. 8b-c). We further calculated the relative distance between CA, DG and their neighborhood by Silhouette label score. For spatial effect analysis, we used the same parameter setting as those adopted in MERFISH data analysis (with the exception of archetype number = 20), with covariates as the concatenation of SIMVI intrinsic variation and the prior deconvolution results. We further calculated the binary positivity index with thres1 = 0.95, thres2 = 0.5.

### Slide-tags human tonsil dataset

We downloaded the dataset along with metadata from Broad Institute Single Cell Portal under the accession number SCP2169 [10]. The data consists of 5778 cells. We selected the top 4,000 highly variable genes from the dataset by Scanpy default. We applied the same set of methods as those in the Slide-seqV2 section. The training settings of all methods were consistent as those used for the MERFISH dataset, with the exception of the batch size for scVI-based methods (SIMVI, scVI, Graph Only, Graph Only + cell type) was set to 1000. We used the Leiden clustering result from SIMVI spatial variation to define the SIMVI niche shown in Fig. 4c. To define the germinal center B cell subset used in Fig. 4d, we first selected the cells that were labeled as germinal center B cells in the original annotation, and then filtered out possible doublet clusters that express *CD247* and *FDCSP* respectively. We used the Leiden clustering result from SIMVI full variation to define the germinal center B cell phases. For the likelihood analysis in Supplementary Fig. 15d, we used MELD [78] to compute a graph filter that smooths a signal on graph defined by embeddings. We used MELD to derive a continuous dark zone likelihood by applying the graph filter estimated from SIMVI spatial variation on the one-hot dark zone label. The parameter *β* is set as 80, and all other parameters were consistent with the MELD default.

To benchmark across different methods, we derived three scores evaluating the cell type preservation, niche preservation, and GC phase preservation (within the set of GC B cells) respectively. The definitions of these scores are consistent with the descriptions in previous sections. To minimize the confirmation bias, we did not include the SIMVI run that was used to generate the annotation results.

### Spatial multiome melanoma dataset

We downloaded the dataset along with metadata from Broad Institute Single Cell Portal under the accession number SCP2176 [10]. The data consists of 2535 cells. We selected the top 2,000 highly variable genes from the dataset by Scanpy default. We applied the same set of methods as in the Slide-seqV2 and Slide-tags human tonsil section. Apart from disentangled SIMVI representations, we further incorporated the full SIMVI variation in the benchmarking. We used the Leiden clustering result from SIMVI spatial variation to define the SIMVI niche shown in Fig. 5c. We derived the tumor niche label and the micoenvironment label (hypoxic or not for tumor 1) by manually segmenting the spatial region (Supplementary Fig. 16a). All implementation details were consistent with those in Slide-tags human tonsil data. To benchmark across different methods, we defined four scores evaluating the cell type preservation, macrophage state preservation (across tumor 1 and tumor 2 regions), hypoxia state preservation (within tumor 1), and SIMVI niche preservation (using the clustering labels derived from SIMVI) respectively. The definition of these scores are consistent with the descriptions in previous sections. To minimize the confirmation bias, we did not include the SIMVI run used to generate the SIMVI niche annotation in the benchmark.

For the ATAC modality preprocessing, we performed TF-IDF transformation on the provided peak matrix [10] and computed the LSI components. We observed that removing the first LSI component has almost no effect on the data embedding (Supplementary Fig. 17f). Therefore we used all LSI components in subsequent analysis. We computed the spatial effect of the LSI components and log normalized gene expressions (number of archetypes = 8 in both cases) with covariates as the concatenation of SIMVI intrinsic variation and one-hot labeling of the cell type annotation. The state 1 and 2 were annotated by Leiden clustering of the ATAC spatial effect (Supplementary Fig. 17g). We performed differential analysis on the gene expression and ATAC peaks using `sc.tl.rank_genes_groups` with default settings. We next computed the Wikipathway enrichment for differential ATAC peaks (transformed to overlapping genes by Ensembl REST API [79] and gene transcription starting sites by the main annotation file from GRCh38.p14 [80]) and differentially expressed genes using EnrichR wrapped by GSEAPY [74, 75]. Motifs were annotated with JASPAR 2020 using the ChromVar implementation [81, 82].

### CosMx melanoma dataset

The study was approved by the Yale Human Investigation Committee protocol and conducted in accordance with the Declaration of Helsinki. Tissue microarray blocks were constructed as previously described [48], containing tumor samples from 60 patients treated with immunotherapy. Slices from the block were submitted to to NanoString for CosMx spatial profiling. Twenty five samples were randomly selected for analysis. The dataset used contains samples from 16 male patients and 9 female patients, ages ranging from 35-90. Among these patients, 11 patients received ipilimumab and nivolumab in combination (IPI+NIVO), 13 received pembrolizumab (PEMBRO), and one patient received nivolumab (NIVO) alone. CosMx Human Universal Cell Characterization RNA Panel was used as the SMI reagent. This panel included genes for cell typing and mapping (243 genes), cell state and function (269 genes), cell-cell interaction (435 genes), and hormone activities (46 genes). No statistical methods were used to predetermine sample size. The data was divided into 11 categories of non-tumor cells and six subclasses of tumor cells by NanoString. The preprocessed dataset consisted of 56,761 cells and 960 genes.

The dataset is composed of only one batch but contains samples from different patients. Therefore we included a slightly different list for benchmarking (Harmony, scVI, scVI using patient label, GraphST, STAGATE, Graph Only, Graph Only + cell type, Graph Only + patient ID, SIMVI). NSF and SpiceMix were not included due to their extensive usage of time and memory, and previous benchmarking results showing that they prioritize intrinsic information and do not have a notable advantage over scVI. Apart from disentangled SIMVI representations, we further incorporated the full SIMVI variation in the benchmarking. We trained the SIMVI model with default settings with the pretraining epoch number = 75. Other training settings of all methods were consistent as those used for the MERFISH dataset, with the exception of the batch size for scVI-based methods (SIMVI, scVI, Graph Only, Graph Only + cell type) was set to 5000. We defined the cell type score as the scIB total score for batch correction (in this case patient label) and biological preservation for cell type, both defined consistently as described in previous sections. We also defined a patient response score to evaluate if a embedding of the CD8 T cell subset preserves the patient response label and maximally integrates different patients. We used only CD8 T cells here to remove the effect of cell composition difference across patient outcomes (Supplementary Fig. 19b), and noting the direct relevance between CD8 T cells and the immunotherapy treatment.

We computed the spatial effect for all cells (number of archetypes = 20) and calculated the continuous positivity index with thres1 = 0.9 (Supplementary Fig. 19i,j). We extracted the macrophage subset for analyses in Fig. 6e-g. For the ligand-receptor analysis, we used CellTalkDB [62] for extracting the ligand-receptor pairs. For each ligand-receptor pair, we computed the sum of the average neighborhood normalized ligand expression (k = 10) and the central cell normalized receptor expression as the ligand-receptor strength for each cell. We further computed the Spearman correlation map between the ligand-receptor strength and gene spatial effect within non-tumor cells. The rows / columns that has an max absolute value above 0.25 is preserved in Fig. 6h. We used `seaborn.clustermap` via the Scanpy wrapper to perform hierarchical clustering on correlation maps, with method=‘centroid’ and other settings were set to be default. The correlation map was further segragated into different regions (Supplementary Fig. 21e). We computed the KEGG enrichment for ligand-receptor pairs and genes in different regions of the correlation map using EnrichR wrapped by GSEAPY [74, 75].

### Empirical running time

All SIMVI experiments were conducted on a high performance Linux server using one NVIDIA A6000 GPU. The most extensive (in both time and memory) training session we ran (on the CosMx melanoma dataset with 56,761 cells and 960 genes) completed the 100 epoches training (75 of the epoches are pretraining epoches) with batch size 5000 in approximately 15 minutes. We also note that the permutation epoch typically requires more time due to its permutation sampling scheme.

## References

[1] Lohoff, T. et al. Integration of spatial and single-cell transcriptomic data elucidates mouse organogenesis. Nature biotechnology 40, 74–85 (2022).

[2] Zhang, M. et al. Spatially resolved cell atlas of the mouse primary motor cortex by merfish. Nature 598, 137–143 (2021).

[3] Fang, R. et al. Conservation and divergence of cortical cell organization in human and mouse revealed by merfish. Science 377, 56–62 (2022).

[4] He, S. et al. High-plex imaging of rna and proteins at subcellular resolution in fixed tissue by spatial molecular imaging. Nature Biotechnology 40, 1794–1806 (2022).

[5] Liu, Y. et al. High-spatial-resolution multi-omics sequencing via deterministic barcoding in tissue. Cell 183, 1665–1681 (2020).

[6] Liu, Y., Enninful, A., Deng, Y. & Fan, R. Spatial transcriptome sequencing of ffpe tissues at the cellular level. bioRxiv 2020–10 (2020).

[7] Zhang, D. et al. Spatial epigenome–transcriptome co-profiling of mammalian tissues. Nature 616, 113–122 (2023).

[8] Stickels, R. R. et al. Highly sensitive spatial transcriptomics at near-cellular resolution with slide-seqv2. Nature biotechnology 39, 313–319 (2021).

[9] Vickovic, S. et al. High-definition spatial transcriptomics for in situ tissue profiling. Nature methods 16, 987–990 (2019).

[10] Russell, A. J. et al. Slide-tags enables single-nucleus barcoding for multimodal spatial genomics. Nature 625, 101–109 (2024).

[11] Chen, A. et al. Spatiotemporal transcriptomic atlas of mouse organogenesis using dna nanoball-patterned arrays. Cell 185, 1777–1792 (2022).

[12] Wei, X. et al. Single-cell stereo-seq reveals induced progenitor cells involved in axolotl brain regeneration. Science 377, eabp9444 (2022).

[13] Gilbert, S. F. Developmental biology. (sinauer associates, Inc, 2010).

[14] Dries, R. et al. Giotto: a toolbox for integrative analysis and visualization of spatial expression data. Genome biology 22, 1–31 (2021).

[15] Weber, L. M., Saha, A., Datta, A., Hansen, K. D. & Hicks, S. C. nnsvg: scalable identification of spatially variable genes using nearest-neighbor gaussian processes. bioRxiv 2022–05 (2022).

[16] Hao, M., Hua, K. & Zhang, X. Somde: a scalable method for identifying spatially variable genes with self-organizing map. Bioinformatics 37, 4392–4398 (2021).

[17] Sun, S., Zhu, J. & Zhou, X. Statistical analysis of spatial expression patterns for spatially resolved transcriptomic studies. Nature methods 17, 193–200 (2020).

[18] Zhu, J., Sun, S. & Zhou, X. Spark-x: non-parametric modeling enables scalable and robust detection of spatial expression patterns for large spatial transcriptomic studies. Genome Biology 22, 1–25 (2021).

[19] Svensson, V., Teichmann, S. A. & Stegle, O. Spatialde: identification of spatially variable genes. Nature methods 15, 343–346 (2018).

[20] Velten, B. et al. Identifying temporal and spatial patterns of variation from multimodal data using mefisto. Nature methods 19, 179–186 (2022).

[21] Townes, F. W. & Engelhardt, B. E. Nonnegative spatial factorization applied to spatial genomics. Nature Methods 20, 229–238 (2023).

[22] Chidester, B., Zhou, T., Alam, S. & Ma, J. Spicemix enables integrative single-cell spatial modeling of cell identity. Nature genetics 55, 78–88 (2023).

[23] Dong, K. & Zhang, S. Deciphering spatial domains from spatially resolved transcriptomics with an adaptive graph attention auto-encoder. Nature communications 13, 1739 (2022).

[24] Long, Y. et al. Spatially informed clustering, integration, and deconvolution of spatial transcriptomics with graphst. Nature Communications 14, 1155 (2023).

[25] Haviv, D. et al. The covariance environment defines cellular niches for spatial inference. Nature Biotechnology 1–12 (2024).

[26] Varrone, M., Tavernari, D., Santamaria-Martínez, A., Walsh, L. A. & Ciriello, G. Cellcharter reveals spatial cell niches associated with tissue remodeling and cell plasticity. Nature Genetics 56, 74–84 (2024).

[27] Dong, M. & Kluger, Y. Geass: Neural causal feature selection for high-dimensional biological data. The Eleventh International Conference on Learning Representations (2023).

[28] Wu, Z. et al. Graph deep learning for the characterization of tumour microenvironments from spatial protein profiles in tissue specimens. Nature Biomedical Engineering 6, 1435–1448 (2022).

[29] Shimonov, S. et al. Sorbet: Automated cell-neighborhood analysis of spatial transcriptomics or proteomics for interpretable sample classification via gnn. bioRxiv 2023–12 (2024).

[30] Fischer, D. S., Schaar, A. C. & Theis, F. J. Modeling intercellular communication in tissues using spatial graphs of cells. Nature Biotechnology 41, 332–336 (2023).

[31] Arnol, D., Schapiro, D., Bodenmiller, B., Saez-Rodriguez, J. & Stegle, O. Modeling cell-cell interactions from spatial molecular data with spatial variance component analysis. Cell reports 29, 202–211 (2019).

[32] Tanevski, J., Flores, R. O. R., Gabor, A., Schapiro, D. & Saez-Rodriguez, J. Explainable multiview framework for dissecting spatial relationships from highly multiplexed data. Genome biology 23, 97 (2022).

[33] Kingma, D. P. & Welling, M. Auto-encoding variational bayes. arXiv preprint arXiv:1312.6114 (2013).

[34] Lopez, R., Regier, J., Cole, M. B., Jordan, M. I. & Yosef, N. Deep generative modeling for single-cell transcriptomics. Nature methods 15, 1053–1058 (2018).

[35] Chernozhukov, V. et al. Double/debiased machine learning for treatment and structural parameters (2018).

[36] Yao, L. et al. A survey on causal inference. ACM Transactions on Knowledge Discovery from Data (TKDD) 15, 1–46 (2021).

[37] Imbens, G. W. & Rubin, D. B. Causal inference in statistics, social, and biomedical sciences (Cambridge university press, 2015).

[38] Korsunsky, I. et al. Fast, sensitive and accurate integration of single-cell data with harmony. Nature methods 16, 1289–1296 (2019).

[39] Mørup, M. & Hansen, L. K. Archetypal analysis for machine learning and data mining. Neurocomputing 80, 54–63 (2012).

[40] Clavreul, S., Dumas, L. & Loulier, K. Astrocyte development in the cerebral cortex: Complexity of their origin, genesis, and maturation. Frontiers in neuroscience 16, 916055 (2022).

[41] Hevner, R. F. Layer-specific markers as probes for neuron type identity in human neocortex and malformations of cortical development. Journal of Neuropathology & Experimental Neurology 66, 101–109 (2007).

[42] Cable, D. M. et al. Cell type-specific inference of differential expression in spatial transcriptomics. Nature methods 19, 1076–1087 (2022).

[43] Allen, C. D., Okada, T. & Cyster, J. G. Germinal-center organization and cellular dynamics. Immunity 27, 190–202 (2007).

[44] Mesin, L., Ersching, J. & Victora, G. D. Germinal center b cell dynamics. Immunity 45, 471–482 (2016).

[45] Siegel, R. L., Miller, K. D., Wagle, N. S. & Jemal, A. Cancer statistics, 2023. Ca Cancer J Clin 73, 17–48 (2023).

[46] Pichler, A. C. et al. Tcr-independent cd137 (4-1bb) signaling promotes cd8+-exhausted t cell proliferation and terminal differentiation. Immunity (2023).

[47] Li, Q. et al. Slc6a8-mediated intracellular creatine accumulation enhances hypoxic breast cancer cell survival via ameliorating oxidative stress. Journal of Experimental & Clinical Cancer Research 40, 168 (2021).

[48] Wong, P. F. et al. Multiplex quantitative analysis of tumor-infiltrating lymphocytes and immunotherapy outcome in metastatic melanoma. Clinical Cancer Research 25, 2442–2449 (2019).

[49] Ma, R.-Y., Black, A. & Qian, B.-Z. Macrophage diversity in cancer revisited in the era of single-cell omics. Trends in immunology (2022).

[50] Christofides, A. et al. The complex role of tumor-infiltrating macrophages. Nature immunology 23, 1148–1156 (2022).

[51] Biancalani, T. et al. Deep learning and alignment of spatially resolved single-cell transcriptomes with tangram. Nature methods 18, 1352–1362 (2021).

[52] Ma, Y. & Zhou, X. Spatially informed cell-type deconvolution for spatial transcriptomics. Nature biotechnology 40, 1349–1359 (2022).

[53] Lopez, R. et al. Destvi identifies continuums of cell types in spatial transcriptomics data. Nature biotechnology 40, 1360–1369 (2022).

[54] Bergenstråhle, L. et al. Super-resolved spatial transcriptomics by deep data fusion. Nature biotechnology 40, 476–479 (2022).

[55] Hu, J. et al. Deciphering tumor ecosystems at super resolution from spatial transcriptomics with tesla. Cell systems 14, 404–417 (2023).

[56] Molnar, C. Interpretable machine learning (Lulu. com, 2020).

[57] Lotfollahi, M. et al. Mapping single-cell data to reference atlases by transfer learning. Nature biotechnology 40, 121–130 (2022).

[58] Vento-Tormo, R. et al. Single-cell reconstruction of the early maternal–fetal interface in humans. Nature 563, 347–353 (2018).

[59] Efremova, M., Vento-Tormo, M., Teichmann, S. A. & Vento-Tormo, R. Cellphonedb: inferring cell–cell communication from combined expression of multi-subunit ligand–receptor complexes. Nature protocols 15, 1484–1506 (2020).

[60] Garcia-Alonso, L. et al. Mapping the temporal and spatial dynamics of the human endometrium in vivo and in vitro. Nature genetics 53, 1698–1711 (2021).

[61] Garcia-Alonso, L. et al. Single-cell roadmap of human gonadal development. Nature 607, 540–547 (2022).

[62] Shao, X. et al. Celltalkdb: a manually curated database of ligand–receptor interactions in humans and mice. Briefings in bioinformatics 22, bbaa269 (2021).

[63] Gayoso, A. et al. A python library for probabilistic analysis of single-cell omics data. Nature biotechnology 40, 163–166 (2022).

[64] Brody, S., Alon, U. & Yahav, E. How attentive are graph attention networks? arXiv preprint arXiv:2105.14491 (2021).

[65] Rombach, R., Blattmann, A., Lorenz, D., Esser, P. & Ommer, B. High-resolution image synthesis with latent diffusion models 10684–10695 (2022).

[66] Zhao, S., Song, J. & Ermon, S. Infovae: Balancing learning and inference in variational autoencoders 33, 5885–5892 (2019).

[67] Xu, C. et al. Probabilistic harmonization and annotation of single-cell transcriptomics data with deep generative models. Molecular systems biology 17, e9620 (2021).

[68] Weinberger, E., Lin, C. & Lee, S.-I. Isolating salient variations of interest in single-cell data with contrastivevi. bioRxiv 2021–12 (2021).

[69] Kipf, T. N. & Welling, M. Semi-supervised classification with graph convolutional networks. arXiv preprint arXiv:1609.02907 (2016).

[70] Vincent, P., Larochelle, H., Bengio, Y. & Manzagol, P.-A. Extracting and composing robust features with denoising autoencoders 1096–1103 (2008).

[71] Papalexi, E. et al. Characterizing the molecular regulation of inhibitory immune checkpoints with multimodal single-cell screens. Nature genetics 53, 322–331 (2021).

[72] Dong, M. et al. Causal identification of single-cell experimental perturbation effects with cinema-ot. Nature Methods 20, 1769–1779 (2023).

[73] Luecken, M. D. et al. Benchmarking atlas-level data integration in single-cell genomics. Nature methods 19, 41–50 (2022).

[74] Xie, Z. et al. Gene set knowledge discovery with enrichr. Current protocols 1, e90 (2021).

[75] Fang, Z., Liu, X. & Peltz, G. Gseapy: a comprehensive package for performing gene set enrichment analysis in python. Bioinformatics 39, btac757 (2023).

[76] Palla, G. et al. Squidpy: a scalable framework for spatial omics analysis. Nature methods 19, 171–178 (2022).

[77] Wang, Q. et al. The allen mouse brain common coordinate framework: a 3d reference atlas. Cell 181, 936–953 (2020).

[78] Burkhardt, D. B. et al. Quantifying the effect of experimental perturbations at single-cell resolution. Nature biotechnology 39, 619–629 (2021).

[79] Martin, F. J. et al. Ensembl 2023. Nucleic acids research 51, D933–D941 (2023).

[80] Frankish, A. et al. Gencode reference annotation for the human and mouse genomes. Nucleic acids research 47, D766–D773 (2019).

[81] Fornes, O. et al. Jaspar 2020: update of the open-access database of transcription factor binding profiles. Nucleic acids research 48, D87–D92 (2020).

[82] Schep, A. N., Wu, B., Buenrostro, J. D. & Greenleaf, W. J. chromvar: inferring transcription-factor-associated accessibility from single-cell epigenomic data. Nature methods 14, 975–978 (2017).

