## Supplementary Information for "SIMVI reveals intrinsic and spatial-induced states in spatial omics data"

### Contents

**I Supplementary notes** 3

**1 Theoretical support for the SIMVI model** 3
- 1.1 Problem setting 3
- 1.2 Assumptions 5
- 1.3 Main theoretical result 6
- 1.4 Connections between the theoretical support and SIMVI design 16

**2 Spatial effect estimation** 17
- 2.1 Transforming spatial variation into archetypes 17
- 2.2 Estimating spatial effect via double machine learning 18
- 2.3 The underlying assumptions for spatial effect estimation and the positivity index 19

**3 Discussions on hyperparameter selection** 19

**II Supplementary Figures** 22

- Supplementary Figure 1 22
- Supplementary Figure 2 23
- Supplementary Figure 3 24
- Supplementary Figure 4 25
- Supplementary Figure 5 26
- Supplementary Figure 6 27
- Supplementary Figure 7 28
- Supplementary Figure 8 29
- Supplementary Figure 9 30
- Supplementary Figure 10 31
- Supplementary Figure 11 32
- Supplementary Figure 12 33
- Supplementary Figure 13 34
- Supplementary Figure 14 35
- Supplementary Figure 15 36
- Supplementary Figure 16 37
- Supplementary Figure 17 38
- Supplementary Figure 18 39
- Supplementary Figure 19 40
- Supplementary Figure 20 41
- Supplementary Figure 21 42
- Supplementary Figure 22 43

### Supplementary notes

#### 1 Theoretical support for the SIMVI model

In this part, we show that under suitable assumptions, it is possible to disentangle intrinsic and spatial-interaction induced variations in spatial omics data. Specifically, there exists a (smooth and locally injective) function mapping inferred intrinsic variations to ground truth intrinsic variations. Additionally, there exists an invertible linear transformation between ground truth spatial variations and inferred spatial variations. This serves as a theoretical basis for the SIMVI model. We state our problem setting as follows.

##### 1.1 Problem setting

We consider the following probabilistic model. Let $G = (V, E)$ be a graph, whose nodes $i \in V = \{1, 2, \ldots, n\}$ are cells, and $E$ contains an edge $(i, j)$ if the two cells $i, j$ are neighbors. We denote the set of neighbors for node $i$ as $N(i) \subset V$. We assume that the graph $G$ is $k$-regular ($k > 1$), meaning that each node has a constant degree $k > 1$. We denote the adjacency matrix of $G$ as $W \in \mathbb{R}^{n\times n}$.

We assume that the gene expression vector $\boldsymbol{x}^i$ for the cell at node $i$ is sampled from a distribution parametrized by two sets of latent variables, defined as follows. The first set, denoted as $\boldsymbol{z}^{\pi(i)} \in \mathbb{R}^{d_1}$, represents the intrinsic variability of cell $i$. Generation of $\boldsymbol{z}^\pi$ involves two steps to account for possible spatial proximity across cells with certain intrinsic properties. We first sample a "bag of cells" $\boldsymbol{z} = [\boldsymbol{z}^1, \boldsymbol{z}^2, \ldots, \boldsymbol{z}^n]$ with $\{\boldsymbol{z}^i\}$ i.i.d., then we generate a permutation $\pi$ and assign cell $i$ in the bag to the node $\pi(i)$ on the spatial graph. The permutation $\pi$ is generated conditioning on both $\boldsymbol{z}$ and the graph $G$. The second set of latent variables, denoted as $\boldsymbol{s}^i \in \mathbb{R}^{d_2}$, characterizes the contribution of spatial interaction. We assume that $\boldsymbol{s}^i$ depends on the intrinsic variabilities $\boldsymbol{z}^{\pi N(i)}$ of the cells which are neighbors of cell $i$. Formally, $(\boldsymbol{z}^\pi, \boldsymbol{s})$ are generated by the following procedure:

- Generation of the unordered intrinsic variation $\boldsymbol{z} = [\boldsymbol{z}^1, \boldsymbol{z}^2, \ldots, \boldsymbol{z}^n]$, where $\boldsymbol{z}^i \overset{\text{i.i.d}}{\sim} \mathcal{N}(0, I_{d_1})$.
- Generation of the spatial-intrinsic-aware permutation $\pi : [n] \to [n]$. We assume the probability of a specific spatial allocation $\pi$, is determined by a potential function $U_G : \mathbb{R}^{d_1\times n} \to \mathbb{R}_+$. The probability of the permutation $\pi$ is of form:

$$p(\pi|\boldsymbol{z}) = \frac{1}{Z} U_G(\boldsymbol{z}^\pi), \tag{1}$$

  where $Z$ is an appropriate normalization constant. After sampling of $\pi$, $\boldsymbol{z}^{\pi(i)}$ indicates the intrinsic variation for the cell at node $i$.
- Generation of the spatial variation $\boldsymbol{s}^i$ for cell $i$. It depends on the neighborhood intrinsic variation $\boldsymbol{z}^{\pi(N(i))}$, and follows a normal distribution parametrized by a matrix $\boldsymbol{A} \in \mathbb{R}^{d_2\times kd_1}$ and a fixed full rank covariance matrix $\boldsymbol{\Sigma_s} \in \mathbb{S}^{d_2}_{++}$:

$$\boldsymbol{s}^i|\boldsymbol{z} \sim \mathcal{N}(\boldsymbol{A}\boldsymbol{z}^{\pi(N(i))}, \boldsymbol{\Sigma_s}); \tag{2}$$

- Generation of the gene expression vector $\boldsymbol{x}^i$ at cell $i$:

$$\forall i, \boldsymbol{x}^i \sim NB(f(\boldsymbol{z}^{\pi(i)}, \boldsymbol{s}^i), g). \tag{3}$$

  Here, $NB$ stands for the negative binomial distribution, and $f : \mathbb{R}^{d_1} \times \mathbb{R}^{d_2} \to \mathbb{R}^p$ is a continuous smooth function representing the mean of the negative binomial distribution. The vector $g \in \mathbb{R}^p$ characterizes the dispersion of the distribution (consistent with the "gene" dispersion setting in scvi-tools [1]).

Additional properties and assumptions for the generative process are described below.

◇ The marginal density of $\boldsymbol{z}^\pi$ at a point $z^\pi \in \mathbb{R}^{d_1\times n}$ can be expressed as:

$$p(\boldsymbol{z}^\pi = z^\pi) = \int p(\boldsymbol{z}^\pi = z^\pi|\boldsymbol{z})p(\boldsymbol{z})d\boldsymbol{z} \propto U_G(z^\pi)p(\boldsymbol{z} = z^\pi). \tag{4}$$

◇ We consider a specific case of $U_G$ that corresponds to the Gaussian Markov random field. In this case, $U_G$ can be decomposed as follows:

$$U_G(z^\pi) = \prod_{(i,j)\in E} \exp(-\frac{1}{2}(z^{\pi(i)})^T \mathbf{\Omega} z^{\pi(j)}), \tag{5}$$

where $\mathbf{\Omega} \in \mathbb{R}^{d_1\times d_1}$ is a coefficient matrix representing the colocalization tendency. Thus, $\boldsymbol{z}^\pi$ follows a Gaussian distribution:

$$p(\boldsymbol{z}^\pi = z^\pi) \propto \exp(-\frac{1}{2}\sum_{(i,j)\in E}(z^{\pi(i)})^T \mathbf{\Omega} z^{\pi(j)} - \frac{1}{2}\sum_i (z^{\pi(i)})^T z^{\pi(i)}) = \exp(-\frac{1}{2}(z^\pi)^T \mathbf{\Omega}_G z^\pi), \tag{6}$$

where $\mathbf{\Omega}_G = W \otimes \mathbf{\Omega} + I$, $\otimes$ stands for Kronecker product, and $W$ denotes the adjacency matrix of $G$.

◇ We additionally assume that the marginal distribution of $(\boldsymbol{z}^{\pi(i)}, \boldsymbol{z}^{\pi(N(i))})$ is identical across all nodes. According to Eq. (6), this distribution is jointly (non-degenerate) Gaussian, which we denote as follows:

$$\begin{gathered}(\boldsymbol{z}^{\pi(i)}, \boldsymbol{z}^{\pi(N(i))}) \sim \mathcal{N}(0, \mathbf{\Sigma}); \quad \mathbf{\Sigma} \in \mathbb{S}_{++}^{d_1\times(k+1)}, \\ \mathbf{\Sigma} = \left(\begin{array}{c|c} I_{d_1} & \mathbf{\Sigma}_1^T \\ \hline \mathbf{\Sigma}_1 & \mathbf{\Sigma}_0 \end{array}\right), \ \mathbf{\Sigma}_1 \in \mathbb{R}^{kd_1\times d_1}, \mathbf{\Sigma}_0 \in \mathbb{R}^{kd_1\times kd_1}.\end{gathered} \tag{7}$$

◇ Combining Eqs. (2, 7), we may formulate an equivalent generative process for $\boldsymbol{s}^i$ that only involves $\boldsymbol{z}^{\pi(i)}$. Considering the cell at node $i$, applying Eqs. (2, 7) yields the following result based on the conditional distribution formula for Gaussians:

$$\begin{gathered}\boldsymbol{s}^i|\boldsymbol{z}^{\pi(i)} \sim \mathcal{N}(0, \mathbf{\Sigma}_{\boldsymbol{s}}) + \boldsymbol{A}\boldsymbol{\eta}; \quad \boldsymbol{\eta} \sim \mathcal{N}(\mathbf{\Sigma}_1 \boldsymbol{z}^{\pi(i)}, \mathbf{\Sigma}_0 - \mathbf{\Sigma}_1\mathbf{\Sigma}_1^T); \\ \boldsymbol{s}^i|\boldsymbol{z}^{\pi(i)} \sim \mathcal{N}(0, \mathbf{\Sigma}_{\boldsymbol{s}} + \boldsymbol{A}(\mathbf{\Sigma}_0 - \mathbf{\Sigma}_1\mathbf{\Sigma}_1^T)\boldsymbol{A}^T) + \boldsymbol{A}\mathbf{\Sigma}_1\boldsymbol{z}^{\pi(i)}.\end{gathered} \tag{8}$$

Denote

$$\boldsymbol{A}' = \boldsymbol{A}\mathbf{\Sigma}_1, \quad \mathbf{\Sigma}' = \mathbf{\Sigma}_{\boldsymbol{s}} + \boldsymbol{A}(\mathbf{\Sigma}_0 - \mathbf{\Sigma}_1\mathbf{\Sigma}_1^T)\boldsymbol{A}^T. \tag{9}$$

Then we have

$$\boldsymbol{s}^i|\boldsymbol{z}^{\pi(i)} \sim \mathcal{N}(0, \mathbf{\Sigma}') + \boldsymbol{A}'\boldsymbol{z}^{\pi(i)}. \tag{10}$$

The fact that $\mathbf{\Sigma}_{\boldsymbol{s}}$ has full rank implies that the matrix $\mathbf{\Sigma}'$ is also full rank. Now the marginal covariance of $\boldsymbol{s}^i$ is $\boldsymbol{A}'\mathbf{\Sigma}_1\mathbf{\Sigma}_1^T\boldsymbol{A}'^T + \mathbf{\Sigma}_{\boldsymbol{s}} + \boldsymbol{A}(\mathbf{\Sigma}_0 - \mathbf{\Sigma}_1\mathbf{\Sigma}_1^T)\boldsymbol{A}^T = \mathbf{\Sigma}_{\boldsymbol{s}} + \boldsymbol{A}\mathbf{\Sigma}_0\boldsymbol{A}^T$. We suppose the marginal distribution of $\boldsymbol{s}^i$ to be standard normal, i.e., $I = \mathbf{\Sigma}_{\boldsymbol{s}} + \boldsymbol{A}\mathbf{\Sigma}_0\boldsymbol{A}^T$. In this case, $\mathbf{\Sigma}_{\boldsymbol{s}}$ can be expressed by $\boldsymbol{A}, \mathbf{\Sigma}$.

◇ We assume that $f$ is injective and that its inverse is smooth, making $f$ a diffeomorphism. In particular, denote $d = d_1 + d_2$, then this implies $p \geq d$. We denote the first $d_1$ components of $f^{-1}$ as $f_z^{-1}$, and the last $d_2$ components of $f^{-1}$ as $f_s^{-1}$. To simplify the problem setting, we assume the constant vector $g$ is fixed and known.

◇ Finally, Eqs. (7,2,3) imply that the joint marginal distribution $(\boldsymbol{z}^{\pi(i)}, \boldsymbol{z}^{\pi(N(i))}, \boldsymbol{s}^i, \boldsymbol{x}^i)$ is stationary in the generative process, meaning it is identical for any node $i$ on the graph. We drop the index $i$ and denote the random variables for each cell as $(\boldsymbol{z}^\pi, \boldsymbol{z}^{\pi N}, \boldsymbol{s}, \boldsymbol{x})$ in the following text.

For the sake of convenience, we also omit the superscript $\pi$ in the notations ($\boldsymbol{z}^\pi$, etc). Henceforth, we use bold symbols $\boldsymbol{z}, \boldsymbol{s}$, etc. to denote random variables, and use $z, s$, etc. to denote values of the corresponding random variables.

We denote the range of $f(\boldsymbol{z}, \boldsymbol{s})$ as $\mathcal{P} = \{f(z,s)|(z,s) \in \mathbb{R}^d\}$. We denote by $\theta = (\mathbf{\Sigma}, \boldsymbol{A}, f)$ the parameters of the true generative model described above. Further, we denote by $p_\theta(x)$ the probability density of a gene expression vector $x \in \mathbb{R}^p$ for any individual cell. Next, we define the set of all possible parameters that yield the same distribution as $p_\theta(x)$:

**Definition 1.** (Parameter family) Let $\Theta$ be the set of all possible parameters that yield the same distribution for the gene expression of a cell:

$$\Theta = \{\tilde{\theta} := (\tilde{\boldsymbol{\Sigma}}, \tilde{\boldsymbol{A}}, \tilde{f}) \,|\, p_{\tilde{\theta}}(x) = p_\theta(x) \text{ and } \tilde{f} : \mathbb{R}^d \to \mathcal{P} \text{ is smooth and invertible with a smooth inverse}\}$$

Clearly, the ground truth parameter $\theta \in \Theta$. Each element $\tilde{\theta} \in \Theta$ defines a distribution of latent variables $(\tilde{\boldsymbol{z}}, \tilde{\boldsymbol{s}})$ generated through Eqs. (7,2) with parameters $(\tilde{\boldsymbol{\Sigma}}, \tilde{\boldsymbol{A}})$. We allow the dimension of $\tilde{\boldsymbol{z}}, \tilde{\boldsymbol{s}}$ to be misspecified (i.e. $\tilde{d}_1 \neq d_1$, $\tilde{d}_2 \neq d_2$). Nevertheless, because $\tilde{f}$ defines a bijective diffeomorphism from $\mathbb{R}^{\tilde{d}_1+\tilde{d}_2}$ to the gene expression space, the composition $\tilde{f}^{-1} \circ f$ defines a bijective diffeomorphism from $\mathbb{R}^{d_1+d_2}$ to $\mathbb{R}^{\tilde{d}_1+\tilde{d}_2}$. Therefore, we have that $\tilde{\theta} \in \Theta \Rightarrow \tilde{d} := \tilde{d}_1 + \tilde{d}_2 = d$.

Our goal here is to prove that the parameters of the generative process can be uniquely identified from the distribution $p(x)$, i.e. model identifiability. Intuitively, we aim to prove that, if one model parameter $(\tilde{\boldsymbol{\Sigma}}, \tilde{\boldsymbol{A}}, \tilde{f}) \in \Theta$ gives rise to the same distribution $p(x)$ as that generated by ground truth $(\boldsymbol{\Sigma}, \boldsymbol{A}, f)$, then $(\tilde{\boldsymbol{\Sigma}}, \tilde{\boldsymbol{A}}, \tilde{f})$ is "equivalent" to $(\boldsymbol{\Sigma}, \boldsymbol{A}, f)$ with respect to an equivalence relationship $\sim$. Formally, we define the specific equivalence relationship of interest, and model identifiability as follows.

Given our primary objective of disentangling different sources of variation, we introduce a novel form of equivalence tailored to our purpose, which we term "disentanglement equivalence":

**Definition 2.** (Disentanglement equivalence) Let $\tilde{\theta} = (\tilde{\boldsymbol{\Sigma}}, \tilde{\boldsymbol{A}}, \tilde{f}) \in \Theta$ be a set of parameters yielding the same distribution $p(x)$ as the ground truth $\theta = (\boldsymbol{\Sigma}, \boldsymbol{A}, f)$. We say that $\tilde{\theta}$ satisfies the disentanglement equivalence relationship w.r.t. $\theta$, denoted as $\tilde{\theta} \sim \theta$, if the following three conditions hold:

$$\text{(i):} \quad \tilde{d}_1 = d_1, \tilde{d}_2 = d_2; \tag{11}$$

$$\text{(ii):} \quad \exists \text{ function } F : \mathbb{R}^{d_1} \to \mathbb{R}^{d_1},\ s.t.\ \forall \rho \in \mathcal{P},\ f_z^{-1}(\rho) = F(\tilde{f}_z^{-1}(\rho)); \tag{12}$$

$$\text{(iii):} \quad \exists \text{ invertible } L \in \mathbb{R}^{d_2 \times d_2}, c \in \mathbb{R}^{d_2},\ s.t.\ f_s^{-1}(\rho) = L\tilde{f}_s^{-1}(\rho) + c. \tag{13}$$

This means that, for any $\rho$, its corresponding ground truth intrinsic variation $f_z^{-1}(\rho)$ is a nonlinear mapping of the inferred intrinsic variation $\tilde{f}_z^{-1}(\rho)$, whereas its ground truth spatial variation $f_s^{-1}(\rho)$ is a linear transformation of the inferred spatial variation $\tilde{f}_s^{-1}(\rho)$.

We next define the model identifiability with respect to the disentanglement equivalence relationship $\sim$. Our definition of model identifiability mostly follows prior works on VAE identifiability [2].

**Definition 3.** (Model identifiability) The ground truth model parameter $\theta = (\boldsymbol{\Sigma}, \boldsymbol{A}, f)$ is $\sim$-identifiable on $\Theta$ if $\forall \tilde{\theta} \in \Theta$, $\tilde{\theta} \sim \theta$ holds.

*Remark* 1. Our theoretical analysis considers the true distribution $p_\theta(x)$. In practice, we can only observe the empirical distribution over a finite number of cells. Moreover, these cells exhibit dependencies within their local neighborhood. Unlike in i.i.d. settings, the strong convergence of this empirical distribution to the true distribution needs additional assumptions to hold, such as ergodicity or strong mixing. These assumptions describe the diminishing influence of individual cells over space, which intuitively holds true for cells in spatial omics datasets. Therefore, we expect the empirical distribution to closely approximate the true one when the number of cells $n \gg 1$.

*Remark* 2. The SIMVI model provides two options for modeling $p(x)$: the Zero-Inflated Negative Binomial (ZINB) and the Negative Binomial (NB) distributions, with ZINB more commonly used. However, these two options introduce only technical differences in the theoretical analysis. Consequently, we focus on the NB case in the theory section.

We next state the assumptions. In the following text, $(\boldsymbol{\Sigma}, \boldsymbol{A}, f)$ refers to the ground truth solution, a specific instance in the parameter set $\Theta$. $(\tilde{\boldsymbol{\Sigma}}, \tilde{\boldsymbol{A}}, \tilde{f})$ refers to **any parameters** within the parameter set $\Theta$.

#### 1.2 Assumptions

**Assumption 1.** *(Non-exploding moments) We assume that the ground truth function $f$ satisfies the element-wise property: $\forall j \in \{1, 2, \ldots, p\}$, $\mathbb{E}f_j^{2m}(\boldsymbol{z}, \boldsymbol{s}) = \mathcal{O}((2m)^{2m})$ for $m \in \mathbb{N}$.*

*Remark* 3. This assumption leads to two direct consequences (via Holder inequality). First, $\forall j \in \{1, 2, \ldots, p\}$, $\mathbb{E}f_j^m(\boldsymbol{z}, \boldsymbol{s}) < \infty$ for any $m \in \mathbb{N}_0$. Second, for a non-negative integer vector $\boldsymbol{m} \in \mathbb{N}_0^p$, we denote $f^{\boldsymbol{m}}(\boldsymbol{z}, \boldsymbol{s}) := \prod_{j=1}^p f_j^{\boldsymbol{m}_j}(\boldsymbol{z}, \boldsymbol{s})$. Then we have that $\forall \boldsymbol{m}, \mathbb{E}f^{\boldsymbol{m}}(\boldsymbol{z}, \boldsymbol{s}) < \infty$.

**Assumption 2.** *(Sufficient variability) We assume that the dimensionality $d_1$ of the intrinsic latent variable $\boldsymbol{z}$, and the dimensionality $d_2$ of the spatial latent variable $\boldsymbol{s}$ satisfy $d_1 \geq d_2$. Also we assume that the ground truth matrix $\boldsymbol{A}' = \boldsymbol{A}\boldsymbol{\Sigma}_1 \in \mathbb{R}^{d_2 \times d_1}$ is of full rank $d_2$.*

*Remark* 4. The matrix $\boldsymbol{A}'$ indicates the underlying contribution of the intrinsic variation on the spatial variation. The assumption on the rank of $\boldsymbol{A}'$ plays a key role in establishing the linear identifiability of $\boldsymbol{s}$ (Eq. (13)). Intuitively, this condition is met when the intrinsic variation of a cell $\boldsymbol{z}^i$ is informative of its neighborhood $\boldsymbol{z}^{N(i)}$. This property is typically satisfied in real spatial omics datasets. If not, a non-linear identifiability result for $\boldsymbol{s}$ still holds, as shown by Lemma 3.

**Assumption 3.** *(Spatial / intrinsic variation covers all possible states) For any $(z, s) \in \mathbb{R}^d$ denote $\rho = f(z, s)$. We assume that for any $(\tilde{\boldsymbol{\Sigma}}, \tilde{\boldsymbol{A}}, \tilde{f}) \in \Theta$, the following two conditions hold. (i). The range of $\tilde{f}_s^{-1}(\rho)$ is $\mathbb{R}^{\tilde{d}_2}$ for any fixed $z$ and $\tilde{f}_z^{-1}(\rho)$; (ii). The range of $z$ is $\mathbb{R}^{d_1}$ for any fixed $s$ and $\tilde{f}_s^{-1}(\rho)$.*

*Remark* 5. Assumption 3 implies the following two properties for any model parameter $\tilde{\theta}$: 1. the "inferred spatial variation" $\tilde{f}_s^{-1}(\rho)$ covers the full space for cells with any ground truth intrinsic property $z$ and "inferred intrinsic variation" $\tilde{f}_z^{-1}(\rho)$; 2. the intrinsic property of $z$ covers the full space for cells with any ground truth spatial property $s$ and "inferred spatial variation" $\tilde{f}_s^{-1}(\rho)$. The assumption is automatically satisfied for ground truth $\theta$. This is enforced in the SIMVI design as we approximate $\tilde{f}_s^{-1}$ by the variational posterior of $\boldsymbol{x}^N$ instead of $\boldsymbol{x}$, and enforce the joint distribution of $(\tilde{f}_z^{-1}(\boldsymbol{\rho}), \tilde{f}_s^{-1}(\boldsymbol{\rho}))$ to be standard normal.

**Assumption 4.** *(Minimal information in the intrinsic variation) We assume that the dimensionality $\tilde{d}_1$ for any $\tilde{\theta} \in \Theta$ is not larger than the ground truth $d_1$: $\forall \tilde{\theta} \in \Theta,\ \tilde{d}_1 \leq d_1$.*

*Remark* 6. Assumption 4 addresses the assymmetric independence regularization term in the SIMVI model in our theoretical setting. The assymmetric independence regularization term constrains the "total information" encoded by $\tilde{\boldsymbol{z}}$. In the theoretical context, the objective aligns with minimizing the dimensionality of $\tilde{\boldsymbol{z}}$.

#### 1.3 Main theoretical result

Prior to the statement and proof of the main theorems, we first provide a number of lemmas that will be used in the proof.

**Lemma 1.** *Suppose $\theta, \tilde{\theta} \in \Theta$ are two parameters that result in the same distribution of $\boldsymbol{x}$. We denote the latent variables generated with $\theta, \tilde{\theta}$ using Eqs. (7,2) as $(\boldsymbol{z}, \boldsymbol{s})$ and $(\tilde{\boldsymbol{z}}, \tilde{\boldsymbol{s}})$ respectively. Then under assumption 1, $f(\boldsymbol{z}, \boldsymbol{s}), \tilde{f}(\tilde{\boldsymbol{z}}, \tilde{\boldsymbol{s}})$ have the same distribution:*

$$p_\theta(x) = p_{\tilde{\theta}}(x) \Rightarrow f(\boldsymbol{z}, \boldsymbol{s}) \overset{d}{=} \tilde{f}(\tilde{\boldsymbol{z}}, \tilde{\boldsymbol{s}}). \tag{14}$$

*Proof.* For a non-negative integer vector $\boldsymbol{m} \in \mathbb{N}_0^p$, we denote $\boldsymbol{x}^{\boldsymbol{m}} := \prod_{j=1}^p \boldsymbol{x}_j^{\boldsymbol{m}_j}$. Since $p_\theta(x) = p_{\tilde{\theta}}(x)$, we have $\forall \boldsymbol{m}, \mathbb{E}_\theta(\boldsymbol{x}^{\boldsymbol{m}}) = \mathbb{E}_{\tilde{\theta}}(\boldsymbol{x}^{\boldsymbol{m}})$. Applying the law of total probability gives

$$\forall \boldsymbol{m}, \int \mathbb{E}(\boldsymbol{x}^{\boldsymbol{m}}|f(z,s))p_\theta(z,s)dzds = \int \mathbb{E}(\boldsymbol{x}^{\boldsymbol{m}}|\tilde{f}(\tilde{z},\tilde{s}))p_{\tilde{\theta}}(\tilde{z},\tilde{s})d\tilde{z}d\tilde{s}. \tag{15}$$

Here $p_\theta(z, s)$ and $p_{\tilde{\theta}}(\tilde{z}, \tilde{s})$ denote the density of $(\boldsymbol{z}, \boldsymbol{s})$ and $(\tilde{\boldsymbol{z}}, \tilde{\boldsymbol{s}})$ respectively. Now we use Eq. (15) to prove the following result regarding the multivariate moments by induction:

$$\forall \boldsymbol{m} \in \mathbb{N}_0^p, \int f^{\boldsymbol{m}}(z,s)p_\theta(z,s)dzds = \int \tilde{f}^{\boldsymbol{m}}(\tilde{z},\tilde{s})p_{\tilde{\theta}}(\tilde{z},\tilde{s})d\tilde{z}d\tilde{s} \tag{16}$$

In the following proof, we use the parameterization of the NB distribution using the success probability vector $\boldsymbol{\alpha} \in \mathbb{R}^p$ and number of successes $\boldsymbol{r} \in \mathbb{N}_0^p$. There is a simple correspondence between the $\alpha$-$r$ parametrization and the mean-dispersion parametrization. Namely for the expectation term, $f(\boldsymbol{z}, \boldsymbol{s}) = \boldsymbol{r}(1 - \boldsymbol{\alpha})/\boldsymbol{\alpha}$; for the dispersion term, $g = 1/\boldsymbol{r}$.

To start with, we derive the $m$-th raw moment of a univariate NB distribution with parameters $(r, \alpha)$. Recall that the moment generating function (MGF) of the NB$(r, \alpha)$ distribution is given by $\psi(t) = (\frac{1-e^t}{\alpha} + e^t)^{-r}$. Thus, the $m$-th moment can be expressed as follows by Faà di Bruno's formula (section 3.4, Theorem A of [3]):

$$\begin{aligned}\frac{d^m}{dt^m}\psi(t)|_{t=0} =& \frac{d^m}{dt^m}[(\frac{1-e^t}{\alpha} + e^t)^{-r}]|_{t=0} \\ =& \sum_{k=0}^{m}(x^{-r})^{(k)}|_{x=\frac{1-e^t}{\alpha}+e^t} \cdot B_{m,k}\left((\frac{1-e^t}{\alpha} + e^t)^{(1)}, \dots, (\frac{1-e^t}{\alpha} + e^t)^{(m-k+1)}\right)|_{t=0}.\end{aligned} \tag{17}$$

Here $B_{m,k}$ stands for the Bell polynomial (also introduced in section 3.4, Theorem A of [3]):

$$B_{m,k}(x_1, x_2, \dots, x_{m-k+1}) = \sum_{\substack{j_1+j_2+\dots+j_{m-k+1}=k, \\ j_1+2j_2+\dots+(m-k+1)j_{m-k+1}=m}} \frac{m!}{j_1!j_2!\cdots j_{m-k+1}!}\left(\frac{x_1}{1!}\right)^{j_1}\left(\frac{x_2}{2!}\right)^{j_2}\cdots\left(\frac{x_{m-k+1}}{(m-k+1)!}\right)^{j_{m-k+1}}. \tag{18}$$

We next simplify Eq. (17) by the following three equalities. The final equality follows from the definition of the Bell polynomial (18).

$$(\frac{1-e^t}{\alpha} + e^t)|_{t=0} = 1; \qquad \forall m \geq 1,\ (\frac{1-e^t}{\alpha} + e^t)^{(m)}|_{t=0} = 1 - \frac{1}{\alpha};$$

$$B_{m,k}(x, x, \dots, x) = c_{m,k}x^k,\ \text{where } c_{m,k} \text{ is a positive constant determined by } m, k.$$

Leveraging these equalities and that $(x^{-r})^{(k)}|_{x=1} = \prod_{l=-r-k+1}^{-r} l$, we have that

$$\begin{aligned}\frac{d^m}{dt^m}[(\frac{1-e^t}{\alpha} + e^t)^{-r}]|_{t=0} =& \sum_{k=0}^{m}(x^{-r})^{(k)}|_{x=1} \cdot B_{m,k}\left(1 - \frac{1}{\alpha}, \dots, 1 - \frac{1}{\alpha}\right) \\ =& \sum_{k=0}^{m}\left(\prod_{l=-r-k+1}^{-r} l\right)c_{m,k}(-\frac{1}{r})^k(r\frac{1-\alpha}{\alpha})^k.\end{aligned} \tag{19}$$

In summary, the $m$-th moment can be expressed by a polynomial with respect to $r\frac{1-\alpha}{\alpha}$. The polynomial is of degree $m$, as its leading term (of order $m$) has a non-zero coefficient given by $\left(\prod_{l=-r-m+1}^{-r} l\right)c_{m,m}(-1/r)^m$. We denote the coefficients $q_{m,k}(r) = \left(\prod_{l=-r-k+1}^{-r} l\right)c_{m,k}(-\frac{1}{r})^k$ and rewrite the polynomial as:

$$\text{The } m\text{-th moment of NB}(r, \alpha) = \sum_{k=0}^{m} q_{m,k}(r)(r\frac{1-\alpha}{\alpha})^k,\ q_{m,m}(r) \neq 0.$$

Inserting $m = \boldsymbol{m}_j$, $r = \boldsymbol{r}_j = 1/g_j$ and $r(1-\alpha)/\alpha = f_j(\boldsymbol{z}, \boldsymbol{s})$ in the above equation leads to

$$\mathbb{E}(\boldsymbol{x}_j^{\boldsymbol{m}_j}|f(z,s)) = \sum_{k=0}^{\boldsymbol{m}_j} q_{\boldsymbol{m}_j,k}(\frac{1}{g_j})f_j^k(\boldsymbol{z}, \boldsymbol{s}),\ q_{\boldsymbol{m}_j,\boldsymbol{m}_j}(\frac{1}{g_j}) \neq 0, \tag{20}$$

with $\mathbb{E}(\boldsymbol{x}_j^{\boldsymbol{m}_j}|\tilde{f}(\tilde{z}, \tilde{s}))$ admitting a similar expansion. Now we prove that Eq. (16) holds by induction, leveraging Eqs. (15,20). We first note that the expectation terms $\mathbb{E}(\boldsymbol{x}^{\boldsymbol{m}}|f(z,s)), \mathbb{E}(\boldsymbol{x}^{\boldsymbol{m}}|\tilde{f}(\tilde{z}, \tilde{s}))$ can be factorized due to the conditional independence of their $p$ components:

$$\mathbb{E}(\boldsymbol{x}^{\boldsymbol{m}}|f(z,s)) = \prod_j \mathbb{E}(\boldsymbol{x}_j^{\boldsymbol{m}_j}|f(z,s)), \quad \mathbb{E}(\boldsymbol{x}^{\boldsymbol{m}}|\tilde{f}(\tilde{z}, \tilde{s})) = \prod_j \mathbb{E}(\boldsymbol{x}_j^{\boldsymbol{m}_j}|\tilde{f}(\tilde{z}, \tilde{s})). \tag{21}$$

Therefore plugging $\boldsymbol{m} \in \{0,1\}^p$ in Eq. (15) directly gives that for all $\boldsymbol{m}$ with $\sum_j \boldsymbol{m}_j \leq 1$,

$$\int f^{\boldsymbol{m}}(z,s)p_\theta(z,s)dzds = \int \tilde{f}^{\boldsymbol{m}}(\tilde{z},\tilde{s})p_{\tilde{\theta}}(\tilde{z},\tilde{s})d\tilde{z}d\tilde{s}. \tag{22}$$

Next we prove the induction step, that if Eq. (16) holds for all $\boldsymbol{m}$ with $\sum_j \boldsymbol{m}_j \leq n$, then it also holds for $\boldsymbol{m}$ with $\sum_j \boldsymbol{m}_j = n+1$. Plugging the factorization (21) and the expression (20) in Eq. (15) yields

$$\int \prod_{j=1}^{p}\left(\sum_{k=0}^{\boldsymbol{m}_j} q_{\boldsymbol{m}_j,k}(\frac{1}{g_j})f_j^k(z,s)\right)p_\theta(z,s)dzds = \int \prod_{j=1}^{p}\left(\sum_{k=0}^{\boldsymbol{m}_j} q_{\boldsymbol{m}_j,k}(\frac{1}{g_j})\tilde{f}_j^k(\tilde{z},\tilde{s})\right)p_{\tilde{\theta}}(\tilde{z},\tilde{s})d\tilde{z}d\tilde{s} \tag{23}$$

Note that all coefficients of form $q_{\boldsymbol{m}_j,k}(\frac{1}{g_j})$ are identical on both sides. Expanding Eq. (23) on both sides, we have

$$\sum_{\boldsymbol{m}'}\int \prod_{j=1}^{p}\left(q_{\boldsymbol{m}_j,\boldsymbol{m}'_j}(\frac{1}{g_j})f_j^{\boldsymbol{m}'_j}(z,s)\right)p_\theta(z,s)dzds = \sum_{\boldsymbol{m}'}\int \prod_{j=1}^{p}\left(q_{\boldsymbol{m}_j,\boldsymbol{m}'_j}(\frac{1}{g_j})\tilde{f}_j^{\boldsymbol{m}'_j}(\tilde{z},\tilde{s})\right)p_{\tilde{\theta}}(\tilde{z},\tilde{s})d\tilde{z}d\tilde{s}.$$

Here $\boldsymbol{m}' \in \mathbb{N}_0^p$ are all vectors that satisfy $\boldsymbol{m}'_j \leq \boldsymbol{m}_j$ for all $j \in \{1,2,\ldots,p\}$. Apart from the sole $\boldsymbol{m}' = \boldsymbol{m}$, all other $\boldsymbol{m}'$ satisfy $\sum_j \boldsymbol{m}'_j \leq n$. By induction, we can cancel these terms, leaving only the term with the highest degree $\boldsymbol{m}$, simplifying the equation to

$$\begin{aligned}
&\int \left(\prod_{j=1}^{p} q_{\boldsymbol{m}_j,\boldsymbol{m}_j}(\frac{1}{g_j})f_j^{\boldsymbol{m}_j}(\boldsymbol{z},\boldsymbol{s})\right)p_\theta(z,s)dzds = \int \left(\prod_{j=1}^{p} q_{\boldsymbol{m}_j,\boldsymbol{m}_j}(\frac{1}{g_j})\tilde{f}_j^{\boldsymbol{m}_j}(\tilde{\boldsymbol{z}},\tilde{\boldsymbol{s}})\right)p_{\tilde{\theta}}(\tilde{z},\tilde{s})d\tilde{z}d\tilde{s} \\
\Rightarrow &\int \left(\prod_{j=1}^{p} q_{\boldsymbol{m}_j,\boldsymbol{m}_j}(\frac{1}{g_j})\right) f^{\boldsymbol{m}}(\boldsymbol{z},\boldsymbol{s})p_\theta(z,s)dzds = \int \left(\prod_{j=1}^{p} q_{\boldsymbol{m}_j,\boldsymbol{m}_j}(\frac{1}{g_j})\right)\tilde{f}^{\boldsymbol{m}}(\tilde{\boldsymbol{z}},\tilde{\boldsymbol{s}})p_{\tilde{\theta}}(\tilde{z},\tilde{s})d\tilde{z}d\tilde{s}.
\end{aligned} \tag{24}$$

Finally, by Eq. (20), $\left(\prod_{j=1}^{p} q_{\boldsymbol{m}_j,\boldsymbol{m}_j}(\frac{1}{g_j})\right)$ is not zero and can be canceled on both sides. Thus the induction is complete. That is, Eq. (16) holds for all $\boldsymbol{m} \in \mathbb{N}_0^p$.

Eq. (16) indicates that $f(\boldsymbol{z},\boldsymbol{s})$ and $\tilde{f}(\tilde{\boldsymbol{z}},\tilde{\boldsymbol{s}})$ have identical multivariate moments for any non-negative order:

$$\forall \boldsymbol{m} \in \mathbb{N}_0^p, \quad \mathbb{E}f^{\boldsymbol{m}}(\boldsymbol{z},\boldsymbol{s}) = \mathbb{E}\tilde{f}^{\boldsymbol{m}}(\tilde{\boldsymbol{z}},\tilde{\boldsymbol{s}}). \tag{25}$$

Now we show that the distributions $f^{\boldsymbol{m}}(\boldsymbol{z},\boldsymbol{s}), \tilde{f}^{\boldsymbol{m}}(\tilde{\boldsymbol{z}},\tilde{\boldsymbol{s}})$ are determined by their non-negative moments (M-det), i.e. Eq. (25) implies $f(\boldsymbol{z},\boldsymbol{s}) \stackrel{d}{=} \tilde{f}(\tilde{\boldsymbol{z}},\tilde{\boldsymbol{s}})$. This is a well-known problem in probability theory called "the moment problem" [4, 5, 6], and we leverage established results for the problem to show the desired results. The theorem 3 and 4 in [5] (formulated as Theorem 2 in [6]) states that, $f, \tilde{f}$ are M-det if all of their marginals $f_j, \tilde{f}_j$ are M-det. [4] provides a list of assumptions for each marginal to be M-det, including our Assumption 1 for $f_j$. We further note that, by Eq. (25), Assumption 1 also holds for $\tilde{f}_j$. Together, we have that both $f^{\boldsymbol{m}}(\boldsymbol{z},\boldsymbol{s}), \tilde{f}^{\boldsymbol{m}}(\tilde{\boldsymbol{z}},\tilde{\boldsymbol{s}})$ are M-det. Therefore by Eq. (25), the desired relation Eq. (14) holds. □

**Lemma 2. (Functional dependence on inferred and ground truth intrinsic variation)** *Under assumptions 3, 4,* $\forall(\tilde{\boldsymbol{\Sigma}}, \tilde{\boldsymbol{A}}, \tilde{f}) \in \Theta$, *there exists a function* $F : \mathbb{R}^{\tilde{d}_1} \to \mathbb{R}^{d_1}$, *such that*

$$f_z^{-1}(\rho) = F(\tilde{f}_z^{-1}(\rho)), \quad \forall \rho \in \mathcal{P}. \tag{26}$$

*This means that for any* $\rho$, *its corresponding ground truth intrinsic variation* $f_z^{-1}(\rho)$ *equals a nonlinear mapping of the inferred intrinsic variation* $\tilde{f}_z^{-1}(\rho)$. *Furthermore, we have* $\tilde{d}_1 = d_1$ *and* $\tilde{d}_2 = d_2$. *Together, (i) and (ii) in Definition 2 holds for any* $\tilde{\theta} \in \Theta$.

*Proof.* Let $(z, s) \in (\mathbb{R}^{d_1}, \mathbb{R}^{d_2})$ be two arbitrary vectors. We denote $\rho = f(z, s)$. Due to the injectivity of $\tilde{f}$, we have

$$(\tilde{f}_z^{-1}(\rho), \tilde{f}_s^{-1}(\rho)) = \tilde{f}^{-1}(\rho) = \tilde{f}^{-1} \circ f(z, s). \tag{27}$$

Before proceeding to the proof, we provide formal characterizations of the Assumption 3 as follows. We define the compatible pair of ground truth and inferred intrinsic variation as $\mathcal{Z}$. We further denote $\mathcal{Z}_1$ as the first part of $\mathcal{Z}$ (inferred intrinsic variation). We also define the feasible set $\mathcal{S}(\tilde{z}, z)$ of $\tilde{f}_s^{-1}(\rho)$, where $(\tilde{z}, z)$ is any element of $\mathcal{Z}$. The sets $\mathcal{Z}, \mathcal{Z}_1, \mathcal{S}(\tilde{z}, z)$ are defined as follows:

$$\begin{aligned} \mathcal{Z} &= \{(\tilde{z}, z) | \exists (z, s) \in (\mathbb{R}^{d_1}, \mathbb{R}^{d_2}),\ s.t.\ \tilde{z} = \tilde{f}_z^{-1} \circ f(z, s)\}; \\ \mathcal{Z}_1 &= \{\tilde{z} | \exists (z, s) \in (\mathbb{R}^{d_1}, \mathbb{R}^{d_2}),\ s.t.\ \tilde{z} = \tilde{f}_z^{-1} \circ f(z, s)\}; \\ \mathcal{S}(\tilde{z}, z) &= \{\tilde{f}_s^{-1} \circ f(z, s) | \exists s \in \mathbb{R}^{d_2},\ s.t.\ \tilde{z} = \tilde{f}_z^{-1} \circ f(z, s)\}. \end{aligned} \tag{28}$$

(i) in Assumption 3 is equivalent to that, $\mathcal{S}(\tilde{z}, z) = \mathbb{R}^{\tilde{d}_2}$ for any $(\tilde{z}, z)$ in $\mathcal{Z}$.

We next note that, as $f(z, s)$ covers the full set $\mathcal{P}$, and $\tilde{f}^{-1}$ defines a bijective map from $\mathcal{P}$ to $\mathbb{R}^{\tilde{d}_1} \times \mathbb{R}^{\tilde{d}_2}$ , we have that $\mathcal{Z}_1 = \mathbb{R}^{\tilde{d}_1}$.

Now we proceed to prove the lemma. By the injectivity of the function $f$, we can define $F_1 :\ \mathbb{R}^d \to \mathbb{R}^{d_1}$ as the composition of $f_z^{-1}$ and $\tilde{f}$, as follows, $F_1 := f_z^{-1} \circ \tilde{f}$. Then due to the injectivity of $f, \tilde{f}$ we have that

$$z = f_z^{-1}(\rho) = F_1(\tilde{f}^{-1}(\rho)) = F_1(\tilde{f}_z^{-1}(\rho), \tilde{f}_s^{-1}(\rho)). \tag{29}$$

Here $(\tilde{f}_z^{-1}(\rho), z)$ is an arbitrary element in $\mathcal{Z}$. Now we fix it as any element $(\tilde{z}, z) \in \mathcal{Z}$. In this case, we have $\mathcal{S}(\tilde{z}, z) = \mathbb{R}^{\tilde{d}_2}$, which implies that

$$F_1(\tilde{z}, \tilde{s}) = z, \quad \forall \tilde{s} \in \mathbb{R}^{d_2} \text{ and } \forall (\tilde{z}, z) \in \mathcal{Z}. \tag{30}$$

Next, we define a function $F : \mathbb{R}^{\tilde{d}_1} \to \mathbb{R}^{d_1}$ as follows:

$$F(q) := F_1(q, 0), \quad \forall q \in \mathbb{R}^{\tilde{d}_1}. \tag{31}$$

Let $\rho \in \mathcal{P}$. Then there exists a unique $(\tilde{z}, \tilde{s})$, as well as a unique $(z, s)$ such that $\tilde{f}(\tilde{z}, \tilde{s}) = f(z, s) = \rho$. Then, for $q = \tilde{f}_z^{-1}(\rho) = \tilde{z}$, by Eq. (29),

$$z = f_z^{-1}(\rho) = F_1(q, \tilde{f}_s^{-1}(\rho)). \tag{32}$$

However, by Eq. (30), the term on the right hand side is also equal to $F_1(q, 0)$. Hence, we obtain that for any $\rho \in \mathcal{P}$,

$$z = f_z^{-1}(\rho) = F(\tilde{f}_z^{-1}(\rho)) \tag{33}$$

as required.

We next show that $d_1 \leq \tilde{d}_1$ as a classical implication of Sard's theorem [7]. Note that $F$ is continuous and smooth following its definition. By Sard's theorem, the image of the critical set of $F$ is of zero-measure. Here, the image of the critical set refers to set $\mathcal{C} = \{F(x) | x \in \mathbb{R}^{\tilde{d}_1}, \text{rank}(J_F(x)) < d_1\}$. If $d_1 > \tilde{d}_1$, then every $x \in \mathbb{R}^{\tilde{d}_1}$ satisfies that $\text{rank}(J_F(x)) \leq \tilde{d}_1 < d_1$. Further note because $z = f_z^{-1}(\rho)$ is arbitrarily selected in $\mathbb{R}^{d_1}$, the range of $F$ is $\mathbb{R}^{d_1}$. Taken together, $\mathcal{C} = \mathbb{R}^{d_1}$, contradicting with that $\mathcal{C}$ must be of zero-measure in $\mathbb{R}^{d_1}$. Therefore, $d_1 \leq \tilde{d}_1$. Thus by Assumption 4, we have $\tilde{d}_1 = d_1$ thus also $\tilde{d}_2 = d_2$. Together with Eq. (33), (i) and (ii) in Definition 2 are satisfied. □

**Lemma 3. (Functional dependence on inferred and ground truth spatial variation)** *Under assumption 3, $\forall (\tilde{\boldsymbol{\Sigma}}, \tilde{\boldsymbol{A}}, \tilde{f}) \in \Theta$, there exists a function $H : \mathbb{R}^{d_2} \to \mathbb{R}^{\tilde{d}_2}$, such that*

$$\tilde{f}_s^{-1}(\rho) = H(f_s^{-1}(\rho)), \quad \forall \rho \in \mathcal{P}. \tag{34}$$

*This means that for any $\rho$, its corresponding inferred spatial variation $\tilde{f}_s^{-1}(\rho)$ equals a nonlinear mapping of the ground truth spatial variation $f_s^{-1}(\rho)$.*

*Proof.* The proof of the lemma uses an analogous procedure as Lemma 2.

Let $(\tilde{z}, \tilde{s}) \in (\mathbb{R}^{\tilde{d}_1}, \mathbb{R}^{\tilde{d}_2})$ be two arbitrary vectors. We denote $\rho = \tilde{f}(\tilde{z}, \tilde{s})$. Analogous to Lemma 2, we define the sets $\mathcal{S}, \mathcal{S}_1, \mathcal{Z}(s, \tilde{s})$ as follows:

$$\begin{aligned} \mathcal{S} &= \{(s, \tilde{s}) | \exists (\tilde{z}, \tilde{s}) \in (\mathbb{R}^{\tilde{d}_1}, \mathbb{R}^{\tilde{d}_2}),\ s.t.\ s = f_s^{-1} \circ \tilde{f}(\tilde{z}, \tilde{s})\}; \\ \mathcal{S}_1 &= \{s | \exists (\tilde{z}, \tilde{s}) \in (\mathbb{R}^{\tilde{d}_1}, \mathbb{R}^{\tilde{d}_2}),\ s.t.\ s = f_s^{-1} \circ \tilde{f}(\tilde{z}, \tilde{s})\} \\ \mathcal{Z}(s, \tilde{s}) &= \{f_z^{-1} \circ \tilde{f}(\tilde{z}, \tilde{s}) | \exists \tilde{z} \in \mathbb{R}^{\tilde{d}_1},\ s.t.\ s = f_s^{-1} \circ \tilde{f}(\tilde{z}, \tilde{s})\}. \end{aligned} \tag{35}$$

(ii) in Assumption 3 is equivalent to that, $\mathcal{Z}(s, \tilde{s}) = \mathbb{R}^{d_1}$ for any $(s, \tilde{s})$ in $\mathcal{S}$.

As $\tilde{f}(\tilde{z}, \tilde{s})$ covers the full set $\mathcal{P}$, and $f^{-1}$ defines a bijective map from $\mathcal{P}$ to $\mathbb{R}^{d_1} \times \mathbb{R}^{d_2}$ , we have that $\mathcal{S}_1 = \mathbb{R}^{d_2}$. By the injectivity of the function $f$, we can define $H_1 : \mathbb{R}^d \to \mathbb{R}^{\tilde{d}_2}$ as the composition of $\tilde{f}_s^{-1}$ and $f$: $H_1 := \tilde{f}_s^{-1} \circ f$. Then due to the injectivity of $f, \tilde{f}$ we have that

$$\tilde{s} = \tilde{f}_s^{-1}(\rho) = H_1(f^{-1}(\rho)) = H_1(f_z^{-1}(\rho), f_s^{-1}(\rho)). \tag{36}$$

Here $(f_s^{-1}(\rho), \tilde{s})$ is an arbitrary element in $\mathcal{S}$. Now we fix it as any element $(s, \tilde{s}) \in \mathcal{S}$. In this case, we have $\mathcal{Z}(s, \tilde{s}) = \mathbb{R}^{d_1}$, which implies that

$$H_1(z, s) = \tilde{s}, \quad \forall z \in \mathbb{R}^{d_1} \text{ and } \forall (s, \tilde{s}) \in \mathcal{S}. \tag{37}$$

Next, we define a function $H : \mathbb{R}^{d_2} \to \mathbb{R}^{\tilde{d}_2}$ as follows:

$$H(q) := H_1(0, q), \quad \forall q \in \mathbb{R}^{d_2}. \tag{38}$$

Let $\rho \in \mathcal{P}$. Then there exists a unique $(\tilde{z}, \tilde{s})$, as well as a unique $(z, s)$ such that $\tilde{f}(\tilde{z}, \tilde{s}) = f(z, s) = \rho$. Then, for $q = f_s^{-1}(\rho) = s$, by Eq. (36),

$$\tilde{s} = \tilde{f}_s^{-1}(\rho) = H_1(f_z^{-1}(\rho), q) \tag{39}$$

However, by Eq. (37), the term on the right hand side is also equal to $H_1(0, q)$. Hence, we obtain that for any $\rho \in \mathcal{P}$,

$$\tilde{s} = \tilde{f}_s^{-1}(\rho) = H(f_s^{-1}(\rho)) \tag{40}$$

as required. □

**Lemma 4. (Extending equivalence on neighborhood)** *Let $s \in \mathbb{R}^{d_2}$ be an arbitrary vector. For each such $s$, we define $V_s$ as an arbitrary open neighborhood of $s$ in $\mathbb{R}^{d_2}$, that includes $s$ itself. Let $\simeq$ be an equivalence relation on $\mathbb{R}^{d_2}$ satisfying the following property: for any $s, s' \in \mathbb{R}^{d_2}$ and their respective open neighborhoods $V_s, V_{s'} \subseteq \mathbb{R}^{d_2}$,*

$$V_s \cap V_{s'} \neq \emptyset \Rightarrow s \simeq s'. \tag{41}$$

*Then $\forall s, s' \in \mathbb{R}^{d_2}, s \simeq s'$.*

*Proof.* The lemma is employed in the proof of the main theorem to extend the local linear identifiability result to $\mathbb{R}^{d_2}$. The proof of the lemma comprises two steps. First, we show that once two open set neighborhoods $V_s, V_{s'}$ can be "connected" by a finite sequence of $\{V_{s_i}\}$, then $s \simeq s'$ holds. Next, we construct this finite sequence for any $V_s, V_{s'}$, thus concluding the proof.

**Step I.** Formally, we first prove that, $\forall s, s' \in \mathbb{R}^{d_2}$, if there exists a finite sequence of $\{V_{s_i}\}_{i=1}^M$, such that

$$1. \quad \forall I \subsetneq \{1, 2, \ldots, M\} \neq \emptyset, \quad (\cup_{i \in I} V_{s_i}) \cap (\cup_{i \in I^c} V_{s_i}) \neq \emptyset. \tag{42}$$

$$2. \quad V_s \cap (\cup_{i=1}^M V_{s_i}) \neq \emptyset, V_{s'} \cap (\cup_{i=1}^M V_{s_i}) \neq \emptyset. \tag{43}$$

Then $s \simeq s'$.

To prove the claim, we construct an ordering of $\{1, 2, \ldots, M\}$ as follows. By Eq. (43), we can find a set $V_{s_l} \in \{V_{s_i}\}_{i=1}^M$ that intersects with $V_s$. We denote it as $V_{s_{[1]}}$. Next, we iterate over the rest elements from $\{V_{s_i}\}_{i \neq l}$ to find an element that intersects with $V_{s_{[1]}}$, and denote it as $V_{s_{[2]}}$. We find the i+1-th (i>1) element $V_{s_{[i+1]}}$ from the yet unselected elements that satisfies $V_{s_{[i+1]}} \cap (\cup_{j=1}^i V_{s_{[j]}}) \neq \emptyset$. By Eq. (42), such a selection is always possible when $i < M$. Otherwise, we have found a non-empty set $I = \{[1], [2], \ldots, [i]\} \subsetneq \{1, 2, \ldots, M\}$ such that $(\cup_{i \in I} V_{s_i}) \cap (\cup_{i \in I^c} V_{s_i}) = \emptyset$, thus contradicting with Eq. (42).

Now by the construction, we have $s_{[1]} \simeq s_{[2]}$ holds by the non-empty intersection of $V_{s_{[1]}}$ and $V_s$. If for $i < M$, $\forall j_1, j_2 \in \{1, \ldots, i\}, s_{[j_1]} \simeq s_{[j_2]}$, then because $V_{s_{[i+1]}} \cap (\cup_{j=1}^i V_{s_{[j]}}) \neq \emptyset$, it must intersect with at least one element in $\{V_{s_{[j]}}\}_{j=1}^i$. Denote the element as $V_{s_{[l]}}$, then we have that $s_{[i+1]} \simeq s_{[l]}$. Thus by transitivity of $\simeq$, $\forall j_1, j_2 \in \{1, \ldots, i+1\}, s_{[j_1]} \simeq s_{[j_2]}$. It follows by induction that $\forall j_1, j_2 \in \{1, \ldots, M\}, s_{[j_1]} \simeq s_{[j_2]}$. This implies that $\forall i, j \in \{1, \ldots, M\}, s_i \simeq s_j$.

Note that $s \simeq s_{[1]}$ due to the non-empty intersection. Because $V_{s'} \cap (\cup_{i=1}^M V_{s_i}) \neq \emptyset$, $\exists m \in \{1, 2, \ldots, M\}$ such that $V_{s'} \cap V_{s_m} \neq \emptyset$. Then we have $s \simeq s_{[1]} \simeq s_m \simeq s'$ holds. Thus we conclude the first step.

**Step II.** Now we construct such finite sequence $\{V_{s_i}\}$ for any $s, s' \in \mathbb{R}^{d_2}$. We consider the line segment set $\mathcal{S}_{s,s'} := \{ts' + (1-t)s | t \in [0, 1]\} \subseteq \mathbb{R}^{d_2}$. Then $\cup_{\bar{s} \in \mathcal{S}_{s,s'}} V_{\bar{s}}$ form an open cover of $\mathcal{S}_{s,s'}$. Note $\mathcal{S}_{s,s'}$ is closed and bounded on $\mathbb{R}^{d_2}$, $V_{\bar{s}}$ is open for any $\bar{s} \in \mathbb{R}^{d_2}$, and $\cup_{\bar{s} \in \mathcal{S}_{s,s'}} V_{\bar{s}}$ covers $\mathcal{S}_{s,s'}$. By Heine–Borel theorem, we can select a finite sequence of $\{s_i\}_{i=1}^N \subset \mathcal{S}_{s,s'}$ such that $\mathcal{S}_{s,s'} \subseteq \cup_i V_{s_i}$. Thus we have $s, s' \in \cup_{i=1}^N V_{s_i}$, which implies that Eq. (43) is satisfied.

It remains to verify that Eq. (42) for the constructed $\cup_{i=1}^N V_{s_i}$. If not, then there exists a non-empty index set $I \subsetneq \{1, 2, \ldots, N\}$ such that $(\cup_{i \in I} V_{s_i}) \cap (\cup_{i \in I^c} V_{s_i}) = \emptyset$. Thus $(\cup_{i \in I} V_{s_i}) \cap \mathcal{S}_{s,s'}$ and $(\cup_{i \in I} V_{s_i}) \cap \mathcal{S}_{s,s'}$ are non-intersecting non-empty open sets on $\mathcal{S}_{s,s'}$ with union $\mathcal{S}_{s,s'}$. This means that $\mathcal{S}_{s,s'}$ is a disconnected set. However, $\mathcal{S}_{s,s'}$ is obviously path-connected thus connected, leading to a contradiction.

As both conditions (42,43) are satisfied for the finite sequence $\cup_{i=1}^N V_{s_i}$, by step I we have that $s \simeq s'$, thus the proof is completed.

□

**Theorem 1.** **(Model parameter identifiability)** *Under assumptions 1, 2, 3, 4, the model parameter* $(\boldsymbol{\Sigma}, \boldsymbol{A}, f)$ *is* $\sim$*-identifiable on* $\Theta$.

*Proof.* We denote the ground truth parameters of the model by $\theta = (\boldsymbol{\Sigma}, \boldsymbol{A}, f)$ and let $\tilde{\theta} = (\tilde{\boldsymbol{\Sigma}}, \tilde{\boldsymbol{A}}, \tilde{f}) \in \Theta$ be an alternative set of parameters that yield the same gene expression distribution for a single cell.

To establish $\sim$-identifiability, we need to verify that (i), (ii), (iii) in Definition 2 hold for $\theta$ and $\tilde{\theta}$. (i) and (ii) hold by Lemma 2. Thereby the main body of the proof is to verify (iii), i.e., Eq. (13). We present an outline of the proof, which consists of five steps. Steps I-III are preparatory, while steps IV-V cover the main proof. In step I, we show that the function $F$ defined in Lemma 2 is locally injective. In step II, we leverage the local injectivity result from step I to demonstrate that the Jacobian volume vol$J_{\tilde{f}^{-1} \circ f}(z, s)$ can be locally factorized. That is, the volume term can be expressed as a product of separate functions of $z$ and $s$. In step III, we describe a construction of local quadruplets $\{z_i, s, \tilde{z}_i, \tilde{s}\}_{i=0}^{d_2}$. In step IV, we evaluate the probability density of $(\boldsymbol{z}, \boldsymbol{s})$ and $f^{-1} \circ \tilde{f}(\tilde{\boldsymbol{z}}, \tilde{\boldsymbol{s}})$ at these local quadruplets, which shows that Eq. (13) is satisfied locally. In step V, we extend the local result to full space, thus verifying Eq. (13).

**Step I.** Consider the function $h : \mathbb{R}^d \to \mathbb{R}^d$, defined as follows,

$$h = f^{-1} \circ \tilde{f} \equiv [f_z^{-1} \circ \tilde{f}, f_s^{-1} \circ \tilde{f}]. \tag{44}$$

Its Jacobian has the following form:

$$J_{f^{-1} \circ \tilde{f}} = \begin{pmatrix} \partial(f_z^{-1} \circ \tilde{f})/\partial \tilde{z} & \partial(f_z^{-1} \circ \tilde{f})/\partial \tilde{s} \\ \partial(f_s^{-1} \circ \tilde{f})/\partial \tilde{z} & \partial(f_s^{-1} \circ \tilde{f})/\partial \tilde{s} \end{pmatrix}. \tag{45}$$

Since both $f$ and $\tilde{f}$ are diffeomorphisms, $f^{-1} \circ \tilde{f} : \mathbb{R}^d \to \mathbb{R}^d$ is a diffeomorphism as well. Hence, $\det J_{f^{-1} \circ \tilde{f}} \neq 0$ for all $(\tilde{z}, \tilde{s}) \in \mathbb{R}^d$. Moreover, by Eq. (30) of Lemma 2, with $F_1 = f_z^{-1} \circ \tilde{f}$, we have $\partial F_1 / \partial \tilde{s} = \partial (f_z^{-1} \circ \tilde{f}) / \partial \tilde{s} \equiv 0$. In addition, by Lemma 2, $\tilde{d}_1 = d_1$. Thus $\partial (f_z^{-1} \circ \tilde{f}) / \partial \tilde{z} \in \mathbb{R}^{d_1 \times d_1}, \partial (f_s^{-1} \circ \tilde{f}) / \partial \tilde{s} \in \mathbb{R}^{d_2 \times d_2}$. Since the upper right block in Eq. (45) is zero, then

$$\det J_{f^{-1} \circ \tilde{f}} = \det \frac{\partial (f_z^{-1} \circ \tilde{f})}{\partial \tilde{z}} \times \det \frac{\partial f_s^{-1} \circ \tilde{f}}{\partial \tilde{s}} \neq 0. \tag{46}$$

Hence, $\det (\partial f_z^{-1} \circ \tilde{f}(\tilde{z}, \tilde{s}) / \partial \tilde{z}) \neq 0$ for all $\tilde{z}$. By Eqs. (30) and (31) of Lemma 2, $f_z^{-1} \circ \tilde{f}(\tilde{z}, \tilde{s}) \equiv f_z^{-1} \circ \tilde{f}(\tilde{z}, 0) = F(\tilde{z})$. Therefore, the Jacobian $J_F$ of the function $F$ satisfies that $\det J_F \neq 0$ for all $\tilde{z} \in \mathbb{R}^{d_1}$. That is, the function $F$ is locally injective.

**Step II.** Next, we consider the inverse of $h$, namely $h^{-1} = \tilde{f}^{-1} \circ f$. Its Jacobian is given by

$$J_{\tilde{f}^{-1} \circ f} = \left( \begin{array}{c|c} \partial (\tilde{f}_z^{-1} \circ f) / \partial z & \partial (\tilde{f}_z^{-1} \circ f) / \partial s \\ \hline \partial (\tilde{f}_s^{-1} \circ f) / \partial z & \partial (\tilde{f}_s^{-1} \circ f) / \partial s \end{array} \right). \tag{47}$$

By Eq. (37), the lower left block of the matrix is 0. Moreover, by Eqs. (37, 38) of Lemma 3, $\tilde{f}_s^{-1} \circ f(z, s) \equiv \tilde{f}_s^{-1} \circ f(0, s) = H(s)$. Hence, the bottom right block above is equal to $\partial H(s) / \partial s$.

Now we consider the upper-left block $\partial (\tilde{f}_z^{-1} \circ f) / \partial z$. Because the function $F(\tilde{z}) = f_z^{-1} \circ \tilde{f}(\tilde{z}, 0)$ is locally injective, the inverse function theorem can be applied to $F$. That is, for any $\tilde{z}_0 \in \mathbb{R}^{\tilde{d}_1}$, there exists a neighborhood of $\tilde{z}_0$ (open set $V_{\tilde{z}_0} \in \mathbb{R}^{\tilde{d}_2}$) such that $F$ is invertible for all $\tilde{z} \in V_{\tilde{z}_0}$. We denote its local inverse in $V_{\tilde{z}_0}$ as $F_{\tilde{z}_0}^{-1}$. By Eq. (26) of Lemma 2, with $\rho = f(z, s)$ it follows that $F(\tilde{f}_z^{-1} \circ f(z, s)) = z$. Hence, we have that for all $(z, s) \in \mathbb{R}^d$ such that $\tilde{f}_z^{-1} \circ f(z, s) \in V_{\tilde{z}_0}$,

$$F_{\tilde{z}_0}^{-1}(z) = \tilde{f}_z^{-1} \circ f(z, s). \tag{48}$$

Thus the upper-left block of the Jacobian $\partial (\tilde{f}_z^{-1} \circ f(z, s)) / \partial z = dF_{\tilde{z}_0}^{-1}(z) / dz$. Combining the above results for the different blocks in the Jacobian, we have

$$J_{\tilde{f}^{-1} \circ f}(z, s) = \left( \begin{array}{c|c} dF_{\tilde{z}_0}^{-1}(z) / dz & \partial (\tilde{f}_z^{-1} \circ f) / \partial s \\ 0 & dH(s) / ds \end{array} \right). \tag{49}$$

Now we consider the volume of this Jacobian, defined as the absolute value of the determinant $\det J_{\tilde{f}^{-1} \circ f}$. For all $(z, s)$ such that $\tilde{f}_z^{-1} \circ f(z, s) \in V_{\tilde{z}_0}$,

$$\mathrm{vol} J_{\tilde{f}^{-1} \circ f}(z, s) := |\det J_{\tilde{f}^{-1} \circ f}(z, s)| = |\det (\frac{dF_{\tilde{z}_0}^{-1}(z)}{dz})| \times |\det \frac{dH(s)}{ds}|. \tag{50}$$

Denote $h_z^{\tilde{z}_0}(z) = |\det (dF_{\tilde{z}_0}^{-1}(z) / dz)|,\ h_s(s) = |\det (dH(s) / ds)|$, then

$$\mathrm{vol} J_{\tilde{f}^{-1} \circ f}(z, s) = h_z^{\tilde{z}_0}(z) h_s(s). \tag{51}$$

We will use this decomposition in the proof below.

**Step III.** For any $\rho_0 \in \mathcal{P}$, let $(\tilde{z}_0, \tilde{s}_0)$ and $(z_0, s_0)$ be the unique vectors such that $\tilde{f}(\tilde{z}_0, \tilde{s}_0) = f(z_0, s_0) = \rho_0$. As in step II, we denote the local inverse of the function $F$ constructed in Lemma 2 (Eq. (31)) at $\tilde{z}_0$ by $F_{\tilde{z}_0}^{-1}$, and its domain as $V_{\tilde{z}_0}$. In this step, we construct local quadruplets $\{(z_i, s, \tilde{z}_i, \tilde{s})\}_{i=0}^{d_2}$ with desirable properties, as a preparation for step IV. We start by showing that we can select $d_2$ vectors $\{z_i\}_{i=1}^{d_2}$, that together with $z_0$, they satisfy the following properties:

1. There exists an open set $V_{s_0}(\rho_0)$ that includes $s_0$, such that

$$\forall i \in \{0, 1, 2, \dots, d_2\},\ \forall s \in V_{s_0}(\rho_0), \quad \tilde{f}_z^{-1} \circ f(z_i, s) \in V_{\tilde{z}_0}; \tag{52}$$

2. There exists a constant $\epsilon_{\rho_0} > 0$ that does not depend on $i$, such that

$$\forall i \in \{1, 2, \dots, d_2\}, \quad \boldsymbol{A}'(z_i - z_0) = \epsilon_{\rho_0} \boldsymbol{e}_i, \tag{53}$$

where $\boldsymbol{A}' \in \mathbb{R}^{d_2 \times d_1}$ is defined in (9), and $\boldsymbol{e}_i \in \mathbb{R}^{d_2}$ is the $i$-th standard unit vector.

We first show that there exists neighborhoods of $z$ and of $s$, $V_{z_0}(\rho_0)$ and $V_{s_0}(\rho_0)$, that include $z_0$ and $s_0$ respectively and satisfy that: $\forall z \in V_{z_0}(\rho_0), \forall s \in V_{s_0}(\rho_0), \tilde{f}_z^{-1}(f(z, s)) \in V_{\tilde{z}_0}$. Thus $(z_i, s)$ satisfies the first property if $z_i \in V_{z_0}(\rho_0)$ for all $i$.

To construct $V_{z_0}(\rho_0)$ and $V_{s_0}(\rho_0)$, we consider the preimage of $V_{\tilde{z}_0}$ with respect to the function $\tilde{f}_z^{-1} \circ f$. By the property of continuous functions, the preimage of an open set is an open set. Thus by the continuity of $\tilde{f}_z^{-1} \circ f$, the preimage $V_{(z_0, s_0)} := \{(z, s) | \tilde{f}_z^{-1} \circ f(z, s) \in V_{\tilde{z}_0}\}$ is an open set. Further note that we have $(z_0, s_0) \in V_{(z_0, s_0)}$. Hence there exists $\delta_{\rho_0} > 0$, such that the $l_\infty$ neighborhood of $(z_0, s_0)$ with radius $\delta_{\rho_0}$ is inside $V_{(z_0, s_0)}$:

$$\mathbb{B}_\infty((z_0, s_0), \delta_{\rho_0}) = \{(z, s) | \|(z, s) - (z_0, s_0)\|_\infty < \delta_{\rho_0}\} \subseteq V_{(z_0, s_0)}.$$

Now we define

$$V_{z_0}(\rho_0) := \mathbb{B}_\infty(z_0, \delta_{\rho_0}) = \{z | \|z - z_0\|_\infty < \delta_{\rho_0}\}; \quad V_{s_0}(\rho_0) := \mathbb{B}_\infty(s_0, \delta_{\rho_0}) = \{s | \|s - s_0\|_\infty < \delta_{\rho_0}\}.$$

Then we immediately have that,

$$\forall z \in \mathbb{B}_\infty(z_0, \delta_{\rho_0}),\ \forall s \in \mathbb{B}_\infty(s_0, \delta_{\rho_0}),\ (z, s) \in \mathbb{B}_\infty((z_0, s_0), \delta_{\rho_0}) \subseteq V_{(z_0, s_0)} \Rightarrow \tilde{f}_z^{-1} \circ f(z, s) \in V_{\tilde{z}_0}. \tag{54}$$

We next show that we can select $d_2$ vectors $z_i$ within $\mathbb{B}_\infty(z_0, \delta_{\rho_0})$ that also satisfy the second property. Because $\boldsymbol{A}' \in \mathbb{R}^{d_2 \times d_1}$ with $d_2 \leq d_1$ and is of full rank $d_2$ by Assumption 2, it has a right inverse $\boldsymbol{A}'^\dagger \in \mathbb{R}^{d_1 \times d_2}$. Now we select $z_i = z_0 + \epsilon_{\rho_0} \boldsymbol{A}'^\dagger \boldsymbol{e}_i$ for $i \in \{1, 2, \dots, d_2\}$. Then there exists a small enough constant $\epsilon_{\rho_0} > 0$ such that $\max_i \|z_i - z_0\|_\infty = \max_i \|\epsilon_{\rho_0} \boldsymbol{A}'^\dagger \boldsymbol{e}_i\|_\infty < \delta_{\rho_0}$. This implies that $\forall i \in \{1, 2, \dots, d_2\}, z_i \in \mathbb{B}_\infty(z_0, \delta_{\rho_0})$. Moreover, $\boldsymbol{A}'(z_i - z_0) = \epsilon_{\rho_0} \boldsymbol{A}' \boldsymbol{A}'^\dagger \boldsymbol{e}_i = \epsilon_{\rho_0} \boldsymbol{e}_i$. Thus both properties (52,53) hold for selected $\{z_i\}_{i=1}^{d_2}$.

With the described $\{z_i\}_{i=1}^{d_2}$ along with $z_0$, we construct quadruplets $\{(z_i, s, \tilde{f}_z^{-1} \circ f(z_i, s), \tilde{f}_s^{-1} \circ f(z_i, s))\}_{i=0}^{d_2}$. Here $s$ is any vector in $\mathbb{B}_\infty(s_0, \delta_{\rho_0})$ and does not depend on $i$. By property (52), $\tilde{f}_z^{-1} \circ f(z_i, s) \in V_{\tilde{z}_0}$, thus by Eq. (48) it is equal to $F_{\tilde{z}_0}^{-1}(z_i)$, which we denote as $\tilde{z}_i$. By Lemma 3, denote $\rho_i = f(z_i, s)$, then we have $\tilde{f}_s^{-1} \circ f(z_i, s) = \tilde{f}_s^{-1}(\rho_i) = H(f_s^{-1}(\rho_i)) = H(s)$ independent of $i$, which we denote as $\tilde{s}$. Together, the quadruplets can be rewritten as $\{(z_i, s, \tilde{z}_i, \tilde{s})\}_{i=0}^{d_2}$. Note that by definition, for any $i \in \{0, 1, 2, \dots, d_2\}$,

$$f(z_i, s) = \tilde{f} \circ \tilde{f}^{-1} \circ f(z_i, s) = \tilde{f}(\tilde{f}_z^{-1} \circ f(z_i, s), \tilde{f}_s^{-1} \circ f(z_i, s)) = \tilde{f}(\tilde{z}_i, \tilde{s}). \tag{55}$$

**Step IV.** By Eq. (14) of Lemma 1, $f(\boldsymbol{z}, \boldsymbol{s}) \stackrel{d}{=} \tilde{f}(\tilde{\boldsymbol{z}}, \tilde{\boldsymbol{s}})$. By the injectivity of $f^{-1}$, this is equivalent to $(\boldsymbol{z}, \boldsymbol{s}) \stackrel{d}{=} h(\tilde{\boldsymbol{z}}, \tilde{\boldsymbol{s}}) := f^{-1} \circ \tilde{f}(\tilde{\boldsymbol{z}}, \tilde{\boldsymbol{s}})$. This means that for any $\rho \in \mathcal{P}$,

$$p((\boldsymbol{z}, \boldsymbol{s}) = f^{-1}(\rho)) = p(h(\tilde{\boldsymbol{z}}, \tilde{\boldsymbol{s}}) = f^{-1}(\rho)). \tag{56}$$

For simplicity, here we omit $\theta, \tilde{\theta}$ in $p_\theta, p_{\tilde{\theta}}$. Analogous to the treatment in [2], we first apply a change of variables formula to the RHS of Eq. (56), by the injectivity of $h$. Taking into account that $h^{-1} \circ f^{-1}(\rho) = \tilde{f}^{-1}(\rho)$, this yields the following formula,

$$p(h(\tilde{\boldsymbol{z}}, \tilde{\boldsymbol{s}}) = f^{-1}(\rho)) = \mathrm{vol} J_{h^{-1}}(f^{-1}(\rho)) p((\tilde{\boldsymbol{z}}, \tilde{\boldsymbol{s}}) = \tilde{f}^{-1}(\rho)).$$

Taking logarithms in Eq. (56) thus gives

$$\log p((\boldsymbol{z}, \boldsymbol{s}) = f^{-1}(\rho)) = \log \mathrm{vol} J_{\tilde{f}^{-1} \circ f}(f^{-1}(\rho)) + \log p((\tilde{\boldsymbol{z}}, \tilde{\boldsymbol{s}}) = \tilde{f}^{-1}(\rho)).$$

Next, we decompose the joint distribution $p(\boldsymbol{z}, \boldsymbol{s})$ into product of marginal and conditional distributions, Namely, we write the left hand side as

$$\log p((\boldsymbol{z}, \boldsymbol{s}) = f^{-1}(\rho)) = \log p(\boldsymbol{z} = f_z^{-1}(\rho)) + \log p(\boldsymbol{s} = f_s^{-1}(\rho) | \boldsymbol{z} = f_z^{-1}(\rho))$$

with the right hand side admitting a similar decomposition. Hence Eq. (56) can be written as follows:

$$\begin{aligned}&\log p(\boldsymbol{z}=f_z^{-1}(\rho))+\log p(\boldsymbol{s}=f_s^{-1}(\rho)|\boldsymbol{z}=f_z^{-1}(\rho))\\=&\log \mathrm{vol}J_{\tilde{f}^{-1}\circ f}(f^{-1}(\rho))+\log p(\tilde{\boldsymbol{z}}=\tilde{f}_z^{-1}(\rho))+\log p(\tilde{\boldsymbol{s}}=\tilde{f}_s^{-1}(\rho)|\tilde{\boldsymbol{z}}=\tilde{f}_z^{-1}(\rho)).\end{aligned} \tag{57}$$

The probability densities $p(\boldsymbol{z}), p(\boldsymbol{s}|\boldsymbol{z})$ and $p(\tilde{\boldsymbol{z}}), p(\tilde{\boldsymbol{s}}|\tilde{\boldsymbol{z}})$ are multivariate Gaussians characterized by Eqs. (7,10) with parameters $(\boldsymbol{\Sigma}, \boldsymbol{A})$ and $(\tilde{\boldsymbol{\Sigma}}, \tilde{\boldsymbol{A}})$ respectively. Together we have: $\forall \rho \in \mathcal{P}$,

$$\begin{aligned}-\frac{d}{2}\log 2\pi-\frac{1}{2}\|f_z^{-1}(\rho)\|^2-\frac{1}{2}|\boldsymbol{\Sigma}'|-\frac{1}{2}(f_s^{-1}(\rho)-\boldsymbol{A}'f_z^{-1}(\rho))^T\boldsymbol{\Sigma}'^{-1}(f_s^{-1}(\rho)-\boldsymbol{A}'f_z^{-1}(\rho))\\=\log \mathrm{vol}J_{\tilde{f}^{-1}\circ f}(f^{-1}(\rho))-\frac{d}{2}\log 2\pi-\frac{1}{2}\|\tilde{f}_z^{-1}(\rho)\|^2-\frac{1}{2}|\tilde{\boldsymbol{\Sigma}}'|-\\\frac{1}{2}(\tilde{f}_s^{-1}(\rho)-\tilde{\boldsymbol{A}}'\tilde{f}_z^{-1}(\rho))^T\tilde{\boldsymbol{\Sigma}}'^{-1}(\tilde{f}_s^{-1}(\rho)-\tilde{\boldsymbol{A}}'\tilde{f}_z^{-1}(\rho))\end{aligned} \tag{58}$$

For any quadruplets $(z, s, \tilde{z}, \tilde{s}) \in (\mathbb{R}^{d_1}, \mathbb{R}^{d_2}, \mathbb{R}^{d_1}, \mathbb{R}^{d_2})$ that satisfy $f(z, s) = \tilde{f}(\tilde{z}, \tilde{s}) = \rho$, Eq. (58) simplifies to

$$\begin{aligned}-\frac{1}{2}\|z\|^2-\frac{1}{2}|\boldsymbol{\Sigma}'|-\frac{1}{2}(s-\boldsymbol{A}'z)^T\boldsymbol{\Sigma}'^{-1}(s-\boldsymbol{A}'z)\\=\log \mathrm{vol}J_{\tilde{f}^{-1}\circ f}(z,s)-\frac{1}{2}\|\tilde{z}\|^2-\frac{1}{2}|\tilde{\boldsymbol{\Sigma}}'|-\frac{1}{2}(\tilde{s}-\tilde{\boldsymbol{A}}'\tilde{z})^T\tilde{\boldsymbol{\Sigma}}'^{-1}(\tilde{s}-\tilde{\boldsymbol{A}}'\tilde{z})\end{aligned} \tag{59}$$

For any $\rho_0 \in \mathcal{P}$, we denote the unique $(\tilde{z}_0, \tilde{s}_0)$ and the unique $(z_0, s_0)$ such that $\tilde{f}(\tilde{z}_0, \tilde{s}_0) = f(z_0, s_0) = \rho_0$. Based on $\rho_0$ and its corresponding $(z_0, s_0, \tilde{z}_0, \tilde{s}_0)$, we consider the quadruplets constructed in step III: $\{(z_i, s, \tilde{z}_i, \tilde{s})\}_{i=0}^{d_2}$. By Eq. (55), we have $f(z_i, s) = \tilde{f}(\tilde{z}_i, \tilde{s})$ for any $i \in \{0, 1, 2, \ldots, d_2\}$. Thus we can plug them in Eq. (59), yielding $d_2 + 1$ equations that must be satisfied for any parameter $\tilde{\theta} \in \Theta$. By Eq. (52), all $\tilde{z}_i \in V_{\tilde{z}_0}$. Thus by Eq. (51) in step II, $\forall i \in \{0, \ldots, d_2\}, \log \mathrm{vol}J_{\tilde{f}^{-1}\circ f}(z_i, s) = \log h_z^{\tilde{z}_0}(z_i) + \log h_s(s)$. Hence the form of the $d_2 + 1$ equations are as follows: $\forall i \in \{0, \ldots, d_2\}, \forall s \in \mathbb{B}_\infty(s_0, \delta_{\rho_0})$,

$$\begin{aligned}-\frac{1}{2}\|z_i\|^2-\frac{1}{2}|\boldsymbol{\Sigma}'|-\frac{1}{2}(s-\boldsymbol{A}'z_i)^T\boldsymbol{\Sigma}'^{-1}(s-\boldsymbol{A}'z_i)\\=\log h_z^{\tilde{z}_0}(z_i)+\log h_s(s)-\frac{1}{2}\|\tilde{z}_i\|^2-\frac{1}{2}|\tilde{\boldsymbol{\Sigma}}'|-\frac{1}{2}(\tilde{s}-\tilde{\boldsymbol{A}}'\tilde{z}_i)^T\tilde{\boldsymbol{\Sigma}}'^{-1}(\tilde{s}-\tilde{\boldsymbol{A}}'\tilde{z}_i)\end{aligned} \tag{60}$$

To obtain linear equations for $s$ of form Eq. (13), we subtract the first equation for $(z_0, s, \tilde{z}_0, \tilde{s})$ from the remaining $d_2$ equations, yielding $d_2$ new equations. After this substraction, constant terms, quadratic terms of $s, \tilde{s}$, and the log Jacobian volume term $\log h_s(s)$ all cancel out. Furthermore, recall that by Eq. (53), $\boldsymbol{A}'z_i - \boldsymbol{A}'z_0 = \epsilon_{\rho_0}\boldsymbol{e}_i$ for all $i \in \{1, 2, \ldots, d_2\}$. Hence, the $d_2$ equations can be simplified as: $\forall i \in \{1, \ldots, d_2\}$,

$$\begin{aligned}2\log h_z^{\tilde{z}_0}(z_i)-2\log h_z^{\tilde{z}_0}(z_0)+\|z_i\|^2-\|z_0\|^2-2\epsilon_{\rho_0}\boldsymbol{e}_i^T\boldsymbol{\Sigma}'^{-1}s+(\boldsymbol{A}'z_i)^T\boldsymbol{\Sigma}'^{-1}(\boldsymbol{A}'z_i)-(\boldsymbol{A}'z_0)^T\boldsymbol{\Sigma}'^{-1}(\boldsymbol{A}'\tilde{z}_0)\\=\|\tilde{z}_i\|^2-\|\tilde{z}_0\|^2-2(\tilde{\boldsymbol{A}}'(\tilde{z}_i-\tilde{z}_0))^T\tilde{\boldsymbol{\Sigma}}'^{-1}\tilde{s}+(\tilde{\boldsymbol{A}}'\tilde{z}_i)^T\tilde{\boldsymbol{\Sigma}}'^{-1}(\tilde{\boldsymbol{A}}'\tilde{z}_i)-(\tilde{\boldsymbol{A}}'\tilde{z}_0)^T\tilde{\boldsymbol{\Sigma}}'^{-1}(\tilde{\boldsymbol{A}}'\tilde{z}_0)\end{aligned} \tag{61}$$

Now we rewrite these $d_2$ equations in matrix form. We define matrices

$$\boldsymbol{B}_{\rho_0} \in \mathbb{R}^{d_2\times d_2}, \text{ with i-th row } \boldsymbol{B}_{\rho_0 i} = -2\epsilon_{\rho_0}\boldsymbol{e}_i^T\boldsymbol{\Sigma}'^{-1};$$

$$\tilde{\boldsymbol{B}}_{\rho_0} \in \mathbb{R}^{d_2\times d_2}, \text{ with i-th row } \tilde{\boldsymbol{B}}_{\rho_0 i} = -2(\tilde{\boldsymbol{A}}'(\tilde{z}_i-\tilde{z}_0))^T\boldsymbol{\Sigma}'^{-1}.$$

Finally we denote the constant term $\boldsymbol{c}_{\rho_0} \in \mathbb{R}^{d_2}$ with $\boldsymbol{c}_{\rho_0 i} = -2\log h_z^{\tilde{z}_0}(z_i) + 2\log h_z^{\tilde{z}_0}(z_0) - \|z_i\|^2 + \|z_0\|^2 - (\boldsymbol{A}'z_i)^T\boldsymbol{\Sigma}'^{-1}(\boldsymbol{A}'z_i) + (\boldsymbol{A}'z_0)^T\boldsymbol{\Sigma}'^{-1}(\boldsymbol{A}'\tilde{z}_0) + \|\tilde{z}_i\|^2 - \|\tilde{z}_0\|^2 + (\tilde{\boldsymbol{A}}'\tilde{z}_i)^T\tilde{\boldsymbol{\Sigma}}'^{-1}(\tilde{\boldsymbol{A}}'\tilde{z}_i) - (\tilde{\boldsymbol{A}}'\tilde{z}_0)^T\tilde{\boldsymbol{\Sigma}}'^{-1}(\tilde{\boldsymbol{A}}'\tilde{z}_0)$. Then Eq. (61) can be simplified as

$$\boldsymbol{B}_{\rho_0}s = \tilde{\boldsymbol{B}}_{\rho_0}\tilde{s} + \boldsymbol{c}_{\rho_0}.$$

By the construction of the quadruplets, this equation holds for any $s \in \mathbb{B}_\infty(s_0, \delta_{\rho_0})$ and the corresponding $\tilde{s} = H(s)$. Furthermore, the definition of $\boldsymbol{B}_{\rho_0}$ implies that $\boldsymbol{B}_{\rho_0} = -2\epsilon_{\rho_0}\boldsymbol{\Sigma}'^{-1}$ , therefore it is invertible. We denote $L_{\rho_0} = \boldsymbol{B}_{\rho_0}^{-1}\tilde{\boldsymbol{B}}_{\rho_0}$ and $c_{\rho_0} = \boldsymbol{B}_{\rho_0}^{-1}\boldsymbol{c}_{\rho_0}$. Together, we have:

$$\forall s \in \mathbb{B}_\infty(s_0, \delta_{\rho_0}), \quad s = L_{\rho_0}\tilde{s} + c_{\rho_0}. \tag{62}$$

We next show the matrix $L_{\rho_0} \in \mathbb{R}^{d_2 \times d_2}$ is invertible. Because $\mathbb{B}_\infty(s_0, \delta_{\rho_0})$ is a non-empty open set, there exists a neighborhood of $s_0$ in $\mathbb{B}_\infty(s_0, \delta_{\rho_0})$ that includes all vectors $\{s_0 + \epsilon_s \boldsymbol{e}_1, \ldots, s_0 + \epsilon_s \boldsymbol{e}_{d_2}\}$, where $\epsilon_s > 0$ is a small enough constant. Hence, $L_{\rho_0}$ must be invertible, otherwise the range of $L_{\rho_0}\tilde{s} + c_{\rho_0}$ would be in a subspace of dimension lower than $d_2$, leading to a contradiction.

**Step V.** In this step, we demonstrate that the local linear relation expressed in Eq. (62) extends globally. Specifically, we prove the existence of universal parameters $L$ and $c$ such that $L_{\rho_0} = L, c_{\rho_0} = c$ for all $\rho_0 \in \mathcal{P}$. Combining this with Eq. (62) leads directly to the formulation presented in Eq. (13).

We first show that, for any pair of points $\rho_0, \rho_0' \in \mathcal{P}$ that result in non-empty intersection: $\mathbb{B}_\infty(s_0, \delta_{\rho_0}) \cap \mathbb{B}_\infty(s_0', \delta_{\rho_0'}) \neq \emptyset$, we have $L_{\rho_0} = L_{\rho_0'}$ and $c_{\rho_0} = c_{\rho_0'}$. Denote the intersection as $\bar{V}_s$. Then $\bar{V}_s$ is the intersection of two open sets thus is also a open set. This means that for a point $s \in \bar{V}_s$, there exists a neighborhood of $s$ that includes all vectors $\{s_i\}_{i=1}^{d_2} = \{s + \epsilon_s \boldsymbol{e}_1, \ldots, s + \epsilon_s \boldsymbol{e}_{d_2}\}$, where $\epsilon_s > 0$ is a small enough constant. Thus following Eq. (62) and the invertibility of $L_{\rho_0}, L_{\rho_0'}$, we have the following two equations hold:

$$\tilde{s} = L_{\rho_0}^{-1} s - L_{\rho_0}^{-1} c_{\rho_0} = L_{\rho_0'}^{-1} s - L_{\rho_0'}^{-1} c_{\rho_0'}; \tag{63}$$

$$\forall i \in \{1, 2, \ldots, d_2\}, \ \tilde{s}_i = L_{\rho_0}^{-1}(s + \epsilon_s \boldsymbol{e}_i) - L_{\rho_0}^{-1} c_{\rho_0} = L_{\rho_0'}^{-1}(s + \epsilon_s \boldsymbol{e}_i) - L_{\rho_0'}^{-1} c_{\rho_0'}. \tag{64}$$

Taking differences of the two equations yield: $\forall i \in \{1, 2, \ldots, d_2\}$,

$$\epsilon_s L_{\rho_0}^{-1} \boldsymbol{e}_i = \epsilon_s L_{\rho_0'}^{-1} \boldsymbol{e}_i \Rightarrow L_{\rho_0} = L_{\rho_0'}. \tag{65}$$

Subtituting $L_{\rho_0'}$ with $L_{\rho_0}$ in Eq. (63) leads to

$$L_{\rho_0}^{-1} c_{\rho_0} = L_{\rho_0'}^{-1} c_{\rho_0'} \Rightarrow c_{\rho_0} = c_{\rho_0'}. \tag{66}$$

Together, $\forall \rho_0, \rho_0' \in \mathcal{P}$,

$$\mathbb{B}_\infty(s_0, \delta_{\rho_0}) \cap \mathbb{B}_\infty(s_0', \delta_{\rho_0'}) \neq \emptyset \Rightarrow L_{\rho_0} = L_{\rho_0'}, c_{\rho_0} = c_{\rho_0'}. \tag{67}$$

Now we demonstrate that Eq. (67) can be extended to the entire space $\mathbb{R}^{d_2}$, thereby proving (iii) in Definition 2. We first define an equivalence relationship with respect to $s$ based on Eq. (67), then prove that this equivalence relationship holds for any $s_0, s_0' \in \mathbb{R}^{d_2}$ by applying Lemma 4. Finally, we show that (iii) in Definition 2 holds.

For any $s_0 \in \mathbb{R}^{d_2}$, we consider $\bar{\rho}_0 = f(0, s_0)$. Plugging $\bar{\rho}_0$ in Eq. (62) leads to

$$\forall s \in \mathbb{B}_\infty(s_0, \delta_{\bar{\rho}_0}), \quad s = L_{\bar{\rho}_0}\tilde{s} + c_{\bar{\rho}_0}. \tag{68}$$

Because $\bar{\rho}_0$ is uniquely determined by $s_0$, we can denote $L_{s_0} := L_{\bar{\rho}_0}, c_{s_0} := c_{\bar{\rho}_0}, \delta_{s_0} := \delta_{\bar{\rho}_0}, V_{s_0} := \mathbb{B}_\infty(s_0, \delta_{s_0})$. Then the above equation can be rewritten as

$$\forall s \in \mathbb{B}_\infty(s_0, \delta_{s_0}) = V_{s_0}, \quad s = L_{s_0}\tilde{s} + c_{s_0}. \tag{69}$$

Similarly we can define $\bar{\rho}_0', L_{s_0'}, c_{s_0'}, \delta_{s_0'}, V_{s_0'}$ for $s_0' \in \mathbb{R}^{d_2}$. Then we define the equivalence relationship $\simeq$ as follows:

$$s_0 \simeq s_0' \Leftrightarrow L_{s_0} = L_{s_0'} \text{ and } c_{s_0} = c_{s_0'}. \tag{70}$$

Plugging $\bar{\rho}_0, \bar{\rho}_0'$ in Eq. (67) leads to that, $\forall s_0, s_0' \in \mathbb{R}^{d_2}$,

$$V_{s_0} \cap V_{s_0'} \neq \emptyset \Rightarrow s_0 \simeq s_0'. \tag{71}$$

We next apply Lemma 4 based on the equivalence relation (70) and the condition (71). This yields

$$\forall s_0, s_0' \in \mathbb{R}^{d_2}, \quad s_0 \simeq s_0'; \quad L_{s_0} = L_{s_0'},\ c_{s_0} = c_{s_0'}. \tag{72}$$

Now we consider arbitrary $\rho_0 = f(z_0, s_0)$ and $\rho_0' = f(z_0', s_0')$ where $z_0, z_0'$ are arbitrarily selected in $\mathbb{R}^{d_1}$. Because $\mathbb{B}_\infty(s_0, \delta_{\rho_0})$ and $\mathbb{B}_\infty(s_0, \delta_{\bar{\rho}_0})$ must intersect (at $s_0$), by Eq. (67) we have that $L_{\rho_0} = L_{\bar{\rho}_0} = L_{s_0}$ and $c_{\rho_0} = c_{\bar{\rho}_0} = c_{s_0}$. Similarly for $\rho'$, we have $L_{\rho_0'} = L_{\bar{\rho}_0'} = L_{s_0'}$ and $c_{\rho_0'} = c_{\bar{\rho}_0'} = c_{s_0'}$. Thus by Eq. (72) we have

$$L_{\rho_0} = L_{s_0} = L_{s_0'} = L_{\rho_0'}; \quad c_{\rho_0} = c_{s_0} = c_{s_0'} = c_{\rho_0'}. \tag{73}$$

That is, $L_{\rho_0} = L_{\rho_0'}$ and $c_{\rho_0} = c_{\rho_0'}$ hold for all $\rho_0, \rho_0' \in \mathcal{P}$. Hence, $L_{\rho_0}, c_{\rho_0}$ in Eq. (62) can be substituted with universal parameters, which we denote as $L, c$ respectively. Note that $s_0 = f_s^{-1}(\rho_0) \in \mathbb{B}_\infty(s_0, \delta_{\rho_0})$ for any $\rho_0$. Hence we can plug $s = f_s^{-1}(\rho_0)$ and $\tilde{s} = H(s_0) = \tilde{f}_s^{-1}(\rho_0)$ (Lemma 3) in Eq. (62) for any $\rho_0$, which leads to

$$f_s^{-1}(\rho_0) = L\tilde{f}_s^{-1}(\rho_0) + c. \tag{74}$$

Therefore, $\forall \rho \in \mathcal{P},\ f_s^{-1}(\rho) = L\tilde{f}_s^{-1}(\rho) + c$ holds; that is, Eq. (13) is satisfied. The other two conditions (i), (ii) have been verified in Lemma 2. Together, we have $\tilde{\theta} \sim \theta$. Therefore, the ground truth model parameter $(\boldsymbol{\Sigma}, \boldsymbol{A}, f)$ is $\sim$-identifiable on the parameter set $\Theta$. □

*Remark* 7. Our theoretical result has the following implications:

1. There exists a (smooth and locally injective) function $F$ from inferred to ground truth intrinsic variations. This means that the inferred intrinsic variations preserve the (non-parametric) neighborhood in the ground truth intrinsic variation space, i.e., cells adjacent in the inferred intrinsic variation space would also be adjacent in the ground truth intrinsic variation space. Therefore, the inferred intrinsic variations can be used for various downstream analysis, such as clustering and visualization.

2. There exists an invertible linear transformation between ground truth and inferred spatial variations. This is owing to the "implicit supervision" of intrinsic variation to the spatial variation, which is uniquely addressed in our work. This means that in addition to preserving the neighborhood, the spatial variation preserves the (Euclidean) geometry of the ground truth spatial variation. This serves as a basis for our spatial effect estimation method that uses linear archetypal analysis.

#### 1.4 Connections between the theoretical support and SIMVI design

SIMVI uses the variational autoencoder (VAE) architecture to estimate intrinsic and spatial-induced latent variables from data. Here we describe how key designs of the SIMVI model align with the established theoretical support enforcing identifiability. In the following text, $(\boldsymbol{z}, \boldsymbol{s})$ refers to ground truth intrinsic / spatial variations, and $(\tilde{\boldsymbol{z}}, \tilde{\boldsymbol{s}})$ refers to inferred intrinsic / spatial variations by the SIMVI model.

- SIMVI models the intrinsic variation $\tilde{\boldsymbol{z}}$ as a variational posterior of a cell's gene expression, and the spatial variation $\tilde{\boldsymbol{s}}$ as a variational posterior of cell neighborhood. The variational posteriors include encoders $\phi_1 : \mathbb{R}^p \to \mathbb{R}^{\tilde{d}_1}, \phi_2 : \mathbb{R}^p \to \mathbb{R}^{\tilde{d}_1}, \psi_1 : \mathbb{R}^{\tilde{d}_1 \times k} \to \mathbb{R}^{\tilde{d}_2}, \psi_2 : \mathbb{R}^{\tilde{d}_1 \times k} \to \mathbb{R}^{\tilde{d}_2}$. The encoder $\phi_1$ is used for both $\tilde{\boldsymbol{z}}$ and $\tilde{\boldsymbol{s}}$:

$$\tilde{\boldsymbol{z}}|\boldsymbol{x} \sim \mathcal{N}(\phi_1(\boldsymbol{x}), \phi_2(\boldsymbol{x})); \tag{75}$$

$$\tilde{\boldsymbol{s}}|\boldsymbol{x}^N \sim \mathcal{N}(\psi_1(\phi_1(\boldsymbol{x}^N)), \psi_2(\phi_1(\boldsymbol{x}^N))) \approx \mathcal{N}(\psi_1(\tilde{\boldsymbol{z}}^N), \psi_2(\tilde{\boldsymbol{z}}^N)). \tag{76}$$

  Notably, the parameters $(\boldsymbol{A}, \boldsymbol{\Sigma}_{\boldsymbol{s}})$ in the generative process Eq. (2) correspond to the terms $\psi_1, \psi_2$ in Eq. (76), by the design of shared encoder $\phi_1$. In practice, $\psi_1, \psi_2$ are modeled by graph attention networks, which are essentially weighted linear average of its inputs with adaptative weights, whereas $\phi_1, \phi_2$ are general non-linear functions implemented by multilayer perceptrons (MLP). Together, SIMVI variational posteriors effectively model the latent variables in the considered generative process.

- The variational posterior design of $\tilde{s}$ ensures that the range of $\tilde{s}|z$ spans $\mathbb{R}^{d_2}$. This is because $\tilde{s}$ can be viewed as a function of $x^N$, and fixing $z$ does not alter the range of $x^N$ in the generative process. Furthermore, SIMVI imposes additional regularizations on the marginals $(\tilde{z}, \tilde{s})$ to enforce their independence. Consequently, conditioning on $\tilde{z}$ does not affect the range of $\tilde{s}$. Overall, the design choices of SIMVI collectively enforce condition (i) in Assumption 3.
- By definition, conditioning on $s$ does not alter the range of $z$. Analogously, the variational posterior design of $\tilde{s}$ enforces that $\text{supp}(z|\tilde{s})$ spans $\mathbb{R}^{d_1}$ for any fixed $\tilde{s}$. Consequently, the SIMVI model also enforces condition (ii) in Assumption 3.
- Our theoretical analysis considers the latent variables $(\tilde{z}, \tilde{s})$ with the smallest dimensionality of $\tilde{z}$ (Assumption 4). While this cannot be perfectly implemented by VAEs, SIMVI employs an asymmetric independence regularization term to minimize the encoded information in $\tilde{z}$, which can be seen as minimizing the "effective dimensionality" of $\tilde{z}$.
- The SIMVI spatial variational posterior design enforces meaningful outputs even under model misspecification, such as when components of $s$ completely overlap with $z$ in certain cells. This scenario often manifests as identical cell types within a specific spatial niche. In such case, SIMVI would appropriately attribute the variation to both intrinsic and spatial-induced factors. This is because, after training the model, this overlapped information is preserved either in $\tilde{z}$ itself or in $\tilde{s}$, and in the latter case, also in $\tilde{z}^N$. Here, the intrinsic property of the spatial neighborhood is equivalent to $\tilde{z}$, ensuring the overlapping information to be encoded in the intrinsic variation $\tilde{z}$, thus also in $\tilde{z}^N$ and $\tilde{s}$. Nevertheless, this represents a scenario where the spatial effect cannot be effectively estimated for these cells / spatial niches. This is further elaborated in the subsequent section on spatial effect estimation.

### 2 Spatial effect estimation

In this section, we no longer distinguish inferred and ground truth variations, and denote the SIMVI intrinsic / spatial variations as $z, s$ respectively. To estimate the spatial effect for cell $i$ within its spatial context, we consider the spatial variation $s$ as a continuous treatment on each cell. This naturally leads to the formulation of the following conditional average treatment effect objective for cell $i$:

$$\text{SE}_i = \mathbb{E}[Y(s = s^i)|z = z^i] - \mathbb{E}[Y(s = s^0)|z = z^i] \tag{77}$$

Here $Y$ represents the normalized expression of the gene of interest, and $s^i$, $z^i$ denote the spatial and intrinsic representation of cell $i$ respectively. $s^0$ indicates the control spatial state. Before proceeding, we introduce the archetype transformation employed to substitute the original spatial variation.

#### 2.1 Transforming spatial variation into archetypes

After training the SIMVI model, we obtain the spatial variation for each cell $s^i$. Ideally, we aim to investigate how individual "mechanisms" within this spatial variation affect gene expression. However, due to the linear identifiability result, the values of each component $s_j$ represent a linear mixture of these mechanisms rather than independent factors. Consequently, we require a linear-transformation-agnostic approach to estimate the effect of spatial microenvironments.

Our key insight is that while spatial variation comprises multi-dimensional mechanisms, spatial microenvironments are typically mutually exclusive. For instance, in data with a layered structure, each cell's spatial environment can be viewed as a weighted average of microenvironments from adjacent layers. Thus, we can consider pure individual spatial mechanisms as "archetypes" within the spatial variation space. Such archetypes can be derived via linear archetypal analysis:

**Definition 4.** (Archetypal analysis [8]) Suppose $S \in \mathbb{R}^{n \times d_2}$ to be the SIMVI spatial variation (or any data of interest), and $C \in \mathbb{R}^{a \times d_2}$ is a (factor) matrix, with each row within $\text{Conv}(S)$ and linear independent. $Q \in \mathbb{R}^{n \times a}$ is a (loading) matrix, with each row $q^i$ satisfying $q^i \geq 0, \|q^i\|_1 = 1$. Then archetypal analysis finds the optimal $Q, C$ satisfying the described properties and minimize

$$\|S - QC\|_F^2.$$

Applying archetypal analysis on SIMVI spatial variation, we now have a new archetype weight matrix $\boldsymbol{Q}$. Replacing the spatial variation with archetype weights in spatial effect analysis has the following advantages:

1. The archetype weight matrix is able to account for the mutually exclusive nature of the spatial microenvironment. It also leads to a natural definition of the control state $\boldsymbol{q}^0 = \mathbf{0}$.
2. The archetype weight matrix is agnostic to the linear transformation. In this case, we can estimate the treatment effect of each component $\boldsymbol{q}_j$ using the continuous treatment effect estimation framework.

Now, we are able to estimate the treatment effect of each entry (archetype) $j$ in the matrix $\boldsymbol{q}_j$, conditioning on the covariate of individual cells. We next introduce the double machine learning framework that we have adopted for estimating the spatial effect. To make the connection between $\boldsymbol{S}$ and $\boldsymbol{Q}$ clearer, in the following text, we use $\boldsymbol{S}'$ and $\boldsymbol{s}'$ to denote $\boldsymbol{Q}$, $\boldsymbol{q}$ respectively.

#### 2.2 Estimating spatial effect via double machine learning

The first step of double machine learning (DML) framework [9] involves fitting two linear regression models $\hat{Y}(\boldsymbol{z})$, $\hat{\boldsymbol{s}}'(\boldsymbol{z})$ and computing residuals $\bar{Y}$, $\bar{\boldsymbol{s}}'$:

$$\begin{aligned}\bar{Y} &= Y - \hat{Y}(\boldsymbol{z});\\ \bar{\boldsymbol{s}}' &= \boldsymbol{s}' - \hat{\boldsymbol{s}}'(\boldsymbol{z}).\end{aligned} \tag{78}$$

After computing the residuals, the DML framework further considers the following linear model:

$$\bar{Y} \sim \theta(\boldsymbol{z})\bar{\boldsymbol{s}}'. \tag{79}$$

Note that $\bar{\boldsymbol{s}}'$ is multidimensional, therefore when $\theta_i(\boldsymbol{z}) = \boldsymbol{z}\beta_i + c_i$, the model can be further written as:

$$\bar{Y} \sim \sum_j \bar{\boldsymbol{s}}'_j \boldsymbol{z}\beta_j + c_j \bar{\boldsymbol{s}}'_j \tag{80}$$

Solving this regression model is equivalent to solving an ordinary linear regression problem with the covariate matrix as $[\bar{\boldsymbol{s}}'_1\boldsymbol{z}, \ldots, \bar{\boldsymbol{s}}'_a\boldsymbol{z}, \bar{\boldsymbol{s}}'_1, \ldots, \bar{\boldsymbol{s}}'_a]$.

After fitting the model, the spatial effect is obtained as

$$\sum_i \bar{\boldsymbol{s}}'_j \boldsymbol{z}\beta_j + c_j \bar{\boldsymbol{s}}'_j. \tag{81}$$

The DML approach can also be used to derive the variance decomposition for the intrinsic and spatial variation. The $R^2$ for the intrinsic variation is defined as $R^2(Y, \tilde{Y}(\boldsymbol{z}))$, whereas the $R^2$ for the spatial variation is defined as $R^2(\bar{Y}, \sum_j \bar{\boldsymbol{s}}'_j \boldsymbol{z}\beta_j + c_j \bar{\boldsymbol{s}}'_j)$.

We note that the spatial effect of individual archetype may also be of interest. However, the described multivariate regression model may have a high level of colinearity by the definition of archetype vectors. Consider the example where we only have two archetypes. In this case, the two components $\bar{\boldsymbol{s}}'_1, \bar{\boldsymbol{s}}'_2$ satisfies $\bar{\boldsymbol{s}}'_1 + \bar{\boldsymbol{s}}'_2 = 1$ for all cells. This indicates a perfect colinearity, in which case the model will arbitrarily attribute the spatial effect to archetype 1 or archetype 2.

Therefore, we propose another variant of the model. Specifically, we consider fitting $a$ independent models, one for each archetype $j$, with $(\beta_j, c_j)$ as the parameters to fit:

$$\text{Model } j: \ \bar{Y} \sim \bar{\boldsymbol{s}}'_j \boldsymbol{z}\beta_j + c_j \bar{\boldsymbol{s}}'_j. \tag{82}$$

After fitting each individual model, the total spatial effect can be obtained via summing the spatial effect for each archetype, while a global $R^2$ metric can be obtained by taking the maximum intrinsic / spatial variation $R^2$ over archetypes. In practice, we observed that the two variants generate highly similar spatial effects (Supplementary Fig. 11c).

#### 2.3 The underlying assumptions for spatial effect estimation and the positivity index

Estimation of the spatial effect is not always possible for all cells. In this section, we discuss the assumptions of the potential outcome framework [10, 11], under which estimation of this term is feasible.

For a comprehensive introduction to the potential outcome framework, refer to [11]. In our context, the treatment variable corresponds to the multidimensional spatial archetype $\boldsymbol{s}'^i$ for cell $i$. We restate the framework's assumptions in our specific context as follows.

**Assumption 5.** *(SUTVA, Stable Unit Treatment Value Assumption, [10, 11, 12]) The treatment assignment of one cell does not affect the spatial effect of another cell (no interference), and there is no hidden variability in treatment levels.*

**Assumption 6.** *(Ignorability) [10, 11, 12]) The potential outcome is independent of the spatial variation conditioning on the intrinsic variation $Y(\boldsymbol{z}, s') \perp \boldsymbol{s}'|\boldsymbol{z}$ ($s'$ is fixed).*

**Assumption 7.** *(Positivity) [10, 11, 12]) $0 < P(\boldsymbol{s}'_j = 1|\boldsymbol{z}) < 1$ for any $z$ of interest and any index $j$. That is, the treatment should not be deterministic on the intrinsic variation.*

Given the data generation process and the identifiability results outlined in the initial section, all assumptions are automatically satisfied. However, in practical applications, while SUTVA and ignorability still likely hold, the positivity assumption may be violated. For instance, distinct cell types might inhabit non-overlapping microenvironments. In such scenarios, spatial archetypes could be determined by intrinsic variation, thus invalidating the positivity assumption.

To assess the validity of the positivity assumption in real data, we propose positivity indices as proxies for potential assumption violations. We begin by defining a label-based positivity index $P_j$ for each archetype $j$. Our calculation requires a pre-specified label denoted as $c \in \{1, 2, \ldots, l\}$, which can be either cell type annotations or SIMVI intrinsic variation clustering labels. We first select the "pure archetype" cells represented by the set $\{i \mid \max_j \boldsymbol{s}'^i_j > 0.5\}$, and binarize the archetype weights for the subset, yielding a one-hot matrix. Next, we group the cells in this one-hot matrix by their label $c$ and compute label frequencies for each archetype, forming a matrix $\boldsymbol{S}_a \in \mathbb{R}^{l \times a}$. We then take the maximum of each column in this matrix, yielding a vector of shape $\mathbb{R}^a$. The $j$-th component of this vector is defined as $P_j$.

A positivity index $P^i$ thus can be defined for each cell $i$ in either binarized or continuous fashions:

$$P^i = \mathbb{1}_{\exists j,\ s.t.\ P_j > \text{thres1}, \boldsymbol{s}'^i_j > \text{thres2}} \quad \text{(Binarized)} \tag{83}$$

$$P^i = \sum_{j: P_j > \text{thres1}} \boldsymbol{s}'^i_j. \quad \text{(Continuous)} \tag{84}$$

The cell positivity index is a per-cell measure that allows evaluation of the spatial effect feasibility for each cell. A high value indicates potential violation of the positivity assumption.

### 3 Discussions on hyperparameter selection

We performed a comprehensive study on the effects of different parameter settings on SIMVI performance (Supplementary Figs. 1,2,3,4). Specifically, we started from a base model with the following settings (Here n_layers denotes the decoder layer number, the encoder layer number was set to 2, the hidden dim (encoder intermediate layer) was set to 128):

```
model = SimVI(adata, kl_weight=1, kl_gatweight=1, lam_mi=5, permutation_rate=0,
reg_to_use='mi', dis_to_use= 'zinb', n_layers = 1,n_intrinsic=20,n_spatial=10)
```

```
model.train(edge_index,batch_size=1000,mae_epoches=0)
```

We altered one parameter in each experiment, leading to 48 parameter configurations. For each configuration, we performed 5 runs with different random seeds. To ensure the robustness of our findings and avoid dataset-specific parameter tuning, we performed the evaluation on both the MTG and STG datasets, identifying shared parameter settings that enhanced the performance. The same set of random seeds was applied across different configurations. However, we note that settings with non-zero permutation rates and MMD regularizations (which sample from Gaussian distributions) may introduce additional randomness.

We list the meaning of shown parameters and their effects as follows. Important parameters are labeled in bold.

- **Latent dimension size** *dim(z),dim(s)*. The performances of intrinsic variations remain consistent across models with varying dimensionalities. However, the layer preservation performances improve as the dimensionality of spatial variation increases.
- **The batch size** *Batch size*. Larger batch sizes correlate with improved performances of layer preservation. Other metrics remain relatively stable across different batch sizes.
- **The KL divergence weights** *kl*. Notably, reducing the KL weight for spatial variation positively impacts the layer preservation performance.
- **MI / MMD regularization strength** *I(MI),I(MMD)*. Interestingly, increasing *I(MI)* and *I(MMD)* leads to performance differences in some cases but does not yield a clear performance improvement. This is because the regularizations are designed to regress out spatial information from intrinsic variation. However, since cell type dominates the variation in the data, improved disentanglement may not translate into a measurable performance gain for intrinsic variations. This observation is consistent with our main benchmarking results, where several methods that do not explicitly disentangle variations still perform well in cell type preservation and batch removal. The essence of *I(MI)* and *I(MMD)* lies primarily in spatial effect estimation, where removing spatial information from intrinsic variation is essential.
- The number of neighbors *n*. Increasing the number of neighbors enhances global layer preservation but diminishes local niche (MYH11+) preservation performance.
- The number of decoder layers *nl*. No consistent performance differences are observed across experiments.
- **The number of pretraining epochs** *p_epochs* **and the permutation rate** *pr*. Both MTG and STG datasets show slight improvements in layer and cell type preservation performances as *p_epochs* increase. However, no consistent trend is observed regarding different permutation rates, likely due to inherent randomness.
- The loss likelihood setting *NB / ZINB*. The two settings result in similar performance in all cases.

Accounting for the results from the hyperparameter analysis and the default in scVI-based models, we set the default parameter configuration of the SIMVI model as follows:

```
model = SimVI(adata, kl_weight=1, kl_gatweight=0.01, lam_mi=1000, permutation_rate=0.5,
reg_to_use='mmd', dis_to_use= 'zinb', n_layers = 1, n_intrinsic=20,n_spatial=20)
```

The MAE epoch number was set to be 25 and the batch size was set as 500. We observed an increased performance of the default setting combining all SIMVI designs, compared with models with alternative configurations (Supplementary Figs. 1,2,3,4). Our comparison may also be seen as an ablation study showing the contribution of each component in the model. We end the section with two remarks.

*Remark* 8. The selection of optimal parameters may vary with the dataset itself, especially the parameters associated with model training. For example, the batch size may not be set as an default value.

*Remark* 9. Overall, we observed consistent performances of SIMVI across independent runs with the same configuration (Supplementary Figs. 1,2). This supports our theoretical identifiability results and is further validated across different datasets.

#### References

[1] Gayoso, A. *et al.* A python library for probabilistic analysis of single-cell omics data. *Nature biotechnology* **40**, 163–166 (2022).

[2] Khemakhem, I., Kingma, D., Monti, R. & Hyvarinen, A. Variational autoencoders and nonlinear ica: A unifying framework 2207–2217 (2020).

[3] Comtet, L. *Advanced Combinatorics: The art of finite and infinite expansions* (Springer Science & Business Media, 2012).

[4] Lin, G. D. Recent developments on the moment problem. *Journal of Statistical Distributions and Applications* **4**, 5 (2017).

[5] Petersen, L. On the relation between the multidimensional moment problem and the one-dimensional moment problem. *Mathematica Scandinavica* 361–366 (1982).

[6] Kleiber, C. & Stoyanov, J. Multivariate distributions and the moment problem. *Journal of Multivariate Analysis* **113**, 7–18 (2013).

[7] Sard, A. The measure of the critical values of differentiable maps (1942).

[8] Mørup, M. & Hansen, L. K. Archetypal analysis for machine learning and data mining. *Neurocomputing* **80**, 54–63 (2012).

[9] Chernozhukov, V. *et al.* Double/debiased machine learning for treatment and structural parameters (2018).

[10] Imbens, G. W. & Rubin, D. B. *Causal inference in statistics, social, and biomedical sciences* (Cambridge university press, 2015).

[11] Yao, L. *et al.* A survey on causal inference. *ACM Transactions on Knowledge Discovery from Data (TKDD)* **15**, 1–46 (2021).

[12] Vegetabile, B. G. On the distinction between" conditional average treatment effects"(cate) and" individual treatment effects"(ite) under ignorability assumptions. *arXiv preprint arXiv:2108.04939* (2021).

### Supplementary Figures

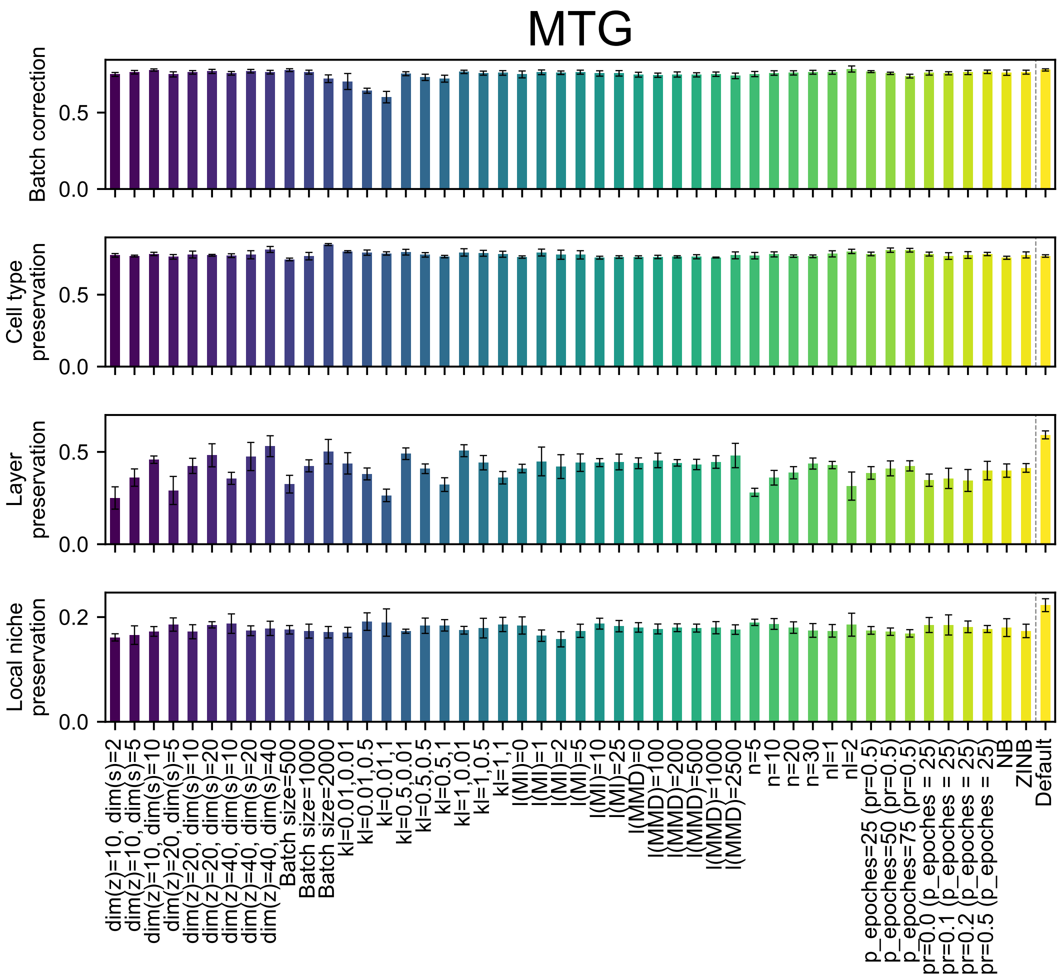


Supplementary Figure 1: Boxplot showing metric scores for different SIMVI parameter configurations on the MERFISH MTG dataset. The bar heights represent the average performance across 5 different random seeds ($n = 10$ for Default). Error bars indicate standard deviations.

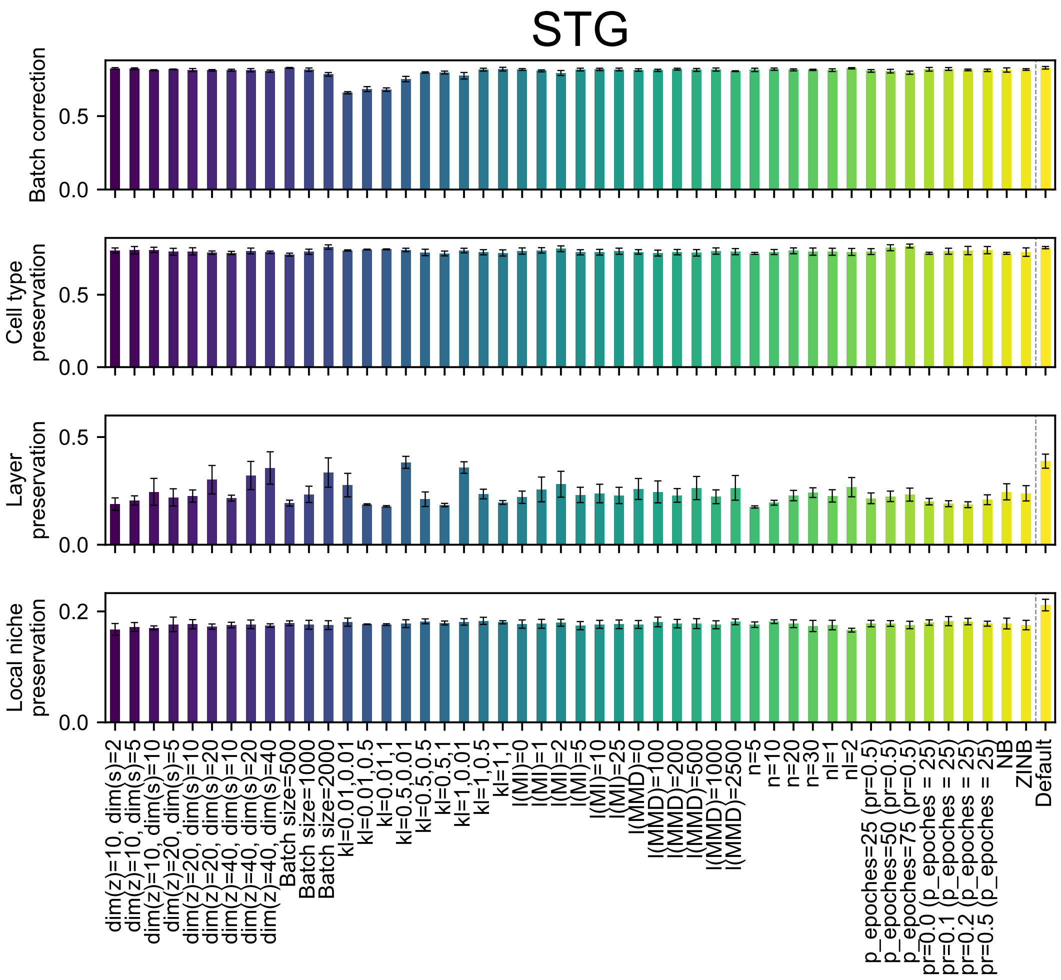


Supplementary Figure 2: Boxplot showing metric scores for different SIMVI parameter configurations on the MERFISH STG dataset. The bar heights represent the average performance across 5 different random seeds ($n = 10$ for Default). Error bars indicate standard deviations.

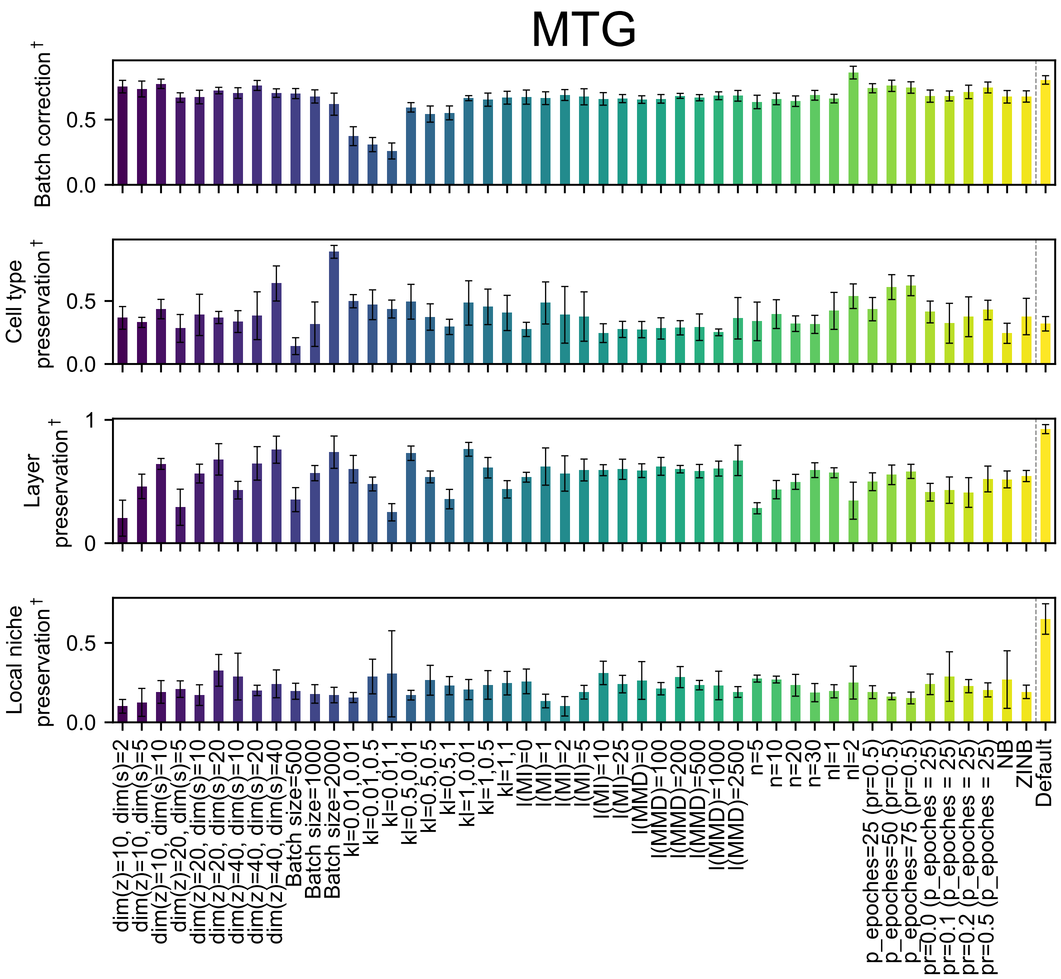


Supplementary Figure 3: Boxplot showing scaled metric scores (`min_max_scale = True` in scib-metrics) for different SIMVI parameter configurations on the MERFISH MTG dataset. The bar heights represent the average performance across 5 different random seeds ($n = 10$ for Default). Error bars indicate standard deviation.

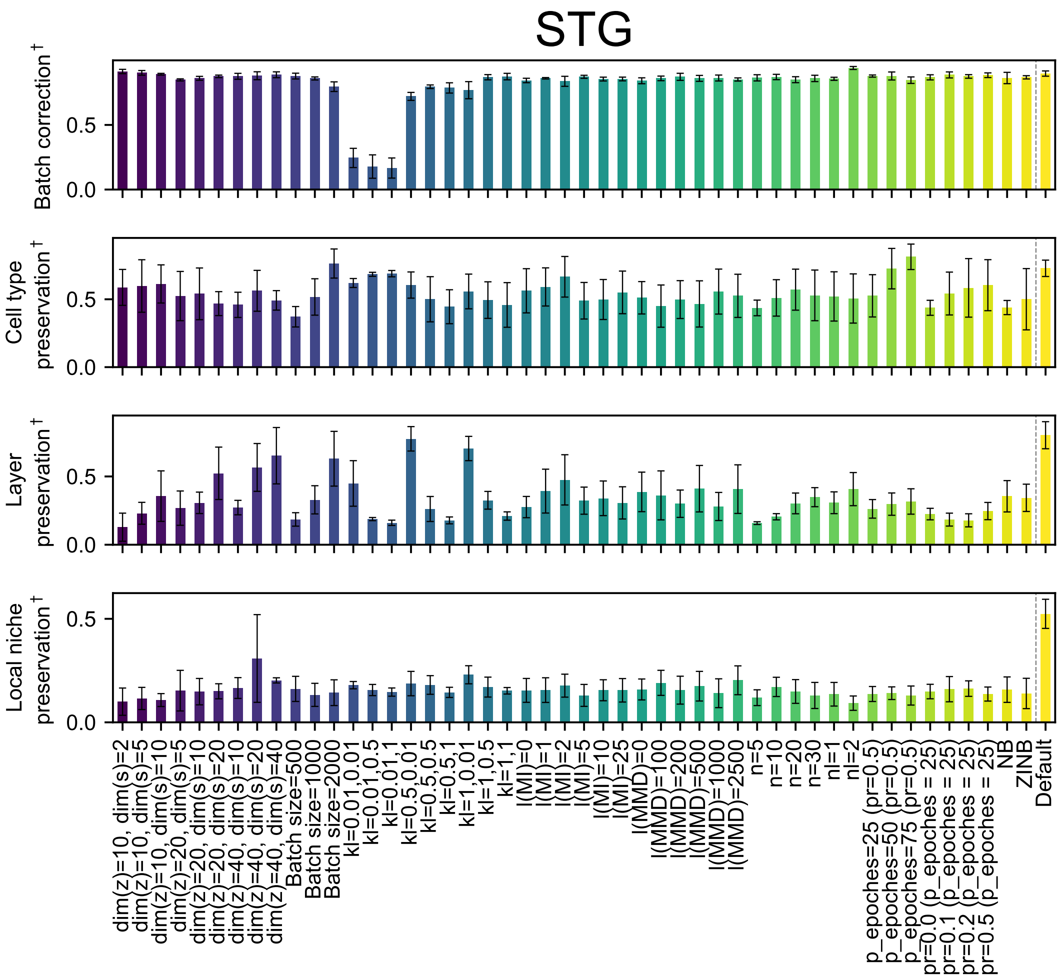


Supplementary Figure 4: Boxplot showing scaled metric scores (`min_max_scale = True` in scib-metrics) for various SIMVI parameter configurations on the MERFISH STG dataset. The bar heights represent the average performance across 5 different random seeds ($n = 10$ for Default). Error bars indicate standard deviation.

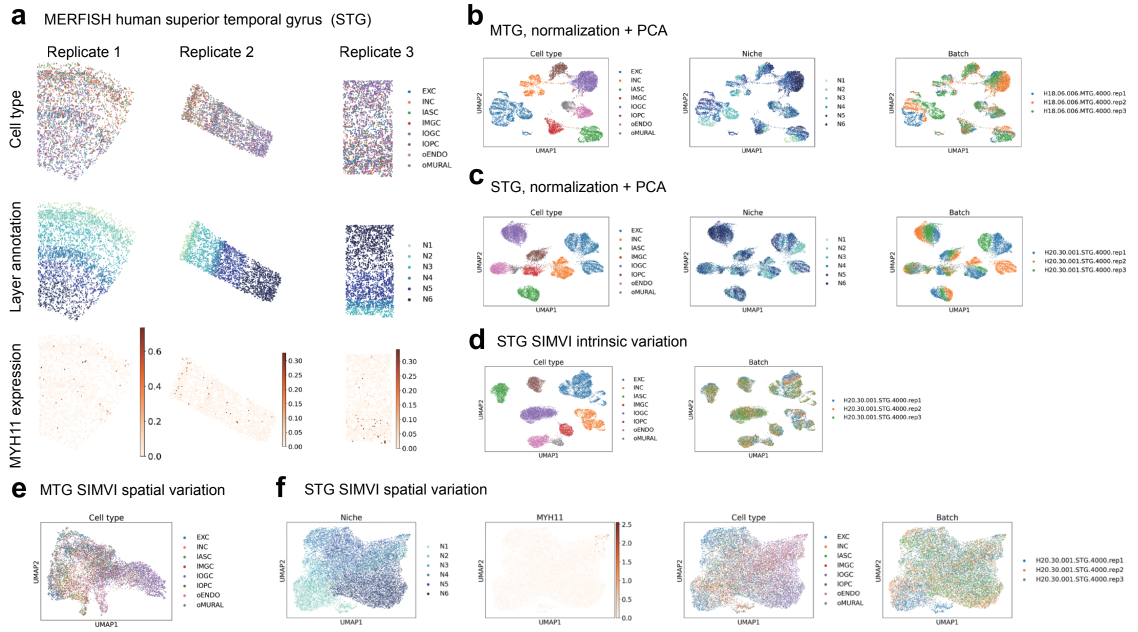


Supplementary Figure 5: Additional visualizations on MERFISH MTG / STG datasets. **a.** Overview of the MERFISH STG data, showing spatial organizations of cell type, layer annotation, and log normalized *MYH11* expression for STG replicates 1-3. **b.** UMAP visualization of the MERFISH MTG dataset using the principal components derived from log normalized gene expression, colored by cell type, layer annotation, and batch label. **c.** UMAP visualization of the MERFISH STG dataset using the principal components derived from log normalized gene expression, colored by cell type, layer annotation, and batch label. **d.** UMAP visualization of the SIMVI intrinsic variation on MERFISH STG dataset, colored by cell type and batch label. **e.** UMAP visualization of the SIMVI spatial variation on MERFISH MTG dataset, colored by cell type. **f.** UMAP visualization of the SIMVI spatial variation on MERFISH STG dataset, colored by layer annotation, log normalized *MYH11* expression, cell type, and batch label.

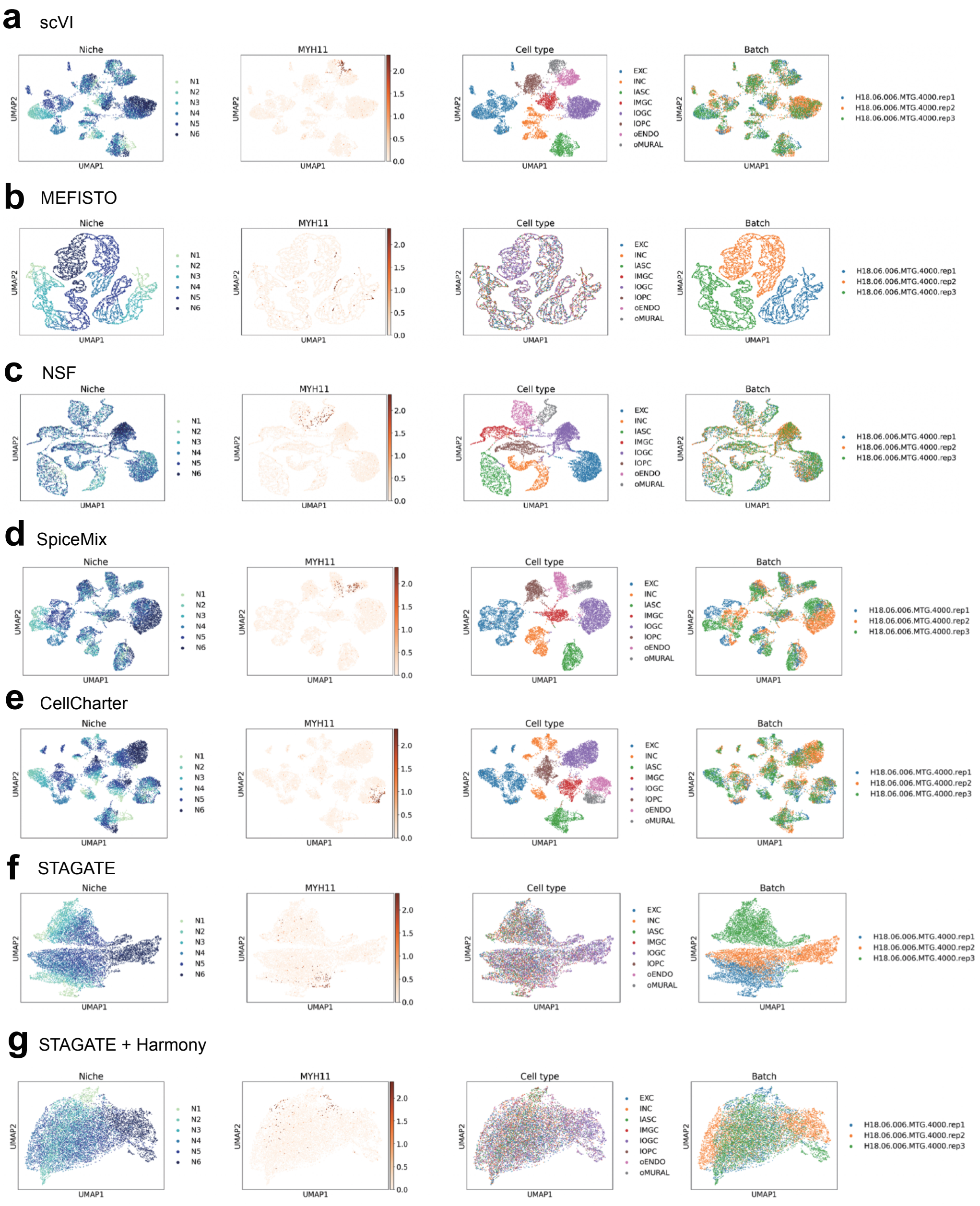


Supplementary Figure 6: UMAP visualization of scVI (**a**), MEFISTO (**b**), NSF (**c**), SpiceMix (**d**), CellCharter (**e**), STAGATE (**f**), STAGATE + Harmony (**g**) for the MTG dataset, colored by layer annotation, log normalized *MYH11* expression, cell type, and batch label.

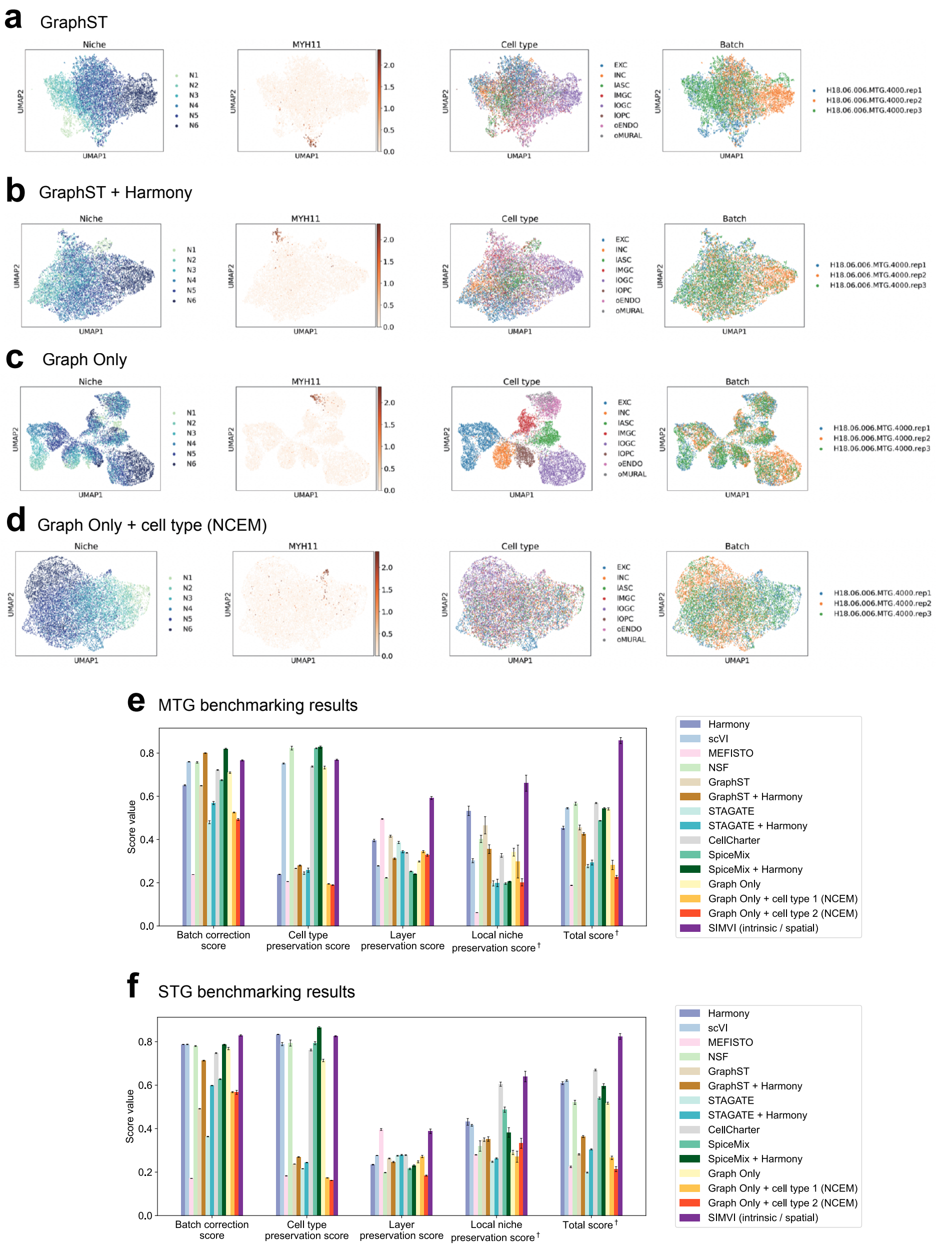


Supplementary Figure 7: Continued UMAP visualization of GraphST (**a**), GraphST + harmony (**b**), Graph Only (**c**), Graph Only + cell type (**d**) for the MTG dataset, colored by layer annotation, log normalized *MYH11* expression, cell type, and batch label. **e.** Bar plots showing metric scores for the MERFISH MTG dataset. The bar heights represent the average performance across 10 different random seeds, with the error bars showing standard errors. The only difference from the results shown in Main Fig. 2 is the calculation of the total score (last column). **f.** Bar plots showing metric scores for the MERFISH STG dataset. The only difference from the results shown in Main Fig. 2 is the calculation of the total score (last column). The bar heights represent the average performance across 10 different random seeds, with the error bars showing standard errors. †: Each individual metric from all experiments is scaled to the range [0,1], then the final score is calculated as the average of these rescaled values.

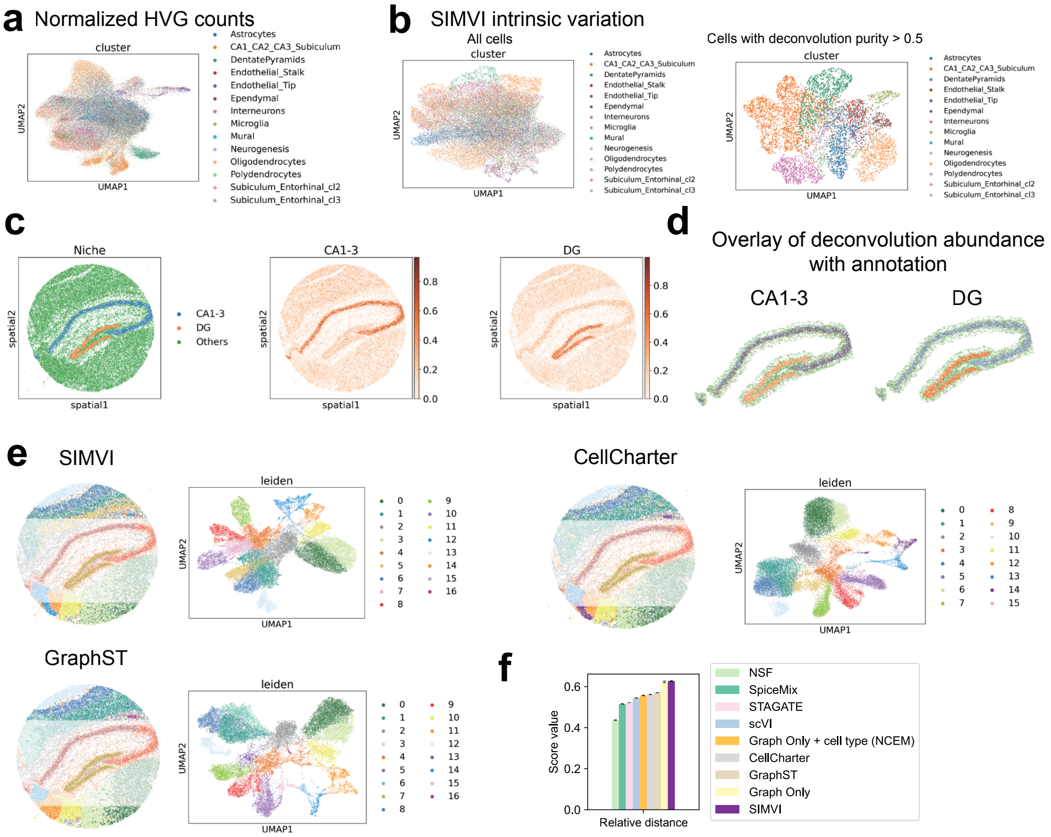


Supplementary Figure 8: Additional analysis results on Slide-seqv2 mouse hippocampus data. **a**. UMAP visualization using the principal components derived from log normalized highly variable gene expression, colored by dominant cell type label. **b**. UMAP visualization for the full data (left) and the cells with deconvolution purity > 0.5 (right) using the SIMVI intrinsic variation, colored by dominant cell type label. **c**. Spatial visualization of the niche annotation, and deconvolution ratio of CA1-3 and DG cells from prior annotation. **d**. Spatial visualization overlaying CA1-3 (left) and DG (right) deconvolution ratio and niche annotation. **e**. Spatial visualization overlaying niche annotation (left), and UMAP visualization colored by Leiden clusters for SIMVI, CellCharter and GraphST. **f**. Benchmarking results of different methods in terms of relative distances between CA, DG, and their neighborhoods. Relative distances are calculated using Silhouette width.

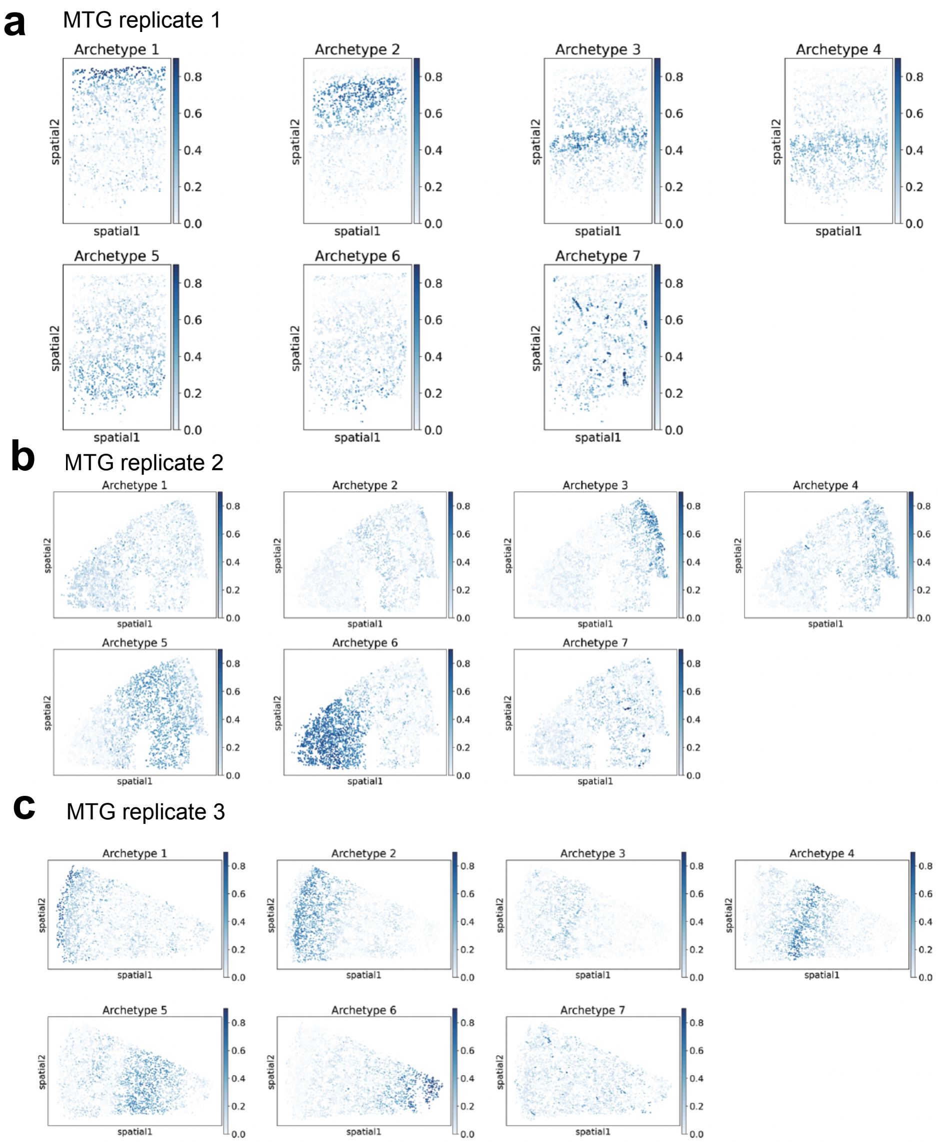


Supplementary Figure 9: Spatial visualization of archetype components in MERFISH MTG replicate 1 (**a**), 2 (**b**), 3 (**c**).

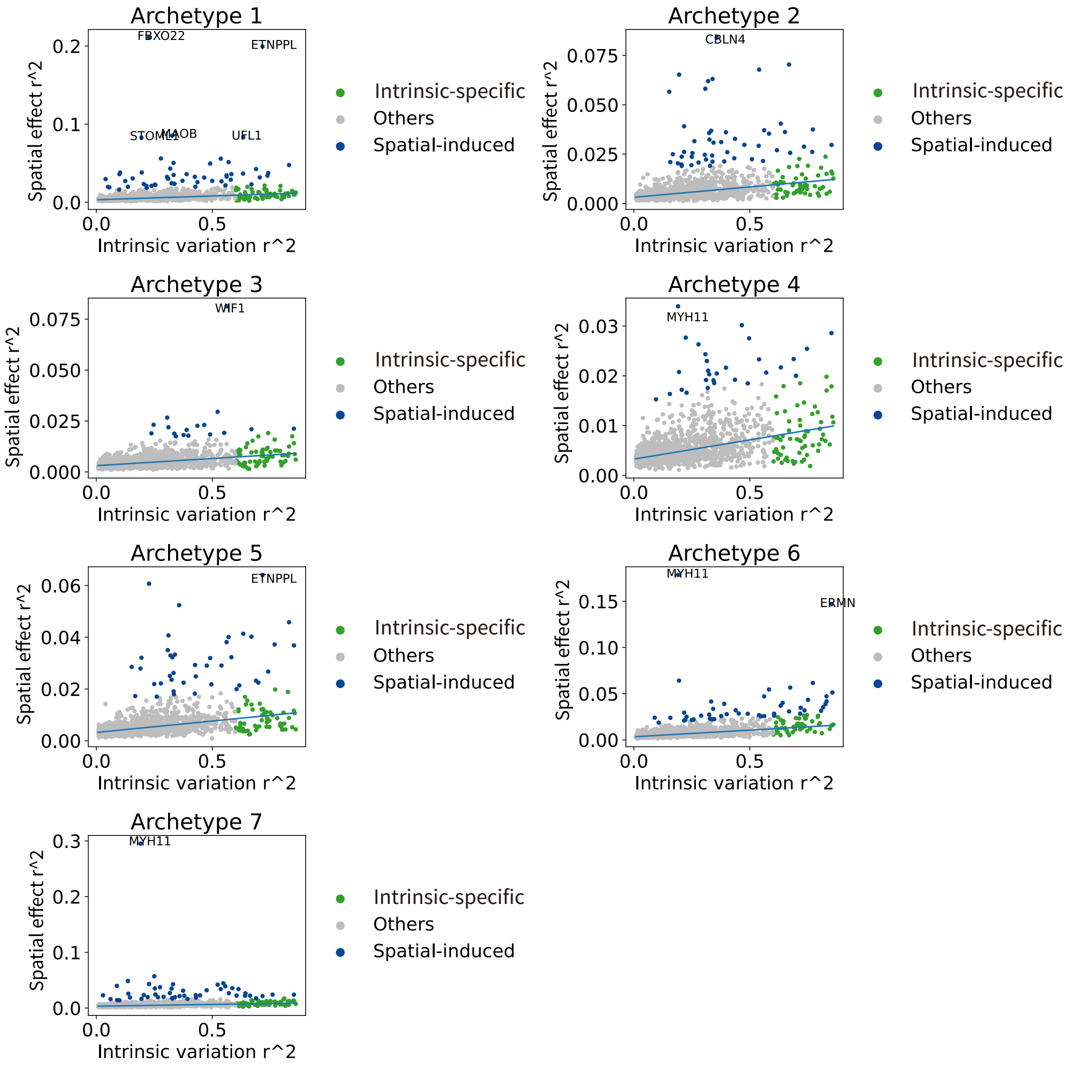


Supplementary Figure 10: $R^2$ scatter plot of each individual archetype for the MERFISH MTG dataset. Genes with scaled Huber regression residual larger than 10 were annotated as spatial-induced. Other genes with intrinsic-specific $R^2$ larger than 0.6 were annotated as intrinsic-specific.

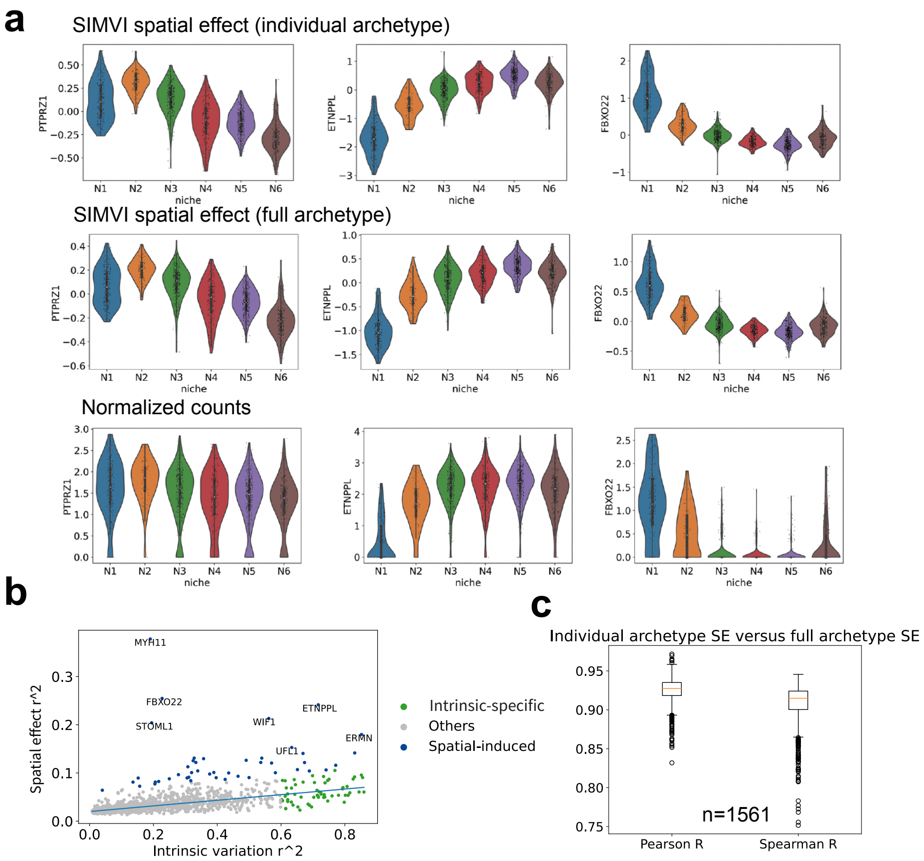


Supplementary Figure 11: Additional visualizations of the SIMVI spatial effect applied in MERFISH MTG data. **a**. Violin plot of SIMVI spatial effect (individual archetype), SIMVI spatial effect (full archetype), and normalized counts for example genes. **b**. $R^2$ decomposition scatter plot using the SIMVI SE full archetype mode. Genes with scaled Huber regression residual larger than 10 were annotated as spatial-induced. Other genes with intrinsic-specific $R^2$ larger than 0.6 were annotated as intrinsic-specific. **c**. Boxplot of Pearson and Spearman correlations across SIMVI SE (individual archetype mode) and SIMVI SE (full archetype mode).

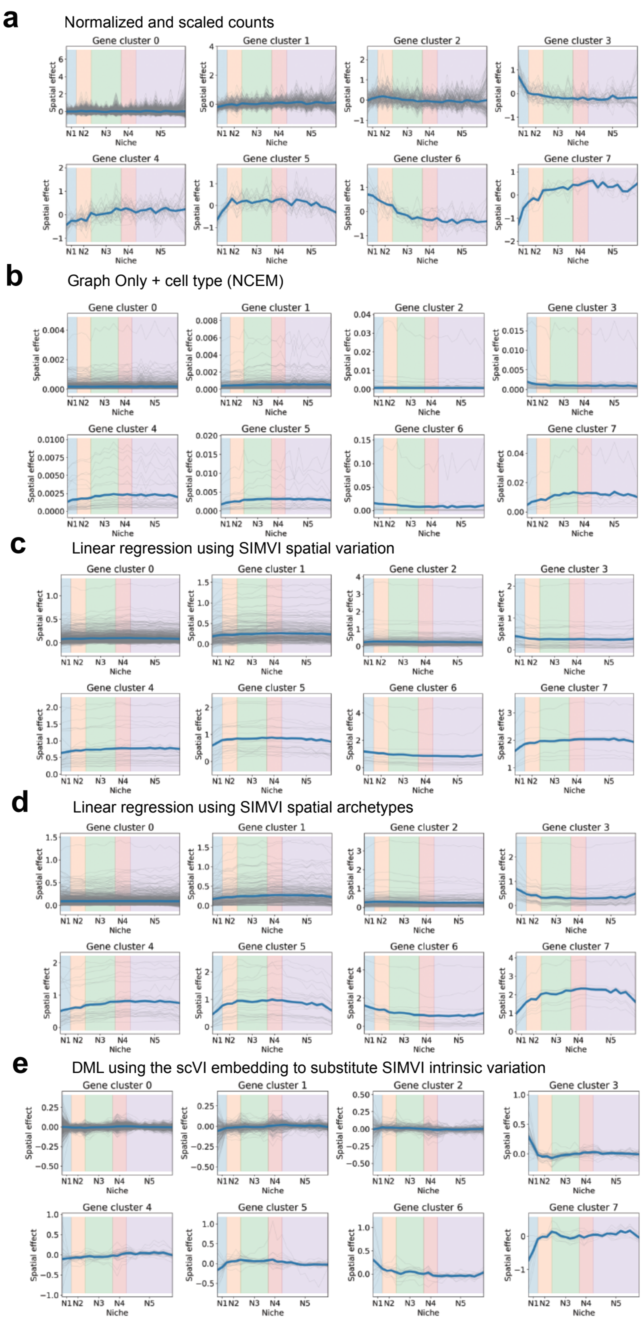


Supplementary Figure 12: Comparison of alternative methods in revealing gene expression spatial patterns in MERFISH MTG data replicate 1. **a**. Log normalized and scaled counts. **b**. Binned normalized expression returned by graph only + cell type (NCEM) model. **c**. Linear regression predictions using SIMVI spatial variation. **d**. Linear regression predictions using SIMVI spatial archetypes. **e**. Binned linear regression predictions using SIMVI spatial archetypes. Data from all panels were binned as described in the Methods section.

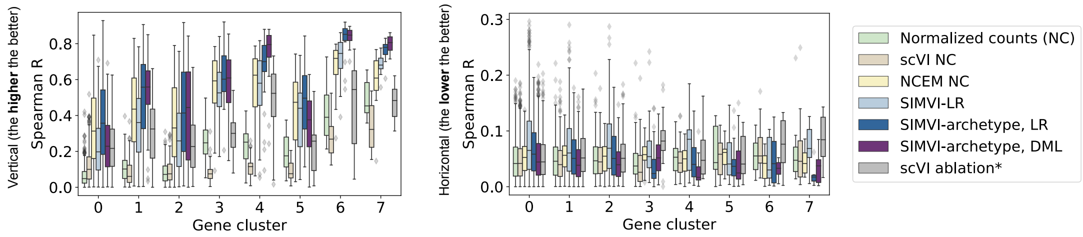


Supplementary Figure 13: Boxplot showing Spearman R on each individual gene of astrocytes from MTG replicate 1 and the spatial vertical coordinate (left) and horizontal coordinate (right). *: scVI ablation refers to the DML approach that fixes SIMVI spatial variation and replaces SIMVI intrinsic variation with the scVI embedding. The gene number (n) is 1002, 265, 176, 40, 31, 23, 16, 8 for each cluster. The top/lower hinge represents the upper/lower quartile and whiskers extend to the largest/smallest value within 1.5× interquartile range from the hinge. The median is shown as the center.

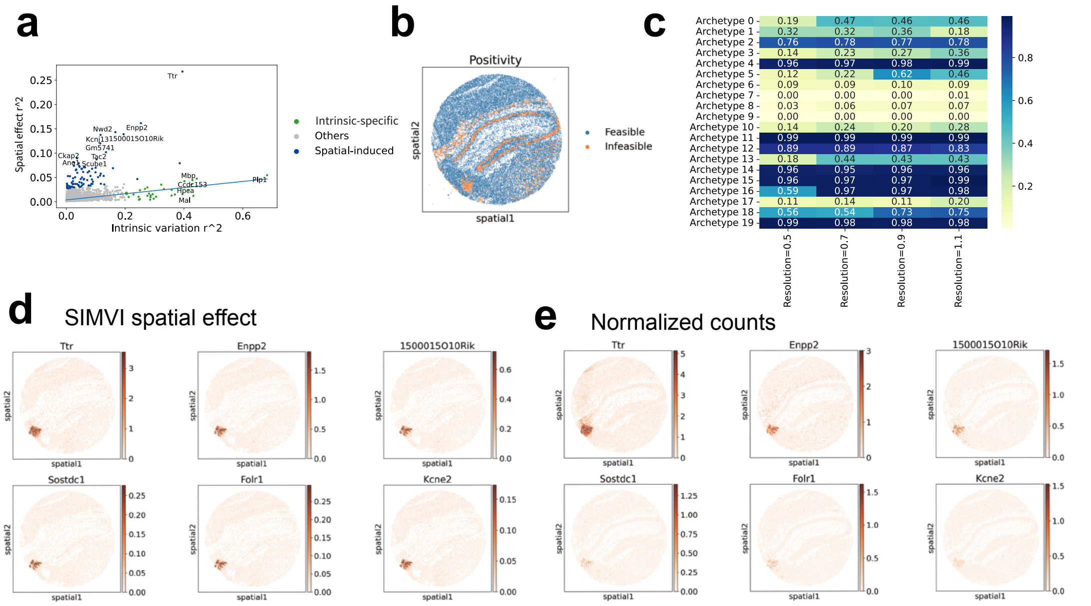


Supplementary Figure 14: Additional visualizations of SIMVI spatial effect for Slide-seqv2 data. **a**. Scatter plot showing the intrinsic variation $R^2$ and spatial effect $R^2$ for each individual gene in Slide-seqV2 mouse hippocampus. Genes with scaled Huber regression residual larger than 100 were annotated as spatial-induced. Other genes with intrinsic-specific $R^2$ larger than 0.2 were annotated as intrinsic-specific. **b**. Spatial visualization of pixels satisfying the positivity condition in Slide-seqV2 mouse hippocampus. **c**. Table showing the effect of Leiden clustering resolution on the archetype positivity index. Higher values indicate potential violation of the positivity condition. **d**. Spatial visualizations of representative genes' spatial effects. **e**. Spatial visualizations of representative genes' log normalized expression in Slide-seqV2 mouse hippocampus.

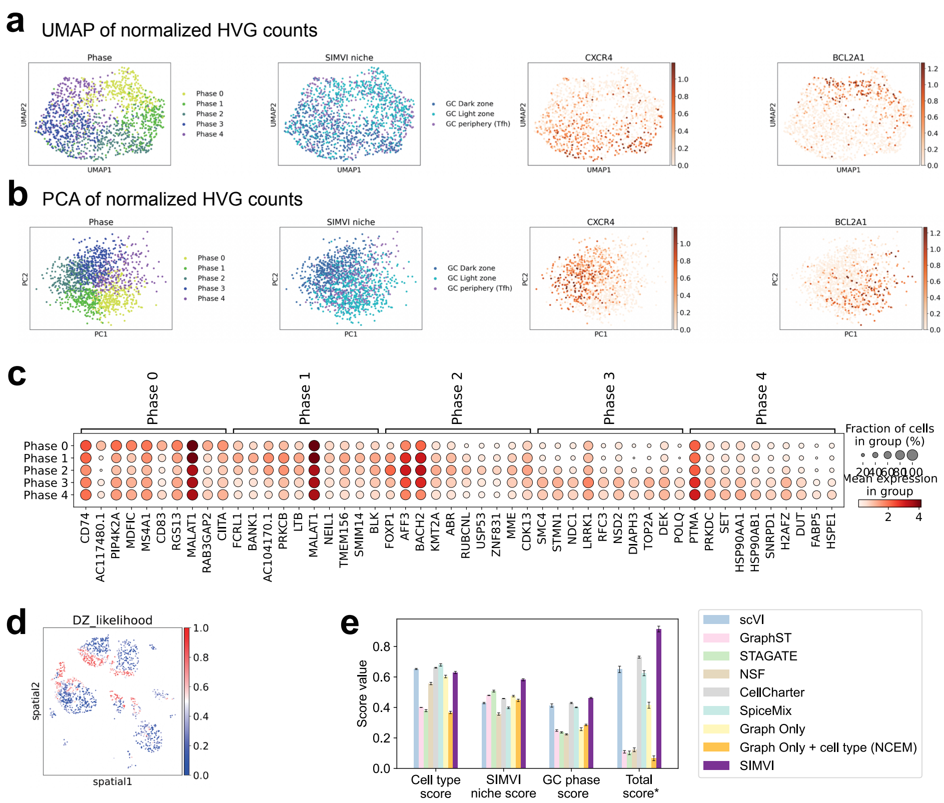


Supplementary Figure 15: Additional analysis results for the Slide-tags human tonsil data. **a**. UMAP visualization of the original data (log normalized highly variable gene expression), colored by annotated phase, niche, *CXCL4* and *BCL2A1* expression. **b**. Visualization of the first two principal components of the original data (log normalized highly variable gene expression), colored by annotated phase, niche, log normalized *CXCL4* and *BCL2A1* expression. **c**. Dotplot showing the marker gene expression across phases. **d**. Spatial visualization of GC B cells colored by dark zone likelihood. **e**. Bar plots showing metric scores for the Slide-tags human tonsil dataset. The bar heights represent the average performance across 10 different random seeds, with the error bars showing standard errors. *: The score is obtained by first averaging the individual score values for each experiment, then rescaling the average score across all experiments to the range [0,1].

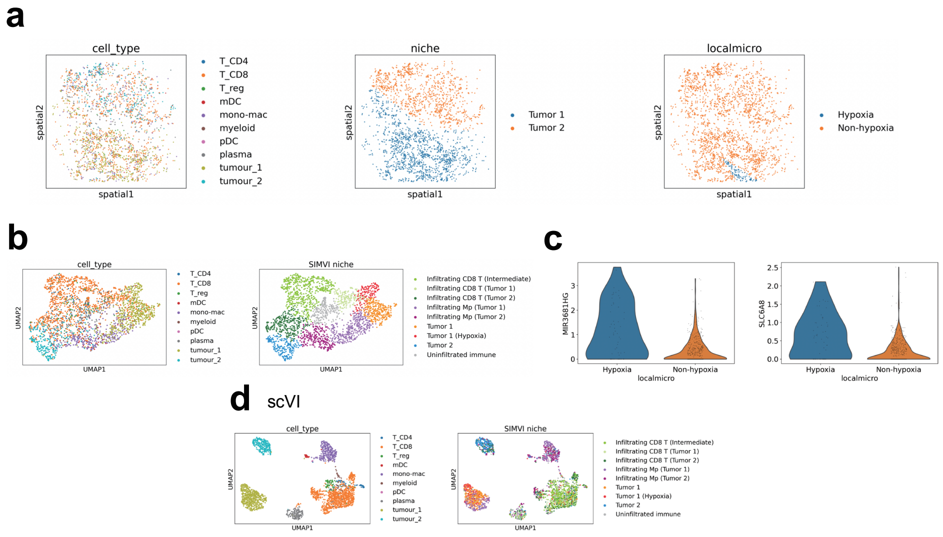


Supplementary Figure 16: Additional visualization of SIMVI analysis on spatial multiome melanoma data. **a**. Spatial visualization colored by cell type, annotated tumor niche, and the annotated local hypoxia microenvironment. **b**. UMAP visualization of the SIMVI spatial variation, colored by cell type and annotated SIMVI niche. **c**. Violin plot showing gene expression in the "hypoxia" microenvironment versus other cells of tumor 1. **d**. UMAP visualization of the scVI representation, colored by cell type and annotated SIMVI niche.

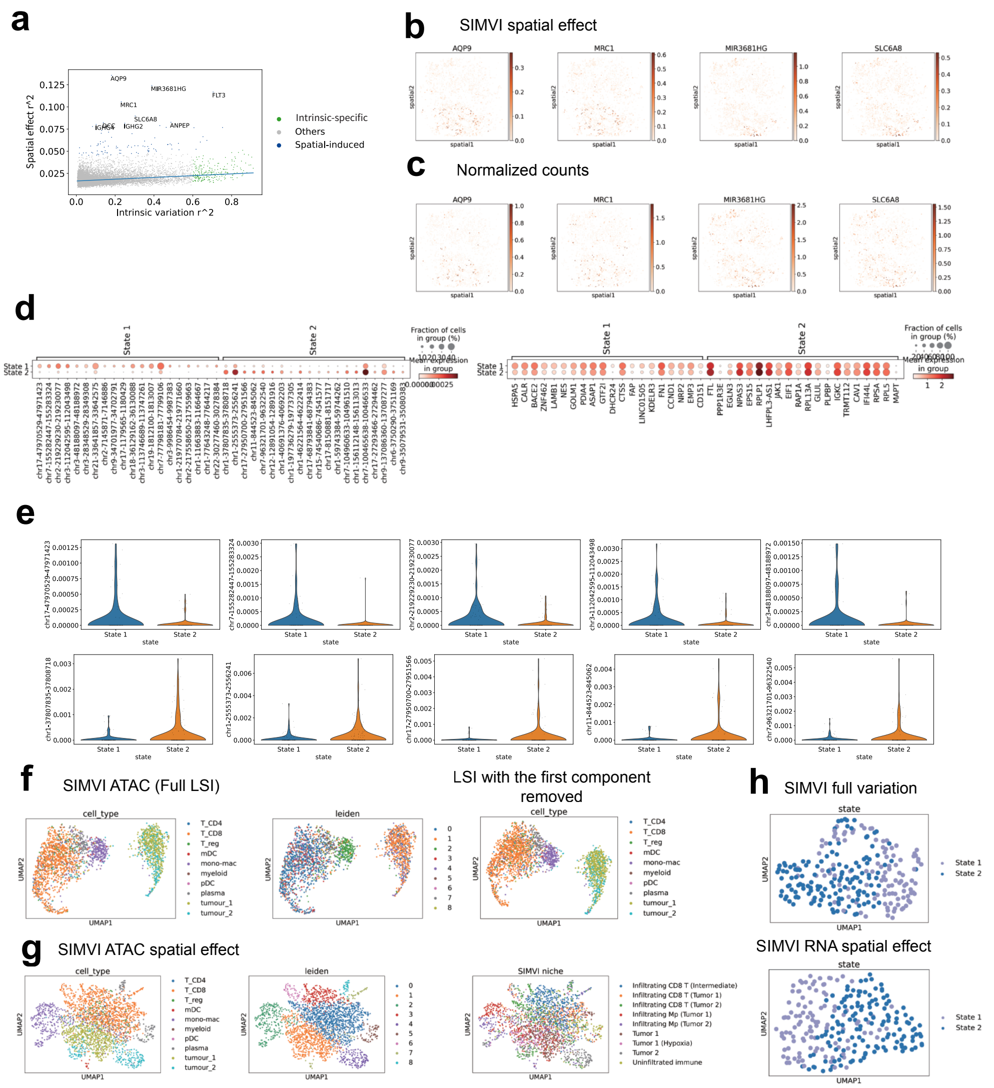


Supplementary Figure 17: Additional results of spatial effect analysis for the spatial multiome melanoma data. **a**. $R^2$ scatter plot of SIMVI spatial effect. Genes with scaled Huber regression residual larger than 10 were annotated as spatial-induced. Other genes with intrinsic-specific $R^2$ larger than 0.6 were annotated as intrinsic-specific. **b**. Spatial visualization of the SIMVI spatial effect of example genes, colored by log normalized expression. **c**. Spatial visualization of the log normalized expression of example genes. **d**. Dotplot of the tumor 2 state representative genes from ATAC (left) and RNA (right) modalities. **e**. Violin plot of representative differential ATAC peaks across tumor 2 states. **f**. UMAP visualization of the LSI embedding of the ATAC modality, colored by cell type and leiden clustering label from SIMVI spatial effect. **g**. UMAP visualization of the SIMVI ATAC spatial effect, colored by cell type label, leiden clustering, and SIMVI niche label. **h**. UMAP visualization of the SIMVI full variation / spatial effect of the tumor 2 state 1 / 2, colored by annotated states.

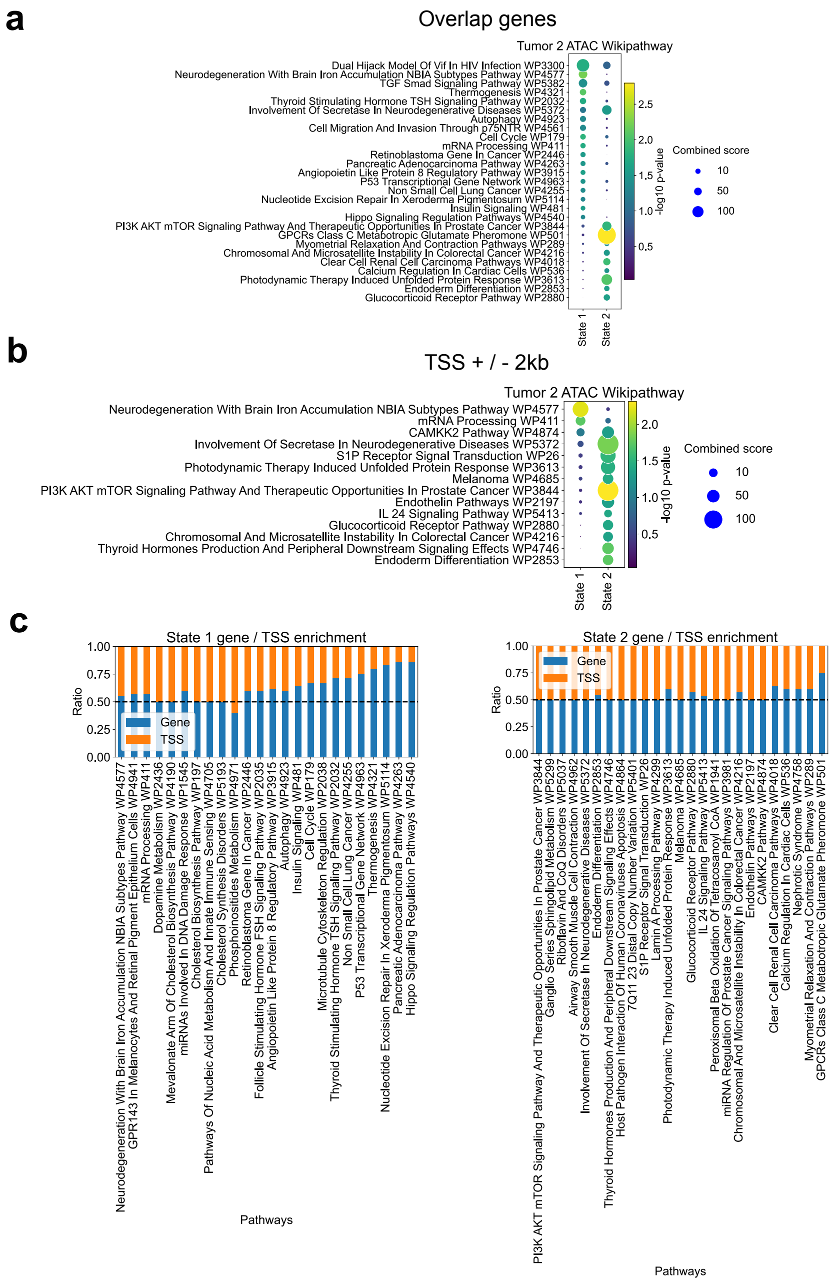


Supplementary Figure 18: Gene set enrichment analyses for ATAC peaks across tumor states. **a**. Dotplot showing selected EnrichR Wikipathway corresponding to genes overlapping with differential ATAC peaks across the two states of tumor 2. **b**. Dotplot showing selected EnrichR Wikipathway corresponding to TSS + / - 2kb overlapping with differential ATAC peaks across the two states of tumor 2. **c**. Comparison of the enriched gene / TSS ratio across significantly enriched Wikipathways for state 1 / 2.

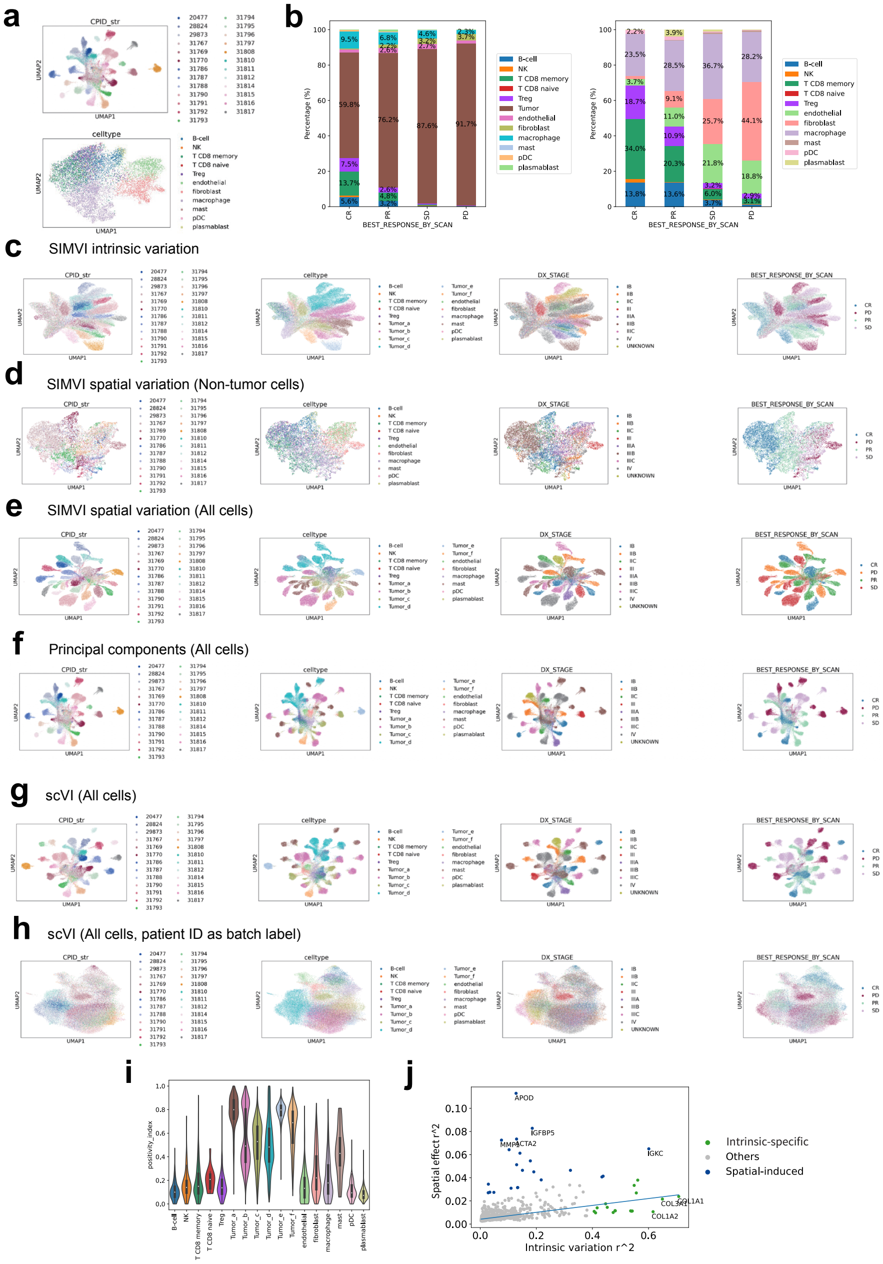


Supplementary Figure 19: Additional results for CosMx melanoma data. **a**. UMAP visualization of log normalized counts for all cells (upper) and non-tumor cells (lower), colored by patient ID and cell type respectively. **b**. Stacked bar plot showing total cell type fractions (left) and non-tumor cell type fractions (right) across patient phenotypes. **c-h**. UMAP visualization of embeddings generated by different methods, colored by patient ID, cell type, tumor stage and patient outcome. **i**. Violin plot showing the summary positivity index across different cell types. **j**. $R^2$ scatter plot for non-tumor cell spatial effect in the CosMx melanoma dataset. Genes with scaled Huber regression residual larger than 20 were annotated as spatial-induced. Other genes with intrinsic-specific $R^2$ larger than 0.4 were annotated as intrinsic-specific.

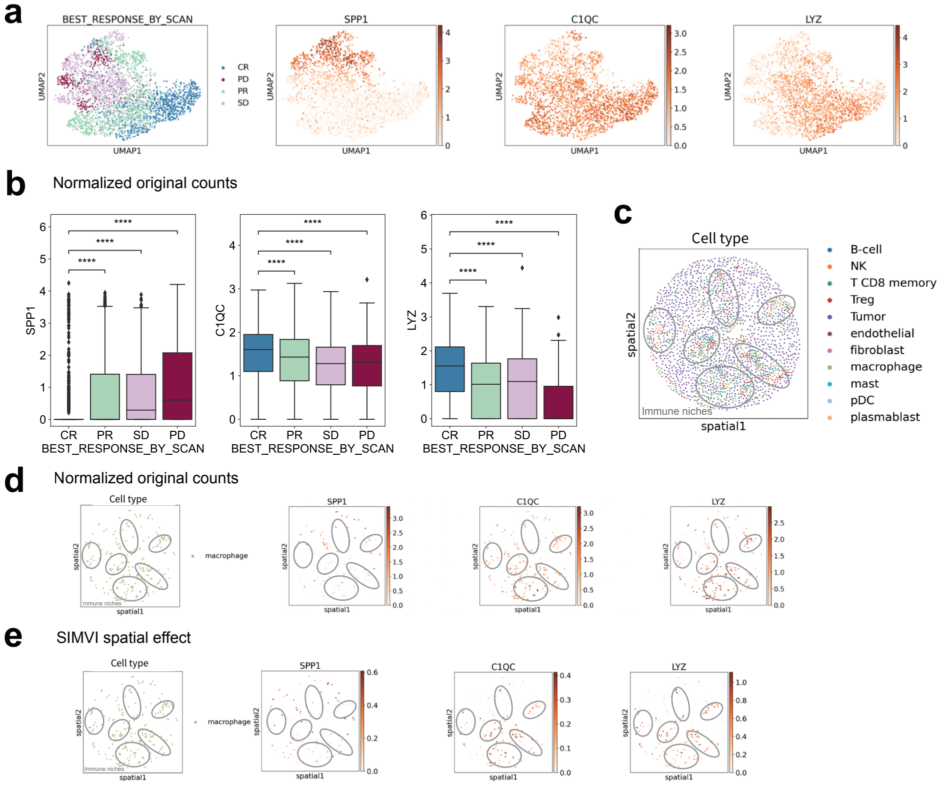


Supplementary Figure 20: Additional results on SIMVI spatial effect for macrophages. **a**. UMAP visualization of macrophages by log normalized gene expression, colored by patient outcome and signature gene expression. **b**. Boxplot showing log normalized expression of signature genes across patient outcomes. Mann-Whitney tests were performed. ****: p-value $< 1 \times 10^{-4}$. **c**. Spatial visualization of an patient example (31788) colored by cell types. Immune cell niches were circled in gray. **d**. Spatial visualization of log normalized signature gene expression in macrophages. Immune cell niches were circled in gray. **e**. Spatial visualization of SIMVI spatial effect for signature genes in macrophages. Immune cell niches were circled in gray.

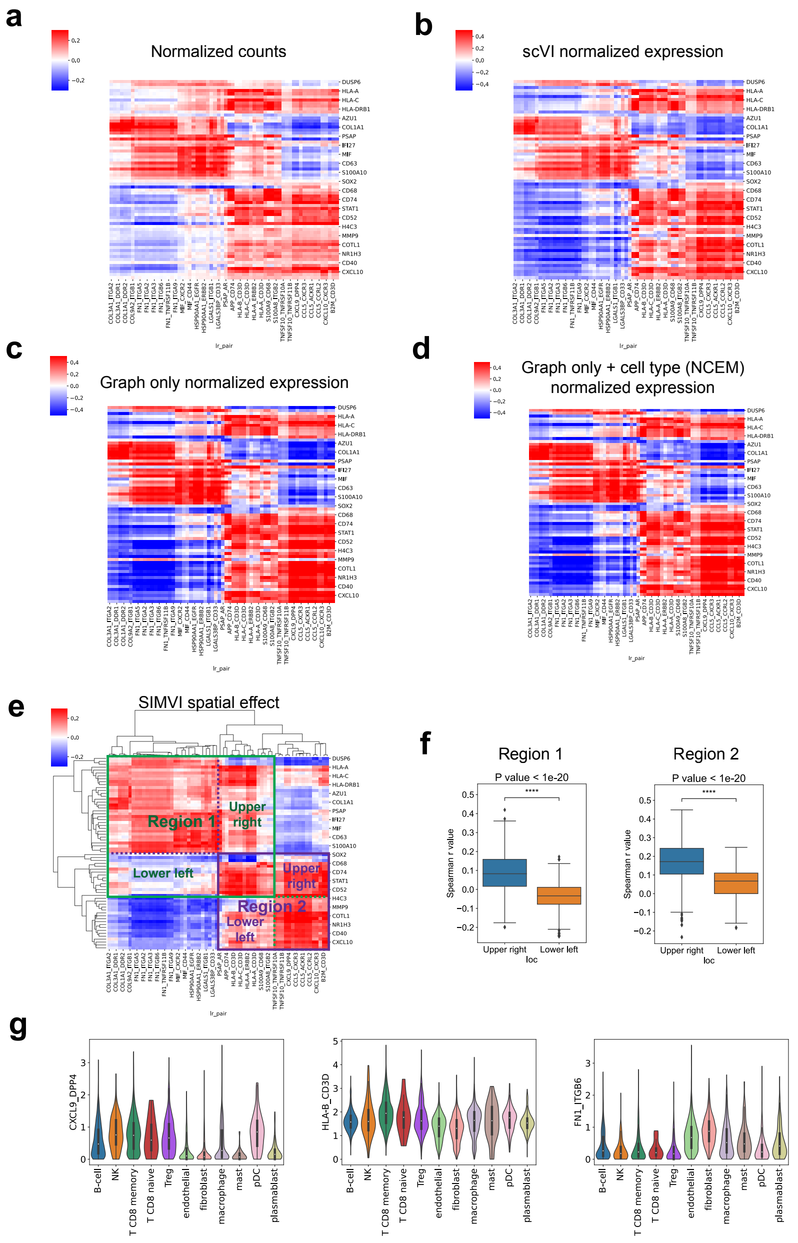


Supplementary Figure 21: Spearman correlation map between ligand-receptor expression level and normalized counts (**a**), scVI normalized expression (**b**), Graph only normalized expression (**c**), Graph only + cell type (NCEM) normalized expression (**d**), SIMVI spatial effect (**e**). The row and column orders of the correlation maps are fixed to match the result of hierarchical clustering on the SIMVI spatial effect cluster map. **f**. Boxplot comparing entries of the SIMVI Spearman correlation map in upper right and lower left locations of region 1 and 2. **g**. Violin plot of example ligand-receptor expression levels across cell types.

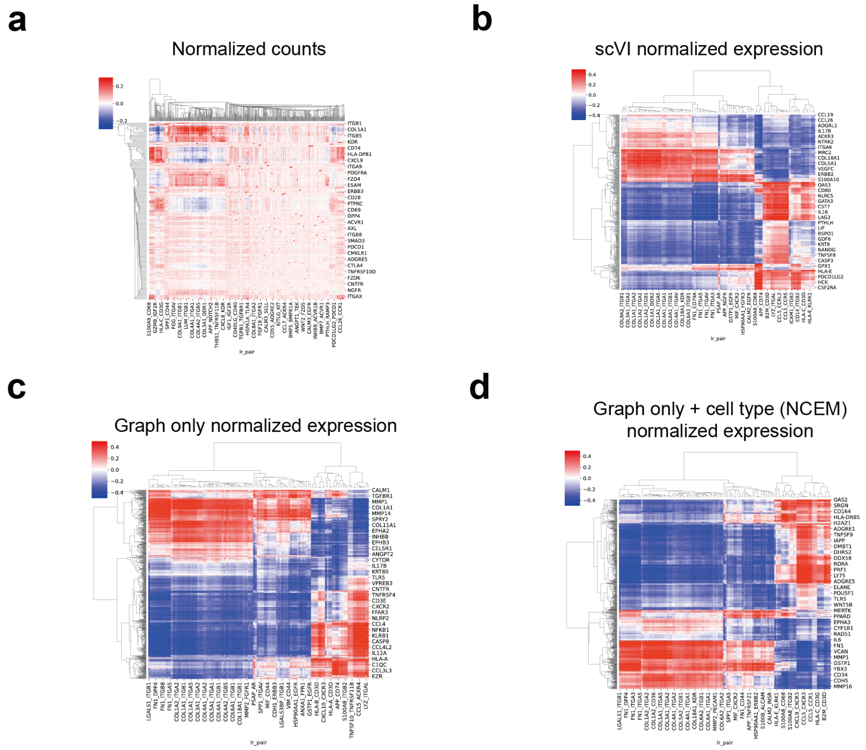


Supplementary Figure 22: Hierarchical clustered Spearman correlation maps between ligand-receptor expression level and normalized original counts (**a**), scVI normalized expression (**b**), Graph only normalized expression (**c**), Graph only + cell type (NCEM) normalized expression (**d**). The rows and columns with max absolute values larger than 0.4 are shown.
